# Macromolecular and cytological changes in fission yeast G0 nuclei

**DOI:** 10.1101/2020.01.11.899559

**Authors:** Zhi Yang Tan, Shujun Cai, Saayli A. Paithankar, Xin Nie, Jian Shi, Lu Gan

## Abstract

When starved of nitrogen, fission yeast *Schizosaccharomyces pombe* cells enter a quiescent “G0” state with smaller nuclei and transcriptional repression. The genomics of *S. pombe* G0 cells has been well studied, but much of its nuclear cell biology remains unknown. Here we use confocal microscopy, immunoblots, and electron cryotomography to investigate the cytological, biochemical, and ultrastructural differences between *S. pombe* proliferating, G1-arrested, and G0 cell nuclei, with an emphasis on the histone acetylation, RNA polymerase II fates, and macromolecular complex packing. Compared to proliferating cells, G0 cells have lower levels of histone acetylation, nuclear RNA polymerase II, and active transcription. The G0 nucleus has similar macromolecular crowding yet fewer chromatin-associated multi-megadalton globular complexes. Induced histone hyperacetylation in G0 results in cells that have larger nuclei and therefore less compact chromatin. However, these histone-hyperacetylated G0 cells remain transcriptionally repressed with similar nuclear crowding. Canonical nucleosomes – those that resemble the crystal structure – are rare in proliferating, G1-arrested, and G0 cells. Our study therefore shows that extreme changes in nucleus physiology are possible without extreme reorganisation at the macromolecular level.

**Summary:** We use multiple cell-biological techniques to compare proliferating and quiescent G0 fission yeast cell nucleus transcription, histone acetylation, macromolecular packing.

## INTRODUCTION

Eukaryotic cells can adopt states that vary greatly in nuclear morphology and chromatin biochemistry and structure. Coordinated phenotypic changes in the nucleus are believed to regulate essential functions like transcription, replication, DNA repair, and stress response. The fission yeast *Schizosaccharomyces pombe* is an excellent model organism for nucleus cell biology because the cells can be enriched in different cell-cycle states, each with distinctive changes in chromosome compaction (Hiraoka et al., 1984; Nurse et al., 1976). When starved of nitrogen for 24 hours, *S. pombe* cells enter a non-dividing state called G0 quiescence (Su et al., 1996) (herein called G0) (**Figure 1A**). G0 cells are shorter and more thermotolerant than proliferating cells (Su et al., 1996). Furthermore, G0 cells have only ∼30% of the mRNA and ∼20% of the rRNA content of proliferating cells (Marguerat et al., 2012), suggesting that transcription is globally repressed.

**Figure 1.**
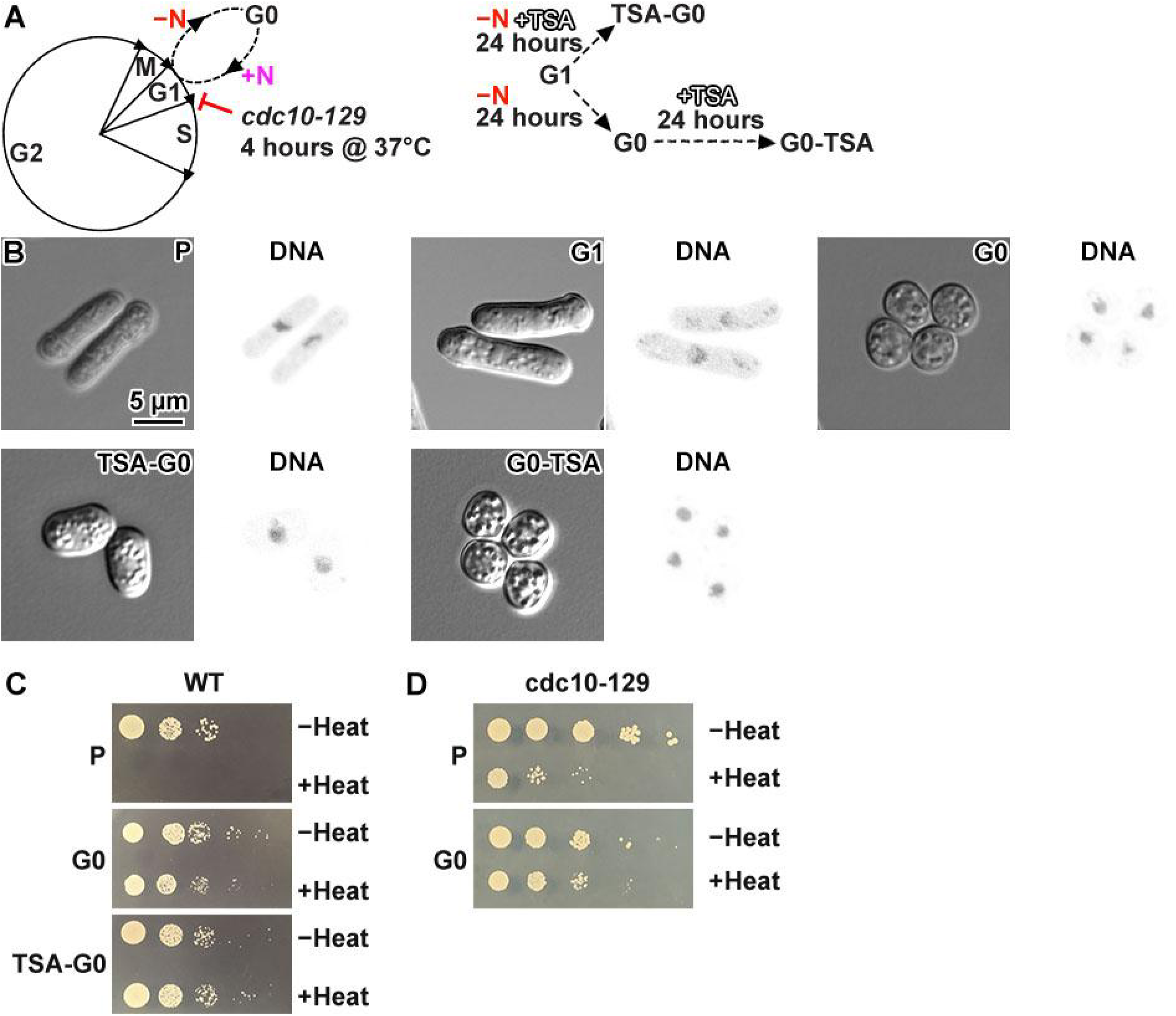
Overview of *S. pombe* cells in proliferative and G0 states. (A) Left: Schematic of the *S. pombe* cell cycle: G1 phase, S phase, G2 phase, and Mitosis (M). Entry to and exit from G0 is shown with the dashed line and depends on the availability of nitrogen (−N / +N). The *cdc10-129* strain arrests in G1 at 37°C. Right: Flowchart illustrating different regimes of TSA treatment. To generate TSA-G0 cells, cells are treated with TSA while they are being starved of nitrogen. To generate G0-TSA cells, G0 cells are treated with TSA. (B) Differential interference contrast (DIC, left) and fluorescence (right, DNA) images of DAPI-stained proliferating (P) cells (most in G2 phase), cells arrested at G1, G0 cells, TSA-treated cells – either treated while they were entering G0 (TSA-G0) or treated after they were already in G0 (G0-TSA). (C) Spot tests of proliferating, G0 and TSA-G0 wild type cells after a 30-minute incubation in their respective growth media (YES for proliferating, EMM−N for G0 and TSA-G0), with and without heat stress. (D) Spot tests of proliferating and G0 *cdc10-129* cells after a 30-minute incubation in their respective growth media (YES for proliferating, EMM−N for G0). The −Heat and +Heat conditions are, respectively, 30°C and 48°C.

Even though *S. pombe* G0 cells have been studied for nearly three decades, many of their cell-biological differences with proliferating cells are unknown; these differences include histone post-translational modifications (histone marks), RNA polymerase localisation and abundance, active-transcription status, and nuclear macromolecular packing. Chromatin is one of the most abundant nuclear components and it controls many of the cell’s essential functions. The basic unit of chromatin is the nucleosome, consisting of approximately 147 bp of DNA wrapped around a histone octamer (Kornberg, 1974; Luger et al., 1997). Nucleosomes resemble ∼10 nm wide, 6 nm thick cylinders and are the most abundant macromolecular complexes in the nucleus. Biochemical factors like histone post-translational modifications control nucleosome packing, which may in turn either inhibit or enable the access of nuclear macromolecular machinery (Luger et al., 2012; Maeshima et al., 2019). Consistent with this idea, repressive and activating histone marks are associated with more compact and open chromatin, respectively (Maeshima et al., 2019; Oomen and Dekker, 2017).

Electron cryotomography (cryo-ET) is a popular form of cryo-EM used to study macromolecular complexes *in situ* in a life-like frozen-hydrated state (Bauerlein and Baumeister, 2021; Bohning and Bharat, 2021; Hylton and Swulius, 2021; Ng and Gan, 2020; Schur, 2019). In a cryo-ET experiment, cells are rapidly frozen, thinned, and then imaged in a way to produce a 3-D image called a cryotomogram. We previously used cryo-ET to study *S. pombe* mitotic chromosomes, which have less conspicuous chromatin compaction than what was seen by fluorescence microscopy: with no well-defined boundary between compacted chromatin and nucleoplasm because the nucleus is crowded with macromolecular complexes (Cai et al., 2018b). We also found that prometaphase cells have fewer chromatin-associated multi-megadalton globular complexes (herein called megacomplexes) than interphase cells. The open *in situ* chromatin organisation in *S. pombe* cell nuclei is consistent with mitotic transcription, indicated by the presence of conserved active RNA polymerase II post-translational markers.

In addition to nuclear macromolecular packing, nucleosome structure itself may have a role in transcriptional regulation *in situ*. Recently, we used cryo focused-ion beam (cryo-FIB) milled cellular cryolamellae samples, an energy-filtered electron-counting camera, and 3-D subtomogram classification to do a cryo-ET study of budding yeast *Saccharomyces cerevisiae* nucleosome structure *in situ*. We found that canonical nucleosomes (those that resemble the crystal structure) account for less than 10% of the expected total in *S. cerevisiae* nuclei (Tan et al., 2023). This result contrasts with our findings in human chromatin structure *in situ*, which has abundant and densely packed canonical nucleosomes in both interphase and mitotic cells (Cai et al., 2018a; Chen et al., 2023). Our previous analysis of *S. pombe* cryosections led us to conclude that these cells may not have abundant canonical nucleosomes; this hypothesis has not been rigorously tested using improved sample-preparation techniques (cryo-FIB-milled lamellae) and better cameras. The large differences between yeast and human chromatin could potentially be explained by large differences in transcription levels. Therefore, combined analysis of nucleus cytology, biochemistry, and ultrastructure may provide clues about how extremely different cell states are regulated.

Here we characterise *S. pombe* cell nuclei in multiple states: G1-arrested cells and proliferating cells as controls compared to G0 cells with and without hyperacetylated histones. To fill in several knowledge gaps about G0 nuclear cell biology, we analyse *S. pombe* using complementary methods: fluorescence microscopy, RNA labelling, immunoblots, and *in situ* cryo-ET. Our study correlates histone acetylation, thermotolerance, transcription, nucleus size, nuclear macromolecular packing, and nucleosome structure. We find that G0 histones can be made hyperacetylated, which inhibits nucleus shrinkage, but does not affect thermotolerance or transcriptional repression. Furthermore, neither hypoacetylation nor hyperacetylation affect large-scale nuclear macromolecular packing or the abundance of canonical nucleosomes. Overall, we conclude that G0 transcriptional repression is robust to perturbations in histone acetylation and nuclear morphology and does not lead to a massive upregulation of canonical nucleosomes.

## RESULTS

### *S. pombe* strains and cell-cycle states

We have analysed how nuclear morphology, histone acetylation, transcription, and nuclear macromolecular complex distribution differ in proliferating and G0 *S. pombe* cells. This study therefore required cells of different strains, cell-cycle states, or combinations thereof (**Table S1)**. Strain MBY99 (herein called wild-type) was used as wild-type and for the generation of new strains that have eGFP fusions to genes at their endogenous loci (described later). Wild-type cells were used to generate asynchronous proliferative cells (herein called proliferating) and G0 cells both without and with hyperacetylated histones. Some experiments were done with the mutant strain *cdc10-129*, which arrests in G1 at elevated temperature. For nascent-RNA labelling experiments, we used the strain yFS240 (Sivakumar et al., 2004), which can incorporate exogenous uridine and its analogues (Hodson et al., 2003). For some cryo-ET controls, we used temperature-sensitive *cdc10-129* cells arrested in G1 phase (Aves et al., 1985) because G1 and G0 cells both have a 1N nuclear DNA content and cells enter G0 from a G1-like state (Mochida and Yanagida, 2006). We also used proliferating cells, which are mostly in G2, the longest cell-cycle phase (Forsburg, 1999). We did not use the strain *cdc10-129* for studies of G0 because when these cells were transferred to EMM−N, they did not have G0 phenotypes (see below).

### Preparation of G0 and TSA-G0 cells

To prepare G0 *S. pombe* cells (**Figure 1A**), we starved proliferating cells of nitrogen by incubating them in Edinburgh Minimal Media without nitrogen (EMM−N) for 24 hours (Costello et al., 1986; Su et al., 1996). G0 cells were prepared from wild-type (972 h-) cells, yFS240 cells, and new strains that express either an eGFP-tagged RNA polymerase II subunit or an eGFP-tagged nuclear pore subunit (details in later sections). To enable the study of histone-hyperacetylation effects, we prepared G0 cells by adding Trichostatin A (TSA) to the EMM−N. TSA is a histone-deacetylase inhibitor (Rundlett et al., 1996) that increases histone acetylation levels in proliferating *S. pombe* cells (Kimata et al., 2008) and induces chromatin decompaction *in situ* in human cells (Toth et al., 2004). To determine if TSA treatment affects G0 chromatin and if these changes depend on the timing of TSA treatment, we subjected *S. pombe* cells to two different regimens. In the TSA-G0 regimen, we incubated proliferating cells in EMM−N plus TSA for 24 hours. In the G0-TSA regimen, we first incubated proliferating cells in EMM−N for 24 hours, then we incubated the cells in EMM−N plus TSA for another 24 hours.

Differential interference contrast microscopy showed that following 24 hours of nitrogen starvation, wild-type G0 cells became shorter (**Figure 1B**) as reported earlier (Su et al., 1996), while fluorescence microscopy of DNA stained with 4′,6-diamidino-2-phenylindole (DAPI) showed that G0 chromosomes did not individualise like in prometaphase-arrested *nda3-KM311* cells (Hiraoka et al., 1984). TSA treatment leads to diverse cell morphologies depending on the genetic background. TSA-G0 wild-type cells were longer than G0 cells but shorter than proliferating cells. G0-TSA cells were shorter like untreated G0 cells (**Figure 1B**). Owing to their simpler preparation, most of our analyses of TSA-treated G0 cells were done with TSA-G0 cells, for which TSA treatment and nitrogen starvation are concurrent during G0 entry. G0 and TSA-G0 cells were less sensitive to 48°C heat stress than proliferating cells (**Figure 1C**), meaning that TSA-treated G0 cells retain an important G0 phenotype. In contrast, *cdc10-129* cells that were transferred to EMM−N were less resistant to heat shock than wild-type G0 cells (**Figure 1D**). Other strains, including yFS240, an Rpb1-eGFP strain, and a Nup97-eGFP strain were all resistant to heat shock and had cell-to-cell heterogeneity after TSA-G0 treatment, with cells longer than TSA-G0 wild-type cells (details in the relevant sections below). To keep the number of phenotypes manageable, we limited the analysis of TSA-G0 treatment to wild-type cells.

### G0 chromatin is hypoacetylated, but can be reversibly hyperacetylated

Histone marks are correlated with chromatin structure and function in yeasts (Millar and Grunstein, 2006; Sinha et al., 2006) and has been characterised in only a few extreme cell states. *S. cerevisiae* G0 cells has altered levels of several histone marks compared to proliferating cells (McKnight et al., 2015; Mews et al., 2014) and also globally repressed transcription (Allen et al., 2006; McKnight et al., 2015). Notably, *S. cerevisiae* G0 cells have lower histone acetylation, which is correlated with oligonucleosome condensation (Robinson et al., 2008; Shogren-Knaak et al., 2006; Tse et al., 1998). We performed immunoblot analysis of *S. pombe* histone acetylation and found that compared to proliferating and G1 cells, the G0 cells had lower levels of H3 acetylated at multiple N-terminal lysines (H3-Ac), but not H4 acetylated at multiple N-terminal lysines (H4-Ac) or at K16 (H4K16ac) (**Figure 2A**). TSA treatment with either the TSA-G0 or G0-TSA regime induced hyperacetylation of H3-Ac, H4-Ac, and H4K16ac at levels equal to or greater than in G0 and even greater than in G1 and proliferating cells. This TSA-induced G0 histone hyperacetylation was reversed after TSA washout from TSA-G0 cells (**Figure 2A**).

**Figure 2.**
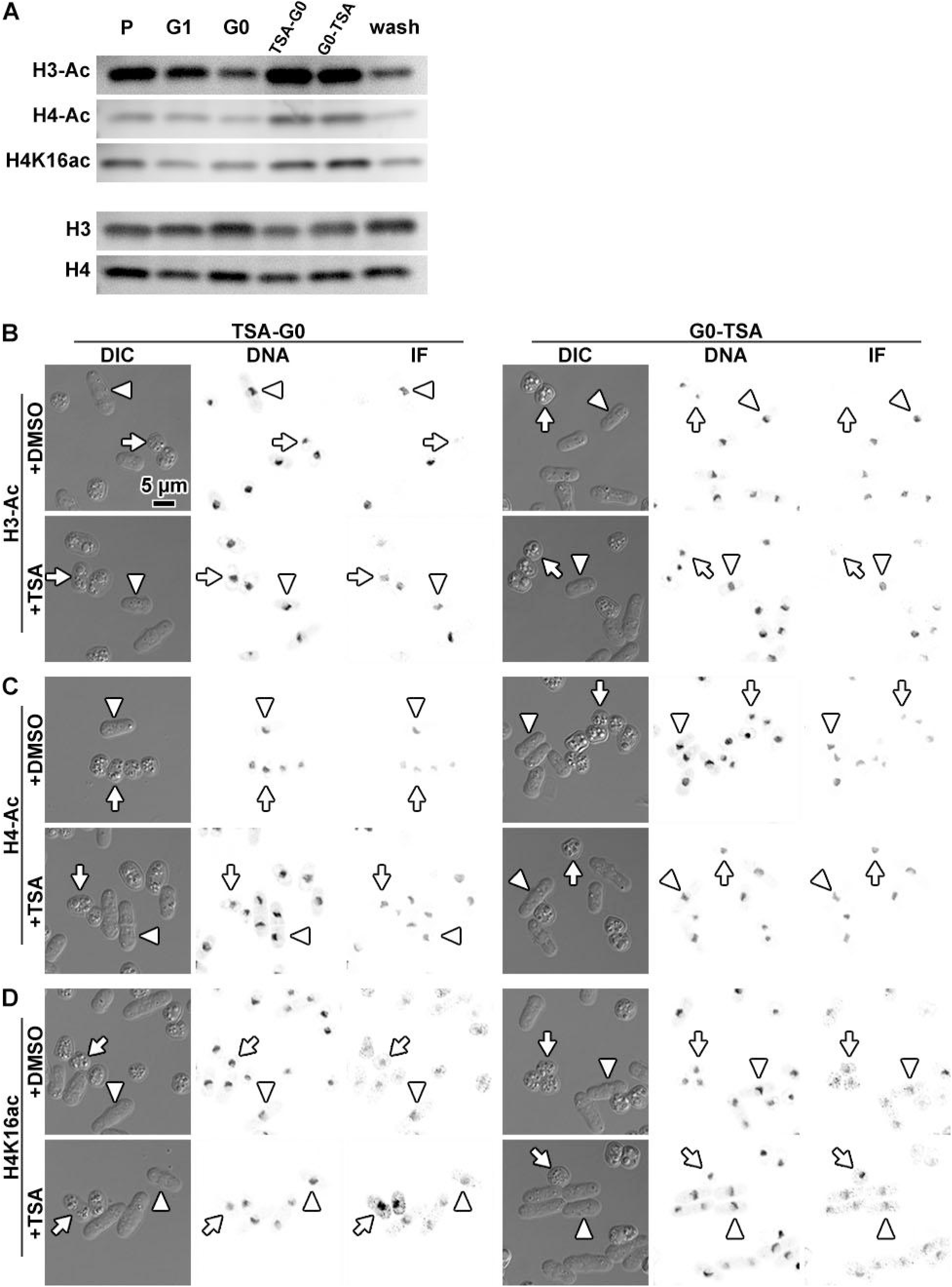
Histone H3 acetylation levels decrease in G0. (A) Immunoblots of *S. pombe* lysates from wild-type proliferating (P), *cdc10-129* G1, and wild-type G0 cells. The G0 cells were either untreated (G0), incubated with TSA in EMM−N for 24 hours (TSA-G0), incubated 24 hours in EMM−N followed by 24 hours of EMM−N plus TSA (G0-TSA), or treated with EMM−N plus TSA for 24 hours followed by TSA washout and then incubated an additional 24 hours in EMM−N (wash). TSA was used at 20 µg/mL TSA final concentration, with DMSO as the carrier such that the final concentration of DMSO in the culture is 0.1%. Loading controls were done with antibodies against the H3 or H4 C-terminus. The uncropped immunoblots are shown in Figure S29. G0 cells were subjected to immunofluorescence imaging to detect (B) H3-Ac, (C) H4-Ac, and (D) H4K16ac. The G0 cells were treated with DMSO (+DMSO) or TSA (+TSA). “TSA-G0” and “G0-TSA” denote TSA treatment during or 24 hours after G0 entry, respectively, like in the immunoblot experiment. To ensure the cells were processed in similar conditions, the G0 and proliferating cells were first fixed separately, then mixed together for immunofluorescence processing. The proliferating cells serve as a common reference. In each subpanel, a representative G0 cell is indicated by an arrow and a representative proliferating cell is indicated by an arrowhead.

We next performed immunofluorescence to check if there is gross cell-to-cell variability i.e., if a subset of G0 cells had much-more histone acetylation than the others. We found that nearly all G0 cells had lower levels of H3-Ac than the typical proliferating cell (**Figure 2B**; compare short vs long cells in the +DMSO rows). In contrast, neither H4-Ac nor H4K16ac levels decreased noticeably in G0 cells (**Figure 2, C and D**; +DMSO rows). Both the immunoblot and immunofluorescence experiments showed that overall histone acetylation decreased in G0. In contrast, both immunoblots and immunofluorescence microscopy (**Figure 2, B – D**; +TSA rows) showed that both TSA-G0 and G0-TSA cells had equal or greater H3 and H4 acetylation than either untreated G0 cells or proliferating cells.

### Transcription is repressed in G0 cells with and without histone hyperacetylation

An early study used RNA-seq to show that G0 cells have fewer transcripts than proliferating cells (Marguerat et al., 2012). To further assess the differences in transcription phenotypes between interphase and G0 cells, we analysed the abundance of RNA polymerase II, active RNA polymerase II, and newly synthesised RNA. In *S. pombe*, the *rpb1* gene encodes the largest subunit of the RNA polymerase II complex and exists as a single copy (Kimura et al., 1997). We tracked RNA polymerase II levels and localisation *in vivo* by creating a strain that has the *rpb1* gene tagged at its C-terminus with eGFP (**Figure S1**). In confocal fluorescence micrographs of living cells, we found that both proliferating and G0 cells had bright Rpb1-eGFP nuclear signals (**Figure 3A**). RNA polymerase II levels are therefore unlikely to be downregulated in G0.

**Figure 3.**
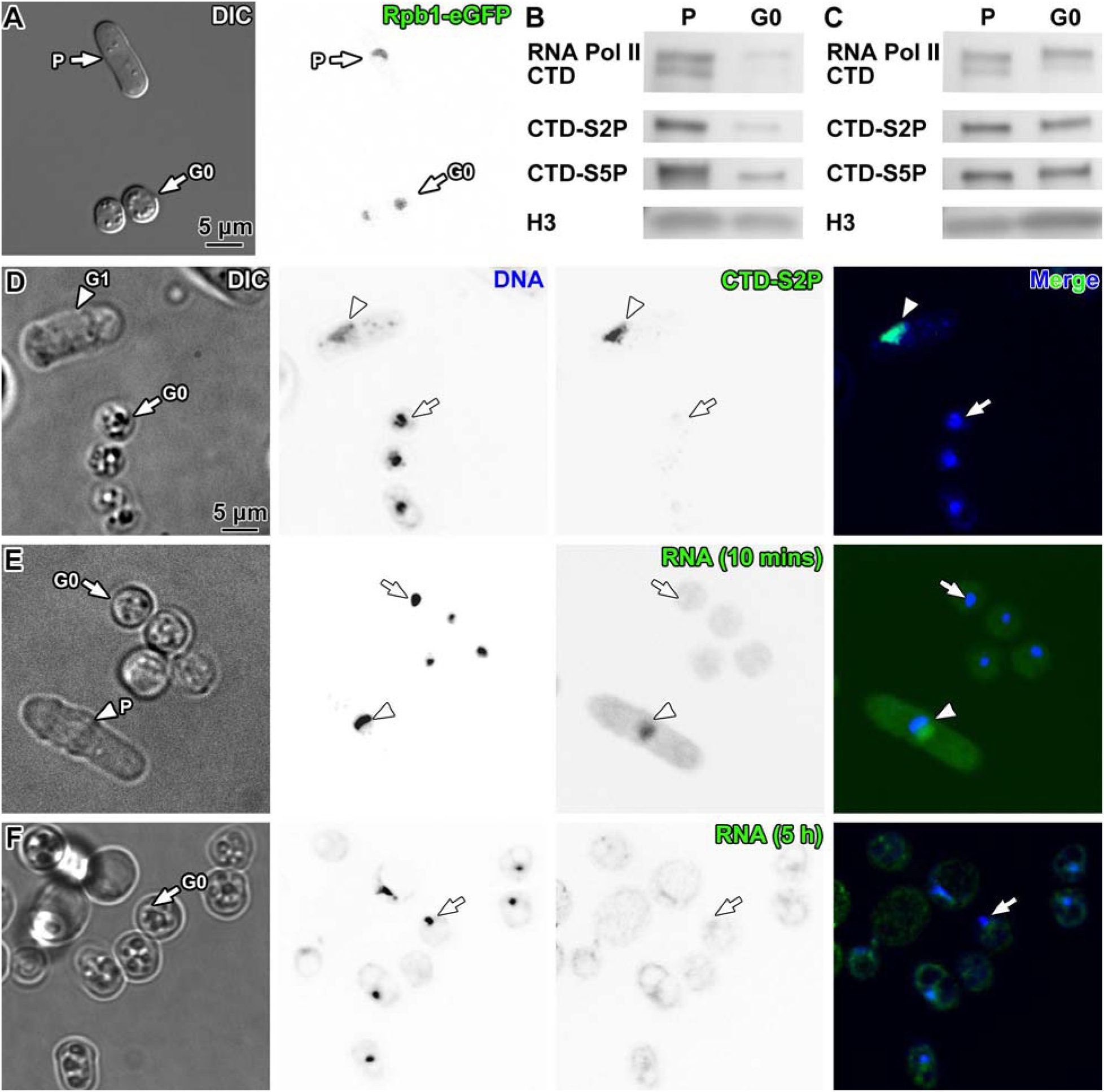
Transcription and RNAPII levels are lower in G0 cells. Fluorescence microscopy analysis of transcription in interphase and G0 cells. (A) Differential interference contrast (DIC) (left) and eGFP (right) live-cell fluorescence microscopy images of Rpb1-eGFP in proliferating (P) and G0 states, in a strain where the *rpb1* gene is tagged with eGFP. Interphase cells were mixed into G0 cell culture just before imaging. A proliferating cell and a G0 cell are indicated. (B) Immunoblots of lysates from proliferating (P) and G0 wild type cells, using an α-CTD antibody, an α-CTD-S2P antibody and an α-CTD-S5P antibody. Loading controls were done with an antibody against the H3 C-terminus. (C) Immunoblots of the same lysates and antibodies as part B, with loading calibrated to have equal α-CTD signal. Uncropped immunoblots are shown in Figure S29. Columns, left to right for parts D-F: differential interference contrast (DIC), DAPI fluorescence (DNA), Alexa Fluor 488 fluorescence (CTD, CTD-S2P, and RNA), and merge. (D) Detection of elongating RNA polymerase II with the anti-CTD-S2P antibody. One G1 cell is indicated by an arrowhead and a G0 cell by an arrow. (E) Strain yFS240 G0 and proliferating (P) cells were incubated with 5-EU and subjected to the Click-iT 5-EU-detection assay, which ligates Alexa Fluor 488 to 5-EU. A proliferating cell is indicated by an arrowhead and a G0 cell by an arrow. To increase the visibility of the weak 5-EU signal, the contrast was adjusted for the entire field of view, resulting in a higher apparent background. (F) G0 cells were incubated with 5-EU for 5 hours, then subjected to Click-iT 5-EU detection as in panel E. One G0 cell is indicated by an arrow. The two larger cells in the upper left are probably dead or dying as a result of prolonged 5-EU exposure.

RNA polymerase II activity may also be controlled by interactions with other transcription machinery, resulting in detectable changes to this enzyme’s post-translational state. So we next performed immunoblots against the Rpb1 subunit’s carboxy-terminal domain (CTD) as well as two post-translational modifications of the Rpb1 CTD, serine 2 (CTD-S2P) and serine 5 (CTD-S5P) phosphorylation. The two phosphorylation marks are conserved markers for transcriptional elongation and transcription initiation respectively (Harlen and Churchman, 2017; Komarnitsky et al., 2000). Immunoblots that were calibrated using total H3 levels showed that G0 cells have less Rpb1 (consistent with the Rpb1-eGFP analysis), CTD-S2P, and CTD-S5P marks than proliferating cells, suggesting that RNA polymerase II initiation and elongation were decreased (**Figure 3B**). Immunoblots that were calibrated to equal CTD levels (using a pan-CTD antibody) showed that G0 cells also had similar levels of both CTD-S2P and CTD-S5P marks per Rpb1 subunit (**Figure 3C**). To characterise G0 RNA polymerase II activity in single cells, we performed immunofluorescence imaging to detect CTD-S2P. To keep the chromatin content comparable, we used *cdc10-129* cells arrested at G1 rather than asynchronous cells. We found that G0 cells had weaker CTD-S2P signals than G1 cells (**Figure 3D**). Therefore, Rpb1 is expressed at lower levels in G0 than in proliferating cells, yet the concentrations of this protein and activity of the enzyme to which it belongs are non-negligible in G0 cell nuclei.

To characterise bulk RNA synthesis in G0 cells, we fluorescently labelled new transcripts in the strain yFS240 (Sivakumar et al., 2004). Strains like yFS240 can incorporate exogenous uridine and its analogues into nascent RNA (Hodson et al., 2003). Newly synthesised RNA molecules that incorporate 5-ethynyl-uridine (5-EU) become detectable after ligation with a fluorescent dye like Alexa Fluor 488 (Jao and Salic, 2008). We therefore treated proliferating yFS240 cells with 5-EU to reveal new transcripts. For negative controls, we incubated wild-type cells with 5-EU, yFS240 cells with 5-EU plus the transcription inhibitor phenanthroline, and yFS240 cells with uridine. All 5-EU and uridine treatments were done for 10 minutes. The chromatin was stained with DAPI. Fluorescence microscopy showed nucleus-localised Alexa Fluor 488 signals above the background only in yFS240 cells that were incubated with 5-EU without transcription inhibitor; no signals were visible in the negative controls (**Figure S2**). In proliferating cells, the newly transcribed RNA appeared more abundant in the hemisphere opposite the DAPI signal (**Figure 3E**). This DAPI-poor (low DNA concentration) region is the *S. pombe* nucleolus (Tanaka and Kanbe, 1986), meaning that the strong 5-EU signal in this hemisphere came from newly synthesised rRNA (**Figure 3E** lower-left cell; and see the nucleolus analysis below). In comparison, 5-EU signal was much weaker in the DAPI-rich (chromatin) hemisphere of the nucleus, where most mRNA is synthesised. Such high levels of rRNA synthesis are consistent with the extremely high levels of ribosome biogenesis in yeasts (Warner, 1999). Compared to proliferating cells, G0 cells had very weak 5-EU signals (**Figure 3E**). To test if 10-minute incubations were too short for sufficient 5-EU-labelled RNA to accumulate in G0 cells, we increased the 5-EU incorporation to 5 hours. The 5-EU-labelled RNA signal thereafter became detectable, albeit weak, in the G0 nucleus (**Figure 3F**). Furthermore, 5-EU was also present in the cytoplasm, possibly from the exported RNAs (**Figure 3F**).

Next, we tested the effect of TSA treatment on transcription in G0 cells. Like untreated G0 cells, TSA-G0 cells had weak CTD-S2P immunofluorescence signal (**Figure S3A**) and weak nuclear 5-EU signals (**Figure S3B**), meaning that transcription in G0 cells remained repressed even when the histones were hyperacetylated. TSA-G0 ‘s Rbp1-eGFP signal was similar to that of G0 cells and fainter than in proliferating cells (**Figure S3C**). Histone hyperacetylation therefore does not have large effects on either the intranuclear localisation or the concentrations of Rpb1 in G0. In summary, RNA transcription is repressed in G0 cells, even when the cells are forced to have histone marks that are associated with transcriptional activity.

### *S. pombe* cells do not undergo rapid transcriptional upregulation after G0 exit

G0 is a reversible state, so G0 exit may reveal clues about the nuclear changes needed to de-repress genes. Many cell types exit G0 when nutrients are restored to the cell culture medium (Marescal and Cheeseman, 2020). Just minutes after G0 *S. cerevisiae* are switched into rich media, they undergo a state called “hypertranscription”, which is detectable as a large increase in transcription markers and nascent RNA (Cucinotta et al., 2021). This finding inspired us to characterise *S. pombe* transcription in G0 exit. We performed immunoblots against the total CTD levels (to account for total RNA polymerase II) and the transcription markers CTD-S2P and CTD-S5P on *S. pombe* exiting G0, and found that compared to histone H3 levels (to calibrate for cell number), total CTD, CTD-S2P and CTD-S5P all remained low for 2 hours, increased at 4 hours, and continued increasing to the highest signal level at 6 hours **(Figure S4A)**. When we compared CTD-S2P and CTD-S5P to total CTD levels, neither CTD-S2P nor CTD-S5P levels increased substantially over 24 hours, with a very minor increase in CTD-S2P levels at 2 hours (**Figure S4B**). These results suggest that transcription in *S. pombe* does not increase as quickly as in *S. cerevisiae* during G0 exit.

To characterise *S. pombe* G0 exit in single cells, we imaged Rpb1 and the transcription markers by fluorescence microscopy. First, we performed live-cell fluorescence time-lapse imaging of G0 Rpb1-eGFP cells deposited on a YES agar pad, which verified that these cells were exiting G0, elongating, and undergoing nuclear division followed by cell division (**Figure S5A; Movie S1**). Since the growth conditions on an agar pad are different from liquid culture, we also analysed Rpb1-eGFP G0 cells suspended in liquid YES medium and collected freshly at different time points. Cells exiting G0 became visibly longer 2 hours after resuspension in rich medium and resembled proliferating cells within 24 hours. The Rpb1-eGFP signal had a very small fluorescence-intensity increase over 6 hours after the rich-media switch (**Figure S5B**). We then subjected yFS240 cells exiting G0 to 5-EU incorporation analysis. Nascent (albeit weak) nuclear RNA signals became visible above background starting at 6 hours after resuspension in rich media (**Figure S6**). In wild-type cells that were exiting G0, levels of the transcription-elongation marker CTD-S2P were weak until 6 hours after the rich-media switch (**Figure S7**). Cells exiting G0 showed weak immunofluorescence signals for the transcription-initiation marker CTD-S5P until 4 hours after the rich-media switch (**Figure S8**) and weak CTD levels until 6 hours after the rich-media switch (**Figure S9**). These combined results indicate the *S. pombe* does not undergo hypertranscription during G0 exit and that transcription markers become non-negligible approximately when cell division restarts.

### G0 nucleus shrinkage is inhibited by TSA treatment

A popular hypothesis of transcriptional regulation is that nuclear macromolecular crowding sterically inhibits the transcription machinery. TSA-treated G0 cells have repressed transcription despite their increased histone acetylation, raising the question: do TSA-G0 cells have less crowded nuclei? The confocal microscopy images above show hints that the larger TSA-G0 cells have bigger nuclei, whose spaciousness may allow for less macromolecular crowding. However, it is difficult to estimate nuclear volume using fluorescence microscopy of DAPI signals because the nuclear boundary is ambiguous. A better strategy is to image a strain that expresses a fluorescent nuclear-envelope protein, measure the nucleus diameter, and then calculate the nuclear volume (Heun et al., 2001; Neumann and Nurse, 2007; Varberg et al., 2022). We therefore created an *S. pombe* strain in the wild-type background in which the nuclear-pore protein Nup97 is tagged with eGFP at its endogenous locus (**Figure S10**). We did not use the *cdc10-129* background because live-cell imaging allows for the selection of G1 cells in a wild-type background – specifically, G1 cells are those that have just finished nucleokinesis. Fluorescence microscopy analysis of Nup97-eGFP-expressing G0 cells confirmed that their nuclei were smaller, with approximately half the nuclear volume of G1 cells (mean volume of G1 nuclei = 5.2 µm^3^, n = 52, and G0 nuclei = 2.8 µm^3^, n = 114; p < 0.0001) (**Figure S11; Movie S2**). We also treated Nup97-eGFP-expressing cells with TSA either during or after entry into G0, then followed up with fluorescence microscopy (**Figure S12, A – C**). TSA-G0 cell nuclei had approximately twice the volume of G0 cell nuclei (mean volume of TSA-G0 nuclei = 5.7 µm^3^, n = 90; p < 0.0001) (**Figure S12D**). In contrast, when the TSA was added to cells that have already entered G0, the resulting G0-TSA nuclei had ∼30% more volume than G0 nuclei (G0-TSA nucleus mean volume = 3.7 µm^3^, n = 138; p < 0.0001) (**Figure S12E**). These results show that nuclei are enlarged more due to TSA treatment while the cell is entering G0 than after it has already entered G0.

### Cryo-ET analysis of *S. pombe* cells

To characterise the ultrastructural details of nuclei in proliferating, G1, G0 and TSA-G0 cells, we used cryo focused-ion-beam milling to prepare cellular cryolamellae (most of which were thinner than 150 nm) and then imaged them by cryo-ET. Defocus phase-contrast cryotomograms of nuclei in G1 and G0 cells revealed crowds of macromolecular complexes (**Figure S13, A and B**), which were previously not visible (Sajiki et al., 2009; Su et al., 1996). The contrast of the G0 nuclear densities in these cryotomograms was lower than in the G1 cells (**Figures S13, C and D**). This contrast difference is likely inherent in the sample and not due to sample preparation because the G0 cell lamella was thinner than the G1 cell lamella (∼135 vs ∼152 nm) and should therefore have the higher contrast expected of thinner samples. Subtomogram analysis of small complexes like nucleosomes is facilitated when the data are collected with a Volta phase plate (VPP), which increases the low-resolution contrast (Asano et al., 2015; Fukuda et al., 2015; Tan et al., 2023). Therefore, the cryo-ET data reported below were collected with the VPP.

### Canonical nucleosomes are rare in Proliferating, G1, G0 and TSA-G0 cells

Our previous attempts to identify canonical nucleosomes *in situ* in VPP cryotomograms of *S. pombe* cryosections did not reveal canonical class averages (Cai et al., 2018b). However, that study used cryosections, which may have sample-preparation-induced structural artefacts, and previous-generation electron detectors, which have lower signal-to-noise ratios (Ruskin et al., 2013). Because the *S. pombe* lysates did reveal canonical nucleosome class averages, we hypothesised that the nucleosomes in these cells may have partially detached DNA *in situ*. Our recent *in situ* study of wild-type *S. cerevisiae* (Tan et al., 2023), which used cryolamellae, a K3 detector, and a VPP, showed that canonical nucleosomes account for < 10% of the expected total, meaning that non-canonical nucleosomes (whose structures remain unknown) are the majority in budding yeast.

To characterise the prevalence of canonical nucleosomes of *S. pombe*, we performed VPP cryotomography of proliferating, G1, G0, and TSA-G0 cell cryolamellae (**Figures 4, S14 – S23**). We then used a workflow that involves template matching followed by 3D classification to attempt to identify and locate the canonical nucleosomes among the nucleosome-like particles (Tan et al., 2023). In this workflow, we minimised model bias by using a smooth cylindrical reference for both template matching and classification. Canonical nucleosome class averages were not found *in situ* in proliferating, G1, or TSA-G0 *S. pombe* cell tomograms (**Figures S24, S25, S27**). Note that the absence of canonical nucleosome class averages in the other cell states reflects the rarity – but not the absence – of canonical nucleosomes. Only the G0 cell tomograms had a class average that vaguely resembles the canonical nucleosome (**Figures 5A, S26**). While the refined canonical nucleosome-like class average vaguely resembles the canonical nucleosome crystal structure (White et al., 2001), it has jagged features (**Figure 5, B and C**). These unusual density features suggest that 3-D classification could not resolve all the heterogeneity, meaning that canonical nucleosomes are also rare in G0 (less than ∼4% of the expected population). Further advances in classification will be needed to improve the estimation of canonical-nucleosome structure abundance.

**Figure 4.**
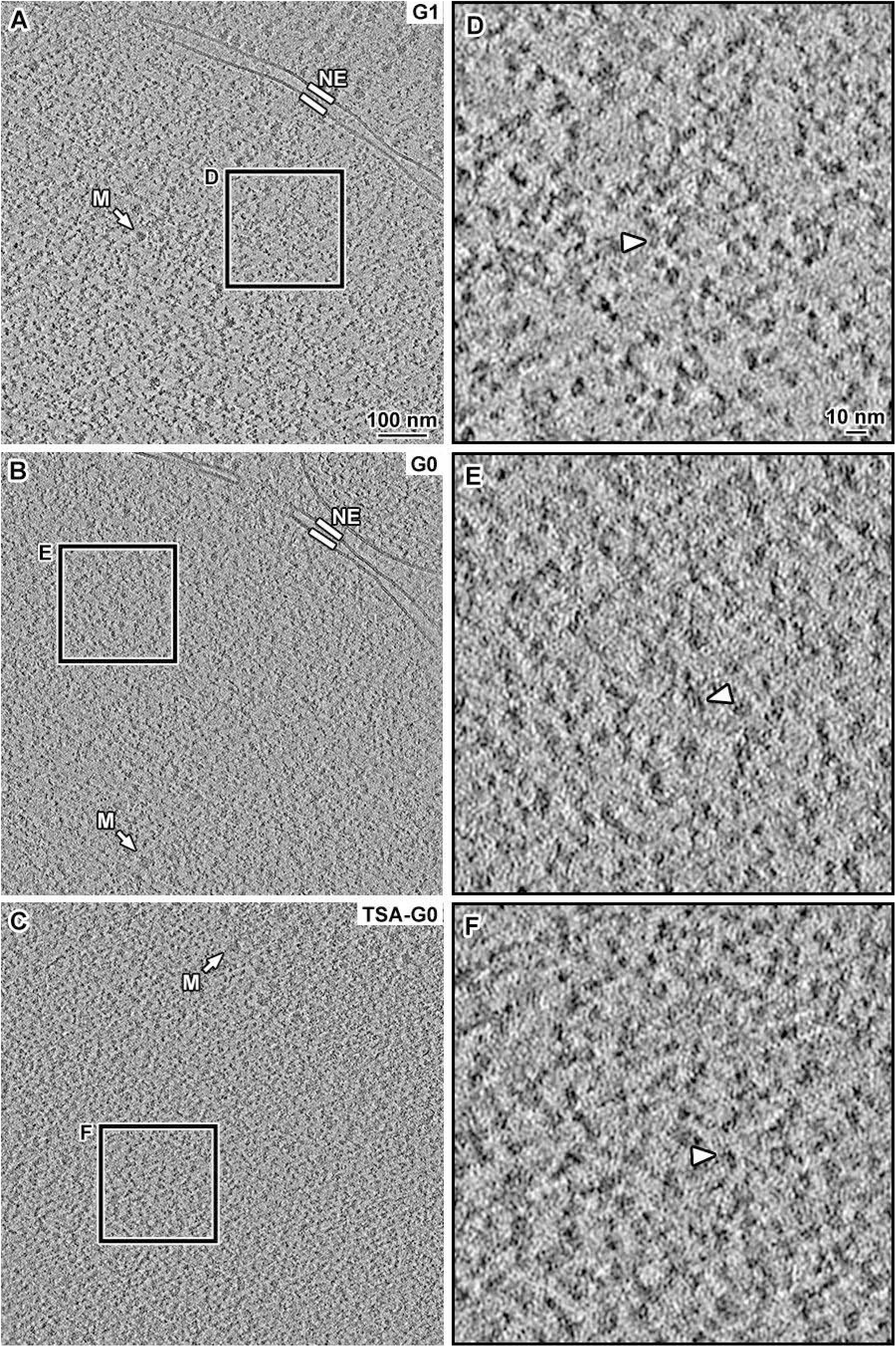
G1, G0 and TSA-G0 cells have dense nucleoplasm. (A) Volta cryotomographic slice (12 nm) of the nucleus in a G1-arrested *cdc10-129* cell. The nuclear envelope (NE) and a megacomplex (M) are indicated. (B) Volta cryotomographic slice (12 nm) of a nucleus in a wild-type G0 cell treated with 0.1% DMSO during G0 entry. The nuclear envelope (NE) and a megacomplex (M) are indicated. (C) Volta cryotomographic slice (12 nm) of the nucleus in a G0 cell treated with 20 µg/mL TSA during G0 entry. (D, E and F) Four-fold enlargements of the regions boxed in panels A, B and C, respectively. Nucleosome-like densities are indicated by arrowheads. Larger fields of view of G1, G0 and TSA-G0 nuclei are available in supplemental figures S15, S19 and S22 respectively.

**Figure 5.**
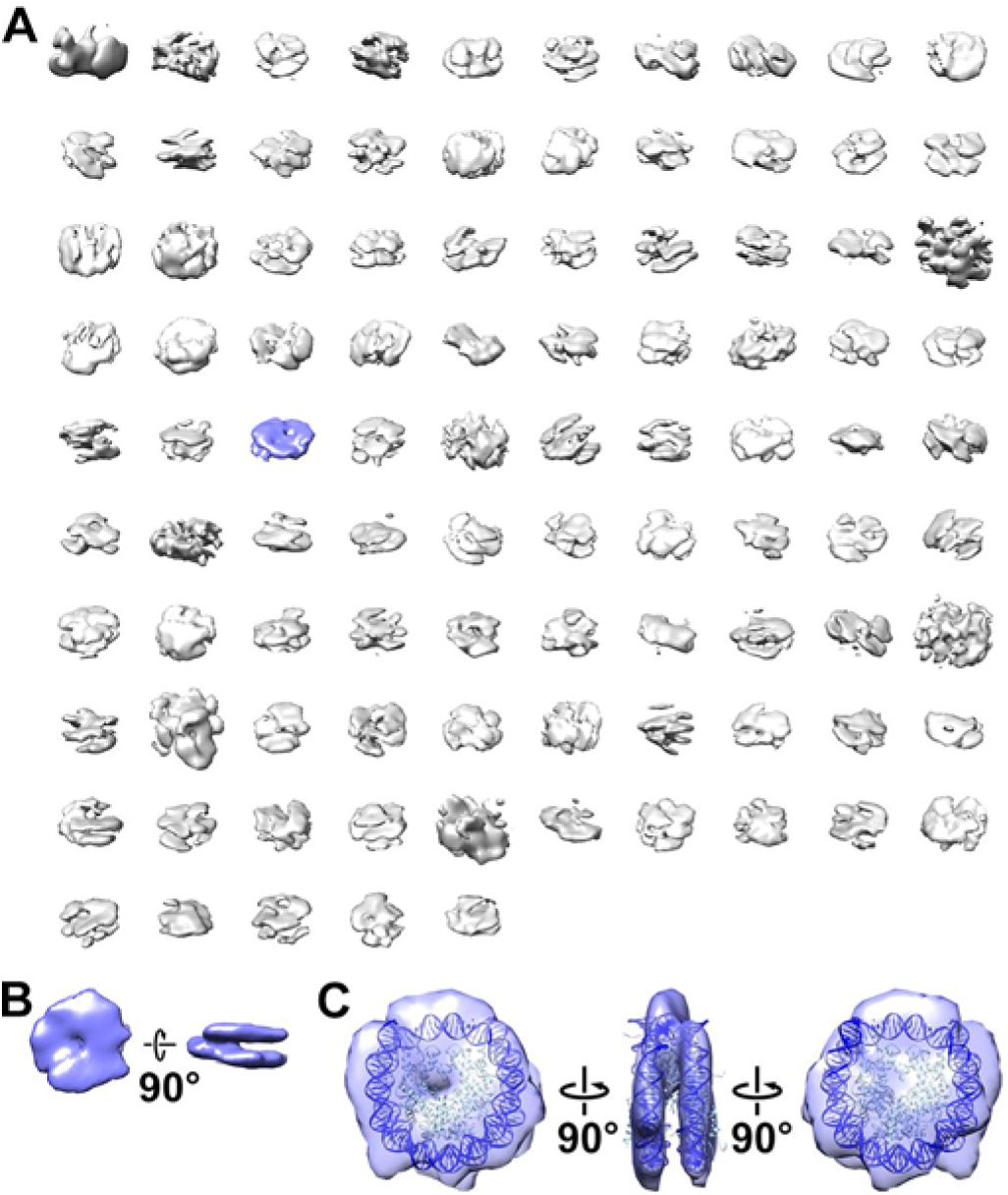
A class average that partially resembles the canonical nucleosome is detected in DMSO-G0 cells *in situ*. (A) Class averages (3-D) of nucleosome-like particles in Volta phase plate (VPP) cryotomograms of DMSO-G0 cryolamellae. The class shaded blue partially resembles a canonical nucleosome. Five classes have no contributing particles. (B) Refined density of candidate nucleosomes observed in DMSO-G0 cryolamellae. (C) Refined density of the candidate G0 *S. pombe* nucleosome class with the crystal structure of the *Saccharomyces cerevisiae* nucleosome (PDB 1ID3) (White et al., 2001) docked. The histones and DNA are light and dark blue respectively. The *S. cerevisiae* nucleosome was used because the crystal structure of the *S. pombe* nucleosome has not been determined. For more details of the workflow, see Figure S26.

### G1, G0 and TSA-G0 nuclei have similar macromolecular complex crowding

Our light-microscopy experiments above showed that G0 cells have smaller nuclei than both G1 (**Figure S11**) and TSA-G0 cells (**Figure S12**), even though they all have the same number of chromosomes. We thus expected that G0 cells would show denser nucleoplasmic particle packing than both G1 and TSA-G0 cells, which would reflect differences in chromatin compaction. Previously, we assumed that most nucleosomes were canonical (not detectable due to suspected technical limitations discussed above), so we had used nearest-neighbour density analysis of nucleosome-like particles to assess chromatin packing in nuclei (Cai et al., 2018b). Here our study rectifies both the sample-preparation and data-quality issues and now shows that non-canonical nucleosomes are the majority species *in situ*. Therefore, we cannot use nearest-neighbour-distance analysis to study chromatin packing because we cannot yet identify the non-canonical nucleosomes. Before nucleosome identification *in situ* in cryo-EM and cryo-ET studies was feasible, Fourier analysis was used to characterise the packing density of macromolecular complexes (Chen et al., 2016; Eltsov et al., 2008; Gan et al., 2013; Scheffer et al., 2011). We therefore evaluated the overall packing of nuclear complexes using radially averaged Fourier power spectra (herein power spectra), which is less sensitive to the identities of the nuclear complexes.

The *S. pombe* nucleus has two major compartments – the nucleolus, which is the site of ribosome biogenesis, and the region outside the nucleolus, which has more chromatin. Nuclear megacomplexes include preribosomes, transcription initiation complexes, spliceosomes, and many other globular complexes that exceed one megadalton molecular weight. We first analysed cryolamella tomograms from the G1 cell cytoplasm as a control (**Figure S14**). Power spectra from the ribosome-regions showed a broad peak at approximately 30 nm spacing (**Figure 6A)**, which reflects the densely packed ribosomes (25 – 28 nm) (Verschoor et al., 1998). To compare the crowding (average centre-to-centre spacing) of all nuclear complexes, we performed Fourier analysis on the G1 (**Figures S15 and S16**), proliferating (**Figures S17 and S18**), G0 (**Figure S19 and S20**) and TSA-G0 (**Figure S22 and S23**) nucleus cryotomograms. The power spectra showed that the nuclei from all four cell types have a broad peak centred around 20 nm (**Figure 6**). As such, proliferating, G1, G0 and TSA-G0 nuclei have similar macromolecular crowding in the nucleoplasm. Furthermore, when the nucleoplasm (**Figure S20**) and the nucleolus (**Figure S21**) in the same G0 nucleus were compared, the power spectra were indistinguishable (**Figure S28**), meaning that these two nuclear subcompartments have similar crowdedness.

**Figure 6.**
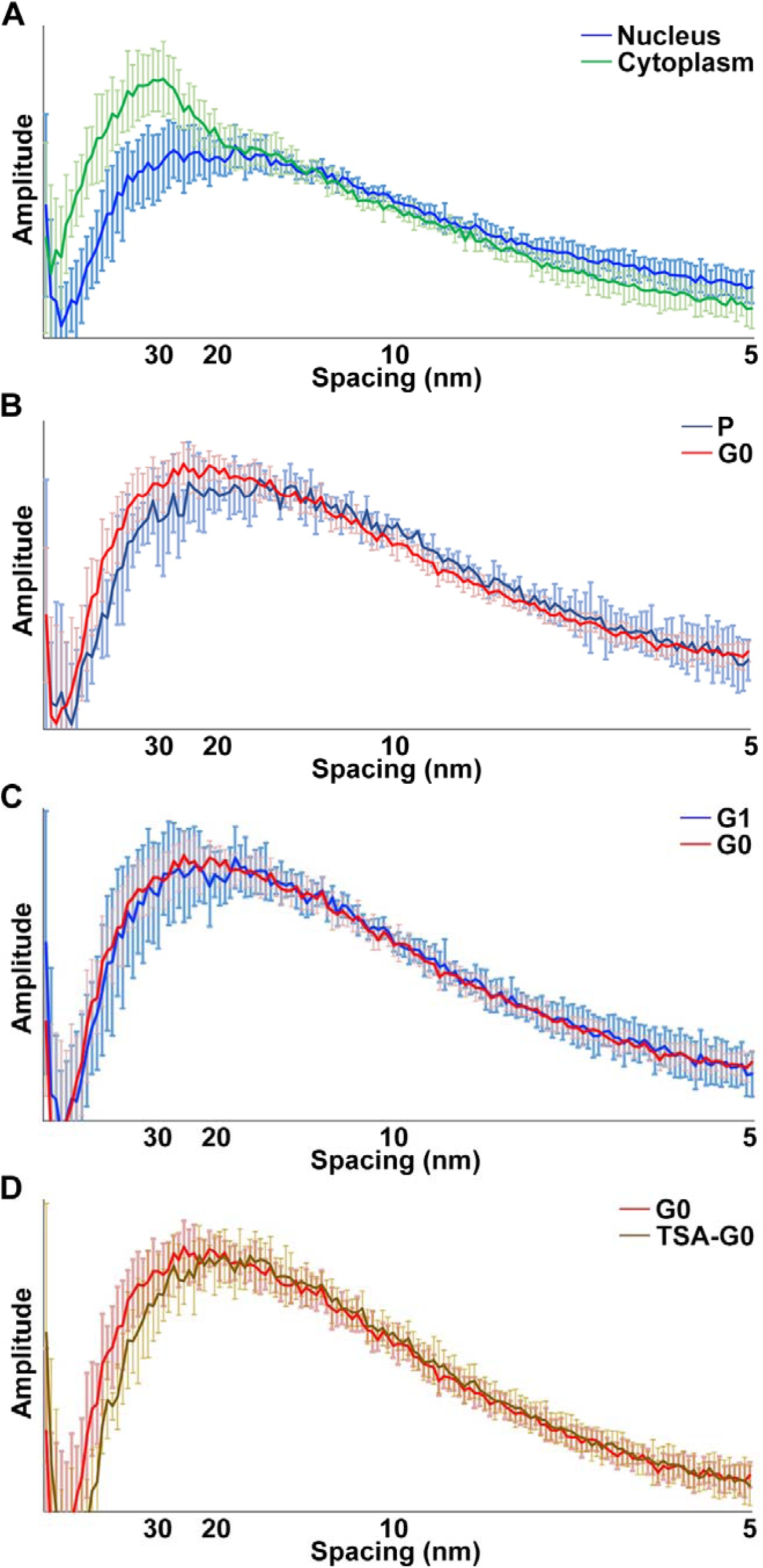
*S. pombe* nuclei do not have large packing differences in various states. Average 1-D Fourier power spectra comparing (A) G1 nuclei (blue) and G1 cytoplasm (green); (B) proliferating (dark blue) and G0 (red) nuclei; (C) G1 (blue) and G0 (red) nuclei; (D) G0 (red) and TSA-G0 (brown) nuclei. For each spacing value, the normalised amplitudes are plotted; see Methods. Error bars indicate the standard deviation at each spacing.

### G0 and TSA-G0 chromatin has fewer megacomplexes than G1

*S. pombe* cells also undergo large nuclear changes when they enter mitosis. Compared to G2 (interphase) cells, prometaphase *S. pombe* cells have weaker (but still detectable) CTD-S2P signals, compacted chromosomes, and fewer megacomplexes in the chromatin (Cai et al., 2018b). These observations suggest a weak positive correlation between megacomplex abundance and transcription. To further explore the relationship between megacomplexes and transcription states, we sought to estimate the concentration of nucleoplasmic megacomplexes in G1, G0 and TSA-G0 cells. Since megacomplexes are too heterogeneous for 3D classification, we manually annotated globular particles larger than ∼15 nm and found that megacomplexes were more abundant in G1 chromatin than in G0 chromatin (mean concentration of G1 = 665 megacomplexes per femtoliter, n = 5; mean concentration of G0 = 179 megacomplexes per femtoliter, n = 10; p = 0.0002) (**Figure 7A**). The G0 nucleoplasmic megacomplex concentration was similar to that of TSA-G0 (mean concentration of G0 = 179 megacomplexes per femtoliter, n = 10; mean concentration of TSA-G0 = 155 megacomplexes per femtoliter, n = 7, p = 0.57) (**Figure 7B**). Therefore, both TSA-G0 and G0 cells have lower concentrations of chromatin megacomplexes than G1 cells.

**Figure 7.**
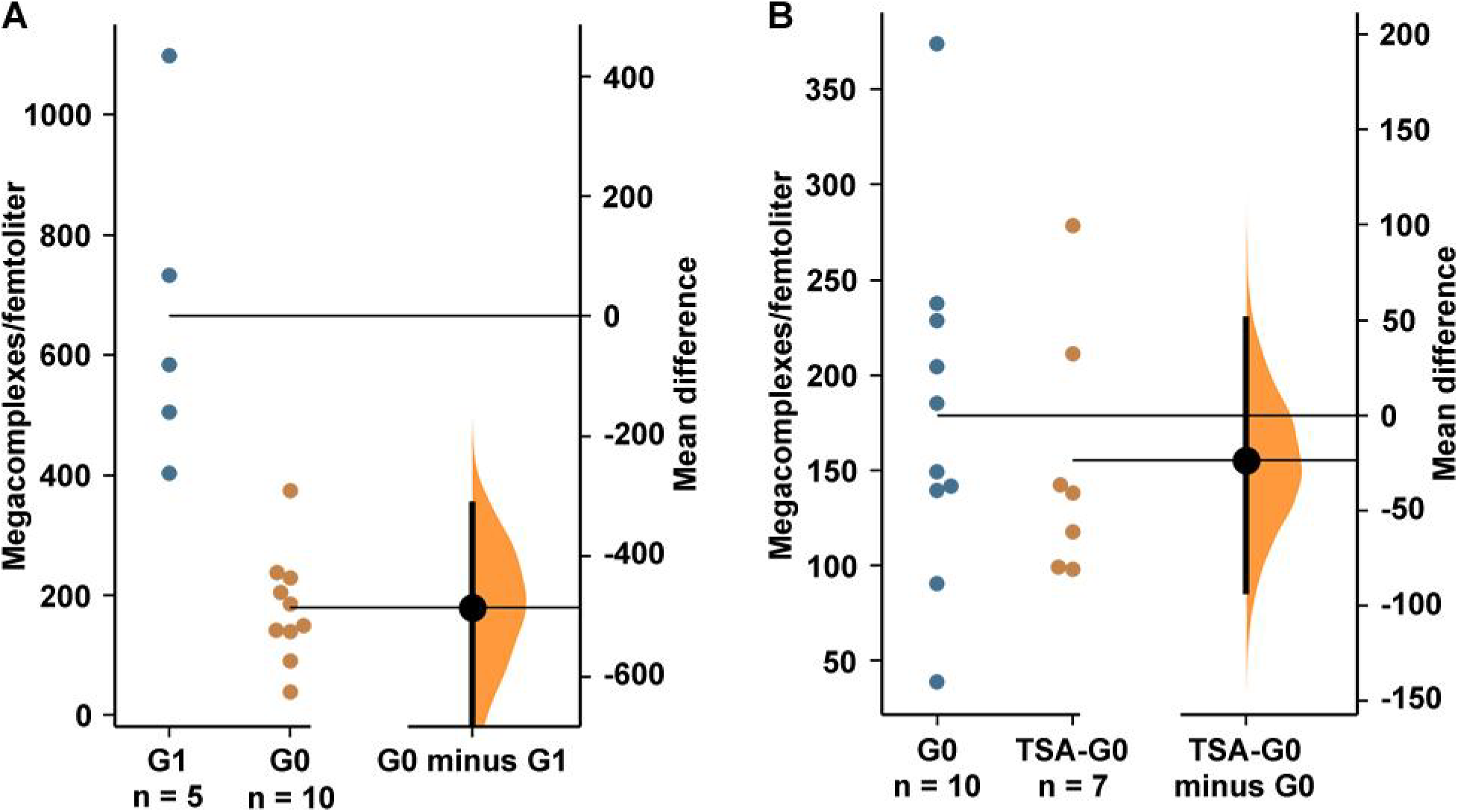
G0 and TSA-G0 cells have fewer megacomplexes in their nucleoplasm than G1. Gardner-Altman plots of (A) the mean difference between G1 and G0 megacomplex concentrations in the nucleoplasm, and (B) the mean difference between G0 and TSA-G0 megacomplex concentrations in the nucleoplasm. In each plot, both groups of samples are plotted on the left axes; the mean difference is plotted on a floating axis on the right as a bootstrap sampling distribution. The mean difference is depicted as a dot; the 95% confidence interval is indicated by the ends of the vertical error bar (Ho et al., 2019).

We next characterised the nucleolus, which can be immunofluorescently localised using the conserved marker fibrillarin (Tollervey et al., 1990). Fibrillarin immunofluorescence images showed that G0 cells had smaller nucleoli than G1 cells (**Figure 8, A and B**), in line with previous traditional EM data (Su et al., 1996). Some G1 and G0 cryotomograms seemingly included a portion of the nucleolus (**Figure 8, C and D**), in which megacomplexes were more abundant than in the rest of the nucleus (**Figure 8, E – H**). The fact that transcription is repressed in the G0 nucleolus even though megacomplexes are abundant there indicates that megacomplex concentration and transcription may be positively correlated in proliferating cells and uncorrelated in G0 vs G1 cells.

**Figure 8.**
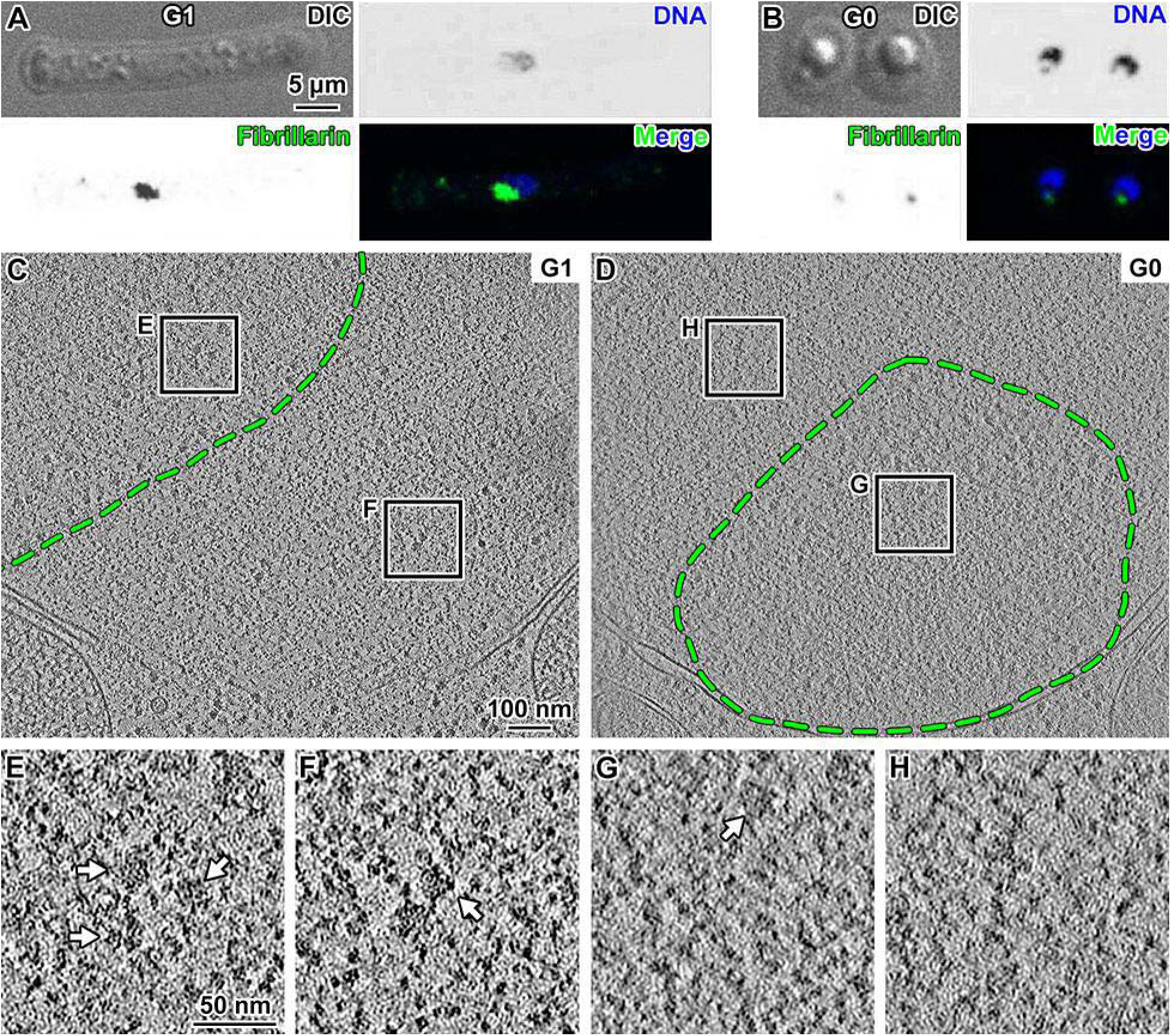
G0 nuclear megacomplexes are predominantly nucleolar. (A and B) Immunofluorescence detection of nucleoli in G1 and G0 cells. DAPI (blue) marks DNA and anti-fibrillarin antibody (green) marks the nucleolus. It is unknown what the origin of the small gap between the G0 cells’ fibrillarin and DNA signals is. (C and D) Volta cryotomographic slices (12 nm) of a G1 and a G0 cell, respectively. The region enclosed by the green dashed line denotes the approximate nucleolar boundary. (E and F) Four-fold enlargements of the G1 nucleolar and chromatin regions boxed in panel C. (G and H) Four-fold enlargements of the G0 nucleolar and chromatin regions boxed in panel G. Example megacomplexes are indicated by arrows in panels E, F, and G.

## DISCUSSION

When *S. pombe* cells enter G0, nearly every cell becomes shorter, undergoes transcriptional repression, and becomes more thermotolerant (Marguerat et al., 2012; Su et al., 1996). Several phenotype changes remained unknown, namely changes in histone acetylation, abundance of a key polymerase subunit, nucleus volume changes, distribution of nuclear macromolecular complexes, and structural state of the nucleosomes. Our study uses different microscopy and biochemistry techniques to fill in these knowledge gaps, giving a clearer picture of the cell-biological phenotypes of extreme cell states (**Figure 9**).

**Figure 9:**
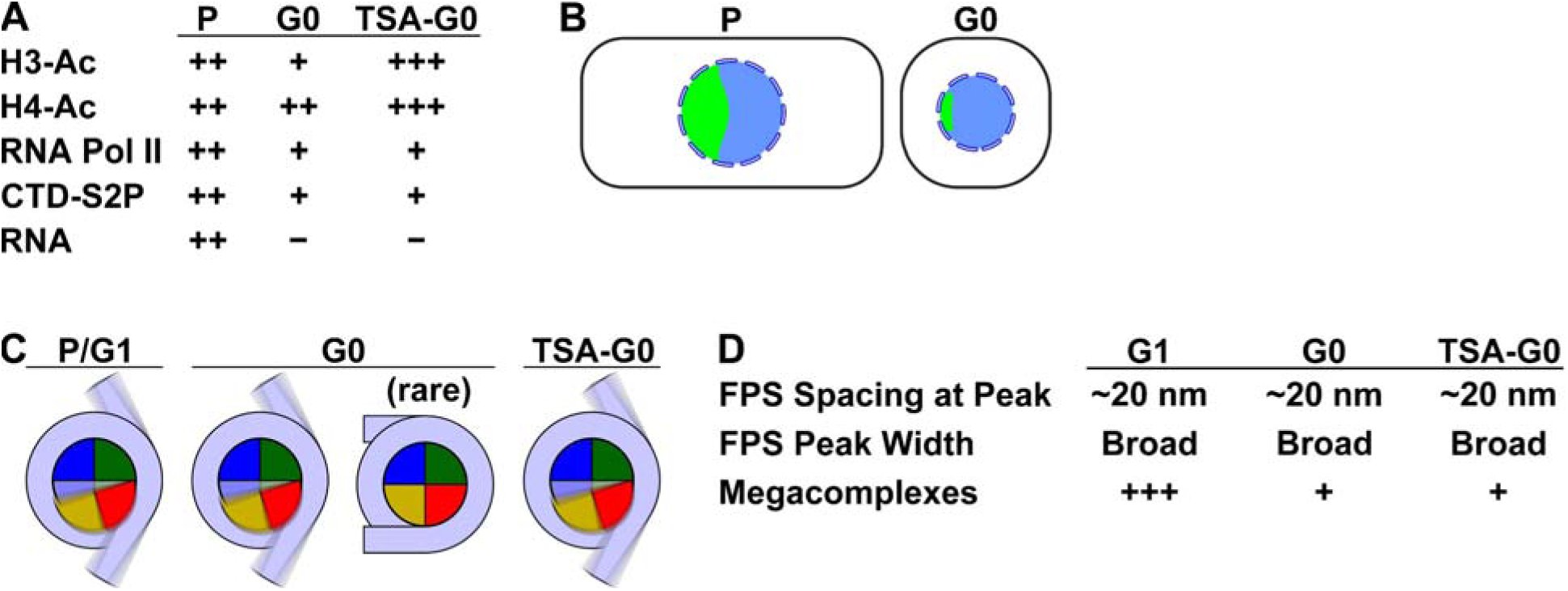
Summary of nuclear phenotypes in G0, G0-like, and proliferative cells. (A) Histone acetylation levels (H3-Ac and H4-Ac) are downregulated in G0 cells compared to proliferating (P) cells and upregulated in G0 cells treated with a histone-deacetylase inhibitor (TSA-G0). RNA Polymerase II levels and activity and transcription levels (RNA) are downregulated in G0 cells compared to proliferating cells and are uncorrelated with changes in G0 histone acetylation. (B) Cartoon of the cytological differences between proliferating (P) and G0 cells. Aside from having a smaller nucleus (blue circle), G0 cells have a disproportionately smaller nucleolus (green). (C) Schematics of DNA (light blue) and histones (shaded pie slices) in the nucleosome disc view. The cartoons only illustrate the 145 – 147 bp of “core” DNA. Majority of nucleosomes in G1, G0 and TSA-G0 are heterogeneous. The blurred appearance represents a large range of positions and orientations that protein and DNA components adopt inside cells, which would result in the absence of a class average resembling a canonical nucleosome; the blurring does not represent molecular motions. Nucleosomes with a structure resembling the canonical nucleosome are a minority conformation in G0, and even then, they have more unwrapped DNA than true canonical nucleosomes. (D) Fourier power spectrum (FPS) peak spacing and width are similar across G1, G0 and TSA-G0 cells. Megacomplex concentrations are higher in G1 cells than in G0 and TSA-G0 cells.

### *S. pombe* G0 transcription repression shares some features with other extreme states

Transcriptional repression has been studied in a few extreme cell states. Early studies found that chromatin compaction is correlated with near-complete transcriptional shutdown during mitosis (Prescott and Bender, 1962; Taylor, 1960). While mitotic chromatin is believed to sterically exclude transcription machinery (Luger et al., 2012; Maeshima et al., 2019), later studies showed that compacted mammalian mitotic chromatin is accessible enough to be bound by a small amount of the transcription machinery and that low levels of transcription or and even upregulation of some genes (Chen et al., 2005; Hsiung et al., 2015; Liang et al., 2015; Palozola et al., 2017; Teves et al., 2018; Teves et al., 2016). In line with previous studies (Marguerat et al., 2012; Su et al., 1996), we demonstrate here that *S. pombe* cell nuclei have lower active transcription in the G0 state than in the proliferative state. Notably, the lower 5-EU incorporation, lower RNA polymerase II levels, and lower levels of Rpb1 CTD-S2P and CTD-S5P markers seen here are all consistent with transcriptional repression.

We previously found that mitotic *S. pombe* chromatin contains megacomplexes and nucleosome-free “pockets”, indicating that the chromatin is unevenly packed (Cai et al., 2018b). This nuclear phenotype is conducive to mitotic transcription (Oliva et al., 2005; Peng et al., 2005; Rustici et al., 2004), which was supported by the observation of active Rpb1 CTD-S2P marks in mitotic chromatin (Cai et al., 2018b). While G0 transcription is on average repressed, there are some genes that expressed at levels similar to or greater than in proliferating cells (Marguerat et al., 2012; Shimanuki et al., 2007). Here we find that some of the cytological and biochemical phenotypes observed in mitotic *S. pombe* cells also hold for G0 cells. Megacomplexes are rarer within G0 chromatin, but abundant in the nucleolus. While the CTD-S2P signal is barely detectable in immunofluorescence images, our immunoblots show that the levels of CTD-S2P and CTD-S5P levels remain the same in G0 when calibrated to total Rpb1. We also find that G0 cells have non-trivial levels of Rpb1-eGFP in the nucleus. Therefore, *S. pombe* G0 cell nuclei also have properties conducive to low levels of transcription.

When *S. cerevisiae* exits G0, it undergoes a process termed hypertranscription, i.e., a large increase in transcription markers within minutes after G0 exit is triggered (Cucinotta et al., 2021). In contrast, *S. pombe* nascent transcripts (assessed by 5-EU incorporation) and RNA polymerase II activity (assessed by CTD-S2P and CTD-S5P marks) remain low for several hours after G0 exit is triggered. Interestingly, *S. cerevisiae* and *S. pombe* require 4 hours (Cucinotta et al., 2021) and 6 to 7 hours (Su et al., 1996), respectively, to re-start proliferation. In both organisms, re-entry into the proliferative state takes approximately 2 – 3 times longer than the cell cycle. It remains to be determined what physiological features account for the difference in G0-exit transcription programs.

### Histone hyperacetylation leads to complex phenotypes

Acetylation marks make chromatin more negatively charged and potentially change chromatin structure and function. For example, TSA treatment causes histone hyperacetylation and chromatin decompaction in both interphase HeLa (Toth et al., 2004) and G0 B cells (Kieffer-Kwon et al., 2017). In *S. cerevisiae* G0 cells, TSA treatment increases chromatin volume (Swygert et al., 2021) while deletion of the RPD3 lysine deacetylase gene (which also leads to increased histone acetylation) prevents transcriptional repression (McKnight et al., 2015). A recent study found that enzymes help control *S. pombe* G0 entry efficiency and survival (Zahedi et al., 2020). One may expect that perturbation of histone acetylation in *S. pombe* G0 cells may lead to large changes in cellular physiology. Compared to untreated G0 cells, TSA-treated G0 cells had hyperacetylated histone H3, longer cell bodies, and larger nuclei (similar volume to G1 nuclei). In contrast, TSA-induced histone hyperacetylation did not lead to a detectable decrease in thermotolerance or a derepression of transcription. Also, *S. pombe* TSA-G0 cell nuclei have similar macromolecular complex crowding as G0 cells, suggesting that the larger TSA-G0 nuclear volume may be packed with additional complexes, whose identities are unknown. A counter-intuitive histone hyperacetylation phenotype was also found in mitotic human fibroblasts (SK-N-SH cells), where TSA treatment does not inhibit progression through mitosis and leaves chromosomes compacted (Kruhlak et al., 2001). Changes in histone-acetylation therefore does not lead to predictable phenotype differences in cells with repressed transcription.

### G0 chromatin does not undergo extreme local compaction

In *S. cerevisiae*, the G0 nucleoplasm contains thicker STEM tomography densities than in interphase nucleoplasm, suggesting that its G0 chromatin is more compact (Swygert et al., 2021). Because *S. pombe* G0 nuclei have half the volume of G1 cells, we expected that that the G0 nuclear contents would also show major rearrangements, such as chromatin compacted into the domains in human perinuclear heterochromatin or the chromatids in mitotic human cells (Cai et al., 2018a; Chen et al., 2023). However, we did not see any regions in the G0 nuclei that resemble domains. Instead, our Fourier analysis showed that proliferating, G1, G0, and TSA-G0 nuclei have a similar crowdedness of macromolecular complexes, indicated by a single broad peak centred at approximately 20 nm. A simple explanation of the similar cryo-ET power spectra between different-sized nuclei is that the smaller G0 nuclei have fewer non-nucleosome nuclear complexes. These studies show that transcriptional repression may be achieved in diverse species and cell types that have different chromatin organisation.

### Canonical nucleosomes are rare in *S. pombe*

We recently found that in proliferating *S. cerevisiae*, less than 10% of the nucleosomes have the canonical structure (Tan et al., 2023). Here, we applied the same *in situ* 3-D classification analysis to *S. pombe* cryotomograms and found one class average in G0 chromatin that resembles the canonical nucleosome, accounting for <4% of the expected nucleosome population. This average had jagged density features, which suggests that misclassified complexes contributed heterogeneity that could not be removed by classification. Therefore, G0 canonical nucleosomes are likely rarer than 4%. No canonical nucleosome class averages were found in the proliferating, G1, and TSA-G0 cells, meaning that canonical nucleosomes are even rarer in these cell types. Note that these classification results mean that canonical nucleosomes are rare, not absent. The rarity of canonical nucleosomes is consistent with earlier findings that *S. pombe* nucleosomes have lower thermal stability and more flexible DNA ends than human nucleosomes *in vitro* (Koyama et al., 2017).

What intracellular factors result in the rarity of canonical nucleosomes inside *S. cerevisiae* and *S. pombe* cells? Canonical nucleosomes are rare in G0 cells even though all three RNA polymerases are repressed (Marguerat et al., 2012), meaning that transcriptional repression does not explain the rarity of canonical nucleosomes in *S. pombe*. Furthermore, the differences in histone acetylation levels in G0, G1, proliferating, and TSA-G0 cells, do not adequately account for the rarity of canonical nucleosomes *in situ*. Further comparative analysis of proliferating and G0 *S. pombe* cells may shortlist the factors that *do not* influence *in situ* canonical nucleosome abundance.

## MATERIALS AND METHODS

### Cell culture and synchronisation

Proliferating wild-type 972 h-cells (strain MBY99) were grown in Edinburgh Minimal Medium (EMM, USBio, Salem, MA) or in yeast extract with supplements (YES; 5 g/L^−1^ yeast extract, 30 g L^−1^ glucose, 75 mg L^−1^ histidine, 75 mg L^−1^ leucine, 75 mg L^−1^ adenine, 75 mg L^−1^ uracil) at 30°C (shaken at 200 – 250 RPM for all cell culture experiments) to OD_600_ = 0.2. Most cell washes were done by pelleting the cells by centrifugation, followed by resuspension in fresh growth media or buffer. To prepare G0 cells, proliferating cells were washed twice in EMM without nitrogen (EMM−N) and then incubated for 24 hours in EMM−N at 30°C. For experiments with *cdc10-129* (strain MBY165; *leu1-32*) cells, the cultures were grown in YES at 30°C. When the OD_600_ reached 0.2, the cultures were transferred to 36°C and incubated for 4 hours, which arrests the majority of cells in G1 phase. For G0 experiments, wild-type cells, yFS240 cells, or cells of strains that express either eGFP-tagged RNA polymerase II or eGFP-tagged nuclear pore subunits were used. *Cdc10-129* cells were not used for G0-cell analysis because after incubation in EMM−N (with leucine, which this strain requires for survival), they did not become shorter and were less thermotolerant than G0 cells of other strains. We suspect this phenotype is a consequence of the supplements, which may act as a sufficient source of nitrogen that prevents G0 entry.

### TSA perturbation

TSA (T8552, Sigma, Merck KGaA, Darmstadt, Germany) was prepared as a 20 mg mL^−1^ stock solution in dimethyl sulfoxide (DMSO). Cells growing in YES or EMM−N were pelleted and then TSA was added following two different regimens. For G0-TSA, proliferating cells were first incubated 24 hours in EMM−N, then 20 mg mL^−1^ TSA was added to 20 µg mL^−1^ (66 µM) final concentration. These cells were then incubated in TSA for 24 hours. For TSA-G0, proliferating cells were resuspended in EMM−N, then 20 mg mL^−1^ TSA was immediately added to 20 µg mL^−1^ final concentration. These cells were incubated for 24 hours. For the washout control, TSA-G0 cells were washed with fresh EMM−N, then incubated 24 hours before immunoblot processing.

### DAPI staining (not for immunofluorescence)

For microscopy of DAPI-stained nuclei, cells were grown in EMM because the DAPI signal is harder to see in YES-grown cells. Proliferating cells (0.5 OD_600_ units) were concentrated by pelleting at 5,000 × g for 1 minute and then resuspended in 1 mL EMM. G0 cell cultures started at OD_600_ = 0.2 and sometimes grew to OD_600_ > 0.5 after 24 hours of nitrogen starvation. These cultures were diluted to OD_600_ = 0.5 with EMM−N. Cell cultures were fixed by adding 37% formaldehyde (#47608-1L-F, Sigma) to 3.7% final concentration, incubated for 90 minutes (all fixation was done 200 – 250 RPM shaking), then collected by centrifugation at 5,000 × g for 1 minute. Cells were then washed twice in phosphate-buffered saline, pH 7.4 (PBS; Vivantis, Selangor Darul Ehsan, Malaysia) and resuspended in 20 µL of PBS with 1 µg mL^−1^ DAPI for G0 cells and 10 ng mL^−1^ DAPI for proliferating cells. To visualise the nuclei of G0 cells, they needed to be treated with 100-fold higher DAPI concentrations than proliferating cells.

Five μl of this sample was then added to a microscope slide and imaged using an Olympus FV3000 Confocal Laser Scanning Microscope (Olympus, Tokyo, Japan) equipped with a 1.35 NA 60× oil-immersion objective lens. The confocal microscopy details are listed in **Table S2**.

### Thermotolerance tests

Cells (OD_600_ range = 0.2 – 1.0 for proliferating, 0.7 for G0, and 0.5 for TSA-G0 – all at 0.01 OD_600_ units per sample) were pelleted at 5,000 × g for 1 minute. The cell pellets were resuspended in 100 µL of 30°C or 48°C YES or EMM−N medium (from 1 mL prewarmed stock) and then immediately placed in either a 30°C shaking incubator or a 48°C heating block, respectively, then incubated for 30 minutes. The 48°C-heated cells were then cooled on ice for 1 minute. Four serial dilutions (10×, 100×, 1,000×, 10,000×) were made for each sample into YES to 45 µL or 50 µL final total volume. The undiluted stock and diluted cultures (5 µL each) were spotted on a dry YES agar plate (2% w/v agar in YES medium). Once the cells were adsorbed onto the agar (∼30 minutes), the plates were turned upside down, sealed with Parafilm®, incubated at 30°C for 2 days, and then photographed.

### Immunoblots

Immunoblot samples were prepared using trichloroacetic acid (TCA) precipitation. The cell pellet (7 – 21 OD_600_ units, subsequently diluted to obtain equal protein loading) was resuspended in 200 µL of 20% TCA on ice. Approximately 0.4 g of glass beads (425 – 600 µm, Sigma) was added to the mixture. The cells were then vortexed for 1 minute, followed by incubation on ice for 1 minute; this vortex-incubation treatment was done four times in total. The cell lysate was then centrifuged at 2,000 × g for 10 seconds to sediment the glass beads. Next, 500 µL ice-cold 5% TCA was mixed with the lysate. The mixture (without glass beads) was transferred to a new tube. Another 500 µL of 5% ice-cold TCA was mixed with the glass beads and this new mixture (without glass beads) was added to the tube in the previous step. The combined mixture was then placed on ice for 10 minutes to precipitate proteins. Then the precipitated proteins were pelleted at 4°C, 15,000 × g for 20 minutes. The supernatant was removed and the pellet was re-centrifuged (either short-spin for 3 seconds or 15,000 × g for 1 minute), followed by the removal of the residual supernatant. The pellet was resuspended in 212 µL of 1× Laemmli sample buffer, followed by the immediate addition of 26 µL of 1 M Tris pH 8 to neutralise the residual TCA. Then the lysates were heated at 100°C for 5 minutes. The lysates were then centrifuged at 15,000 × g for 10 minutes and the clarified supernatant containing solubilized proteins was transferred to a new tube.

Protein loading levels were calibrated using immunoblots against both histone H3 and H4 C-termini, or against Rpb1 C-terminal domain. SDS-PAGE was done with Mini-PROTEAN® TGX™ 4-15% Precast Gels (Bio-Rad, Hercules, CA), electrophoresed for 90 minutes at 80 volts. Precision Plus Protein™ WesternC™ Standards (Bio-Rad) or Invitrogen™ MagicMark™ XP Western Protein Standard (Thermo Fisher Scientific, TFS, Waltham, MA) (2.5 µL) were loaded as molecular-weight markers. The proteins were then transferred to PVDF membrane (Bio-Rad Immun-Blot®) at 100 volts for 30 minutes at 4°C. The membrane was blocked in 2% BSA in 1× Tris-Buffered Saline, 0.1% Tween® 20 Detergent (TBST) for 1 hour at 22°C and then incubated with primary antibody overnight at 22°C. All antibody dilution factors are reported in **Table S3**. Next, the blot was rinsed in TBST for 20 minutes three times at 22°C, and incubated with HRP-conjugated secondary antibody in 2% BSA in TBST for 1 hour at 22°C, with additional StrepTactin-HRP Conjugate (Bio-Rad) if Precision Plus Protein™ WesternC™ Standards ladder was used. The blot was then rinsed in TBST for 10 minutes three times. Finally, the blot was treated with a 50:50 mixture of Clarity Western Peroxide Reagent and Clarity Western Luminol/Enhancer Reagent (Bio-Rad) for 5 minutes before visualisation. The chemiluminescent signals were recorded using an ImageQuant LAS 4000 (Cytiva, Marlborough, MA). Uncropped immunoblots are shown in **Figure S29**.

### Immunofluorescence microscopy

Indirect immunofluorescence microscopy was done using a modified version of our previous protocol (Cai et al., 2018b) as follows. Proliferating, G1, and G0 cells were fixed with 3.7% formaldehyde for 90 minutes at 30°C. Cells were collected by centrifugation at 5,000 × g for 5 minutes and were then resuspended in 1 mL PEM (100 mM PIPES, 1 mM EGTA, 1 mM MgSO_4_, pH 6.9) buffer. The cells were then washed once by centrifugation and resuspension with PEM, followed by resuspension in 1 mL PEMS (1.2 M sorbitol in PEM). Next, the cells were converted to spheroplasts with 25 U/mL Zymolyase (Zymo Research, Irvine, CA) in PEMS and incubated for 75 minutes in a 37°C water bath. All subsequent incubations were done at 10 RPM in a rotator (Invitrogen HulaMixer™). The cells were washed again with PEMS, resuspended in PEMS with 1% Triton X-100 (Sigma) and incubated at 22°C for 5 minutes. Cells were then washed twice with PEM and then incubated in PEMBAL (PEM, 1% bovine serum albumin (BSA), 100 mM l-lysine hydrochloride) for 1 hour at 22°C. The primary and secondary antibodies used for immunostaining are listed, alongside the immunoblot experimental parameters, in **Table S3**. The cells were resuspended in 100 µL of primary antibody diluted in PEMBAL, incubated at 22°C overnight. Next, the cells were washed twice with PEMBAL and resuspended in 100 µL of secondary antibody diluted in PEMBAL, and then incubated at 22°C in the dark overnight. Finally, the cells were washed in PEM, then PBS, then resuspended in 20 µL of DAPI diluted to 1 µg/mL in PBS. Five μl of the sample was then added to a microscope slide and imaged using either an Ultraview Vox spinning-disc confocal microscope (PerkinElmer, Waltham, MA) or an Olympus FV3000 Confocal Laser Scanning Microscope. The images were recorded using either a 60× or a 100× objective lens.

### Nascent RNA detection

Newly synthesised RNA was detected *in situ* using a 5-EU click-chemistry kit (“Click-iT™” C10329, Thermo Fisher Scientific (TFS), Waltham, MA). As a negative control, the RNA-polymerase inhibitor phenanthroline (20 mg mL^−1^ stock in deionized water, P9375, Sigma) was added to 1 mL cell culture to a final concentration of 350 µg/mL. Five-ethynyl-uridine (100 mM stock in deionized water) was added to the cultures of strain yFS240 or to strain MBY99 (the control strain) to 1 mM final concentration and incubated at 30°C in the dark with shaking at 250 RPM for the various durations indicated in the results. Cells were then fixed with 1 mL of 3.7% formaldehyde in PBS for 15 minutes at 22°C; all subsequent incubations were done at 10 RPM in a rotator. Next, the cells were washed once by pelleting at 1,500 × g and resuspending in 1 mL of PBS. The cells were permeabilized with 1 mL of 0.5% Triton X-100 in PBS for 15 minutes at 22°C in the dark and then washed in PBS. Then the cells were treated with a reaction cocktail made with 428 µL of Click-iT RNA reaction buffer, 20 µL of 100 mM CuSO_4_, 1.8 µL of Alexa Fluor® 488 stock solution and 50 µL of Click-iT reaction buffer additive as directed in the Click-iT RNA Alexa Fluor 488 Imaging Kit. This mixture was incubated at 22°C in the dark for 30 minutes. Finally, the labelled cells were washed in Click-iT reaction rinse buffer before they were resuspended in 20 µL of PBS with 1 µg/mL of DAPI.

### Plasmid extraction and linearization

The plasmid pFA6a-link-yoEGFP-Kan was a gift from Wendell Lim & Kurt Thorn (Addgene plasmid # 44900 ; https://www.addgene.org/44900 ; RRID:Addgene_44900) (Lee et al., 2013), given in the form of a DH5-Alpha *Escherichia coli* bacterial stab. Five mL of bacteria were cultured in vent cap tubes shaking at 220 RPM at 37°C overnight, then plasmids were extracted with the QIAprep Spin Miniprep Kit (QIAGEN, Hilden, Germany) following the manufacturer’s instructions.

Extracted plasmids were linearized by digestion with a reaction containing 1 µg of plasmid DNA, 5 µL of 10× rCutSmart™ Buffer (New England BioLabs, Ipswich, MA), 10 units of restriction enzyme (New England BioLabs) topped up to 50 µL with nuclease-free water. *Sal*I restriction enzyme was used for the template pFA6a-link-yoEGFP-Kan plasmid, *Aat*II was used for confirmation of the Gibson Assembly product plasmid (see below). The reaction mixture was heated at 37°C for 15 minutes to digest the DNA, then 80°C for 20 minutes to inactivate the enzymes.

### Strain construction

The strain details are shown in **Table S1**. Primers were from IDT (Integrated DNA Technologies, Inc., Singapore) and listed in **Table S4**. Q5 PCR Master Mix (New England BioLabs) was used for PCR reactions. MBY99 served as the wild-type strain. The insert and backbone fragments were amplified from pFA6a-link-yoEGFP-Kan plasmids linearized with *Sal*I restriction enzyme. Homology fragments were amplified from genomic DNA extracted from MBY99 with the DNeasy Blood & Tissue Kit (QIAGEN) following the manufacturer’s instructions. All PCR reactions contained 1 ng of plasmid template DNA or 100 ng of genomic template DNA plus 0.5 μM of each primer. The PCR program was 98°C for 30 seconds, then 30 cycles of 98°C for 5 seconds, 60°C for 10 seconds and 72°C for 1.5 minutes, then 72°C for 5 minutes.

For *rpb1-EGFP*, a plasmid containing the tagging cassette was created first by Gibson Assembly (Gibson et al., 2009). The fragment containing 1kb of homology to the *rpb1* gene sequence just before the stop codon (5’ homology fragment), the EGFP-Kan module fragment, the fragment containing 1kb of homology to the genomic sequence just downstream of the *rpb1* stop codon (3’ homology fragment), and the backbone fragment were amplified as described above. For each fragment, the reverse primer contained an overhang sequence matching the start of the next fragment. After their amplification, the backbone and 5’ homology fragments were combined by assembly PCR using the same PCR cycle as above, and the same was done with the EGFP-Kan module and 3’ homology fragments. The resulting assembly PCR fragments were purified with the QIAquick PCR Purification Kit (QIAGEN), then assembled into plasmids and cloned into DH5-Alpha *E. coli* with the Gibson Assembly® Cloning Kit (NEB) following the manufacturer’s instructions. The tagging cassette (5’ homology, EGFP-Kan module, 3’ homology) for transformation was then created by extracting the plasmids with the QIAprep Spin Miniprep Kit (QIAGEN) and linearizing the plasmids with *Aat*II restriction enzyme (New England BioLabs) using the digestion reaction as described above. The entire tagging cassette was then amplified using the same PCR cycle as above.

For *nup97-EGFP*, the tagging cassette was created directly through assembly PCR. The *nup97* 5’ homology fragment, EGFP-Kan module fragment, and *nup97* 3’ homology fragment were created as described above, each with the reverse primer containing an overhang sequence homologous to the start of the next fragment. All three fragments were then combined by assembly PCR using the same PCR cycle as above, and the resulting product was directly used for transformation.

Cells were transformed using the lithium acetate / PEG4000 method reported in (Murray et al., 2016) with modifications. Overnight cell culture in YES was diluted to OD_600_ = 0.2 in 25 mL of YES and grown to an OD_600_ of 0.5. Ten mL of cells were collected, centrifuged at 2,500 × g at 22°C for 5 minutes, and the supernatant was removed. The cells were then washed with 5 mL sterile water with centrifugation at 2,500 × g at 22°C for 3 minutes. The cells were resuspended in 1 mL sterile water, transferred to a new 1.5 mL collection tube, centrifuged at 16,000 × g at 22°C for 1 minute, washed in 1 mL of TE/LiAc (10 mM tris, 1 mM ethylenediaminetetraacetic acid (EDTA), 100 mM lithium acetate), with centrifugation at 16,000 × g for 1 minute, and resuspended in 100 μL of the same buffer. One hundred μL of cell suspension was transferred to a new 1.5 mL collection tube containing 10 μL of 2 mg mL^−1^ salmon sperm DNA (Sigma-Aldrich, Burlington, MA) plus 5 μL of PCR-amplified cassette DNA, and incubated for at 22°C for 10 min. 260 µL of TE/LiAc/PEG (10 mM tris, 1 mM EDTA, 100 mM lithium acetate, 40% w/v Polyethylene glycol 4000) was added and incubated with shaking at 22°C for 30 minutes. Forty-three µL of DMSO was added, the suspension was incubated at 42°C for 15 minutes in a water bath, then the suspension was centrifuged at 6,000 x g for 1 minute. The supernatant was removed, the cells were resuspended in 1 mL of EMM−N, and the suspension was transferred to a vented-cap tube shaking at 220 RPM at 30°C for 24 hours. Finally, the cells were transferred to a new 1.5 mL collection tube, centrifuged at 16,000 × g for 1 minute, the medium was removed, and the cells were resuspended in 300 μL of 1× TE buffer, pH 8.0 (10 mM tris, 1 mM EDTA). All this cell suspension was plated on a G418 YES selection plate and incubated for several days at 30°C. G418 selection plates were created by adding G418 to the molten YES agar to a concentration of 200 mg L^−1^ before it was poured.

Newly created strains were authenticated by genomic DNA extraction with the DNeasy Blood & Tissue Kit (QIAGEN) following the manufacturer’s instructions, followed by PCR of integration junctions and Sanger sequencing. Confirmation PCR was performed with a program of 94°C for 2 minutes, 30 cycles of 94°C for 1 minute, 62°C for 1 minute and 72°C for 3.5 minutes, then 72°C for 5 minutes. The PCR products were electrophoresed in a gel containing 2% agarose and FloroSafe DNA Stain in Tris-acetate-EDTA, at 100 Volts for 1 hour, then visualised in a G:Box (Syngene). For sequencing, overlapping PCR fragments were created with the above cycle, purified with the QIAquick PCR Purification Kit (QIAGEN) with water used for the final elution, before the fragments and the sequencing primers were sent for Sanger sequencing (Bio Basic Asia Pacific Pte Ltd, Singapore).

### Analysis of cells exiting G0

Wild-type, yFS240, and Rpb1-eGFP cells were prepared as G0 cultures as described above. For the 0-minute G0 exit, 1 mL of wild-type cells was directly subjected to the immunofluorescence-microscopy procedure above; 1 mL of yFS240 cells was directly subjected to the nascent-RNA-detection procedure above; 1 mL of Rpb1-eGFP cells was directly imaged by confocal fluorescence microscopy; and 25 mL of wild-type cells was pelleted in preparation for the immunoblot procedure above, all without changing the culture medium. For 30 minute, 2 hour, 4 hour, and 6 hour G0 exit, either 1 mL or 25 mL of G0 cells was pelleted, then resuspended in an equal volume of YES and incubated for the respective duration at 30°C before being subjected to the above procedures. For the 24-hour G0 exit, either 50 µL of cells was inoculated in 5 mL of YES, then incubated 24 hours at 30°C before 1 mL of this culture was directly imaged or subjected to the immunofluorescence-microscopy or nascent-RNA-detection procedures; or 250 µL of cells was inoculated in 25 mL of YES, then incubated 24 hours at 30°C before all 25 mL of this culture was pelleted in preparation for the immunoblot procedure. All RNA labelling with 5-EU was performed for 10 minutes so that, for example, the “30 minute G0 exit” sample was first incubated for 30 minutes in YES, then an additional 10 minutes in YES plus 5-EU, followed by fixation for treatment with the Click-iT RNA Alexa Fluor 488 Imaging Kit.

### Fluorescence microscopy of living cells

Cells were first grown to log phase (proliferating) or subjected to nitrogen starvation, with TSA treatment when needed (G0 and TSA-G0). A thin pad agar made from 2% w/v agar in YES (proliferating) or EMM−N (G0 and TSA-G0) medium was prepared and spread on a glass slide before 5 µL of cell culture was applied to the coverslip and pressed against the agar pad, which was trimmed to the size of the coverslip. Samples were imaged with an Olympus FV3000 Confocal Laser Scanning Microscope using a 60× objective lens. Images were captured as Z-stacks thick enough to image the GFP signals from all of the nuclei at each stage position. To capture the moment the nuclei have just divided, time-lapse imaging was performed with intervals of 10 minutes for 3 hours for nucleus size measurement, and with intervals of 30 minutes for 19 hours for G0 exit. These time-lapse data were captured by collecting Z-stacks with Z-drift compensation at each interval. Time-lapse imaging was not performed on most G0 and TSA-G0 cells for nucleus size measurement because time-lapse-imaging tests did not detect any changes in nucleus morphology in these cells during the 3 hours of time-lapse imaging.

### Nucleus volume measurement

Z-stacks of Nup97-eGFP cells were analysed in FIJI (Schindelin et al., 2012). To facilitate the estimation of nuclear volume, spherical nuclei were chosen for analysis; elongated and irregularly shaped nuclei were excluded. For each nucleus in the Z-stack, the Z-slice where the nucleus appears the largest was chosen. The oval selection tool was used to select a circular area extending from the centre of the Nup97-eGFP (green channel) signal from one end of the nucleus to the centre of the Nup97-eGFP signal on the opposite end, after which the diameter of the circle was recorded. For proliferating cells, a time-lapse Z-stack was analysed. Nuclei that had just divided were identified as G1 nuclei for nucleus volume analysis as described above. After all nucleus diameters were recorded, their volumes were calculated from the sphere volume formula 4/3 × π × (diameter / 2)^3^.

### Cryo FIB milling

Proliferating, G1-arrested, G0 and TSA-G0 cells were first plunge-frozen with a Vitrobot, Mark IV (TFS) operated at the respective growth or arrest temperatures (37°C for G1-arrested cells, 30°C for proliferating, G0 and TSA-G0 cells) with 100% relative humidity. Cells were centrifuged at 5,000 × g, the supernatant was removed, then the cells were resuspended in a small volume of the same medium such that the new OD_600_ was 2.5 for proliferating and G1-arrested cells, and 4.0 for G0 and TSA-G0 cells. Four μl of cells at OD_600_ = 2.5 or 4.0 was pipetted onto the carbon side of an electron microscopy grid, manually blotted from the other side of the grid with a filter paper (Whatman, Grade 1) for 1 – 2 seconds, then plunged into 37% ethane/63% propane cooled by liquid nitrogen (Tivol et al., 2008).

Grids were milled with a FEI Helios dual-beam electron microscope following the protocol of Medeiros and Böck (Medeiros et al., 2018). The samples were inserted into the microscope at −158°C using a PolarPrep 2000 Cryo Transfer System (Quorum Technologies, Laughton, UK), then coated with organoplatinum with a gas injection system. The organoplatinum was warmed to 33.85°C and then “Flow” in the xT microscope control GUI was set to “Open” for 15 or 20 seconds, as in the “cold-deposition” method (Hayles et al., 2007). Cells were located on the grid, then milled to generate a 100 to 150 nm thick, 15 µm wide lamella near the centre of a cell, starting with 2.8 nA beam current rough milling, followed by 0.46 nA intermediate milling, and ending with 48 pA for the final polishing step.

### Electron cryotomography

Tilt series of cryolamellae were collected on a Titan Krios (TFS) operated at 300 kV, and equipped with a VPP (TFS), a K3 direct detection camera (AMETEK, Berwyn, PA), and a Gatan imaging filter (AMETEK) operated in zero-loss mode with a slit width of 20 eV. Detailed imaging parameters for lamellae are shown in **Table S5**. Images were captured as super-resolution movie frames at a 1.7 Å pixel size, using SerialEM (Mastronarde, 2005). Movies were aligned with IMOD alignframes (Mastronarde, 1997). The tilt series were automatically coarse aligned using the IMOD program *Etomo* running in batch-tomography mode (Kremer et al., 1996; Mastronarde, 1997; Mastronarde and Held, 2017). Tilt series alignment was done in eTomo by patch tracking.

Contrast-transfer function compensation was done for the few defocus phase-contrast images, but not for the VPP images. To suppress the high-spatial-frequency noise, the tilt series were binned 4 times and low-pass filtered with Etomo 2-D filter parameters µ = 0.35, and σ = 0.05 for a 6.8 Å pixel size. Cryotomograms were reconstructed by weighted back projection using the default Etomo parameters, then trimmed in 3-D to exclude features outside the field of interest. The additional details of each analysed tilt series are in **Table S6**.

### Template matching

Template matching was done using the PEET package (Particle Estimation for Electron Tomography) (Heumann, 2023; Heumann et al., 2011; Nicastro et al., 2006). A rounded 5 nm radius, 6 nm thick cylinder was used as a template. This template was enclosed with a cylindrical mask of 6.1 nm height and 5.4 nm radius, smoothed by convolution with a Gaussian of standard deviation of 0.68 nm. A cubic search grid with a 11 nm spacing was generated with the PEET program *gridInit*. Because this grid extended into the cytoplasm, a nucleus-enclosing boundary model was created within the same grid model file. This boundary model was drawn along the nuclear envelope at the “top” and “bottom” of the cryotomogram. The search points outside the nucleus were excluded with the IMOD command:

clipmodel -bound 2 original.mod new.mod

Template-matching hits that were within 6 nm of each other were considered as duplicates. One of the duplicates was automatically removed.

### Classification analysis of nucleosome-like particles

All scripts prefixed by ot_ are available at https://github.com/anaphaze/ot-tools. Classification was done with RELION 3.0 (REgularised LIkelihood OptimisatioN) (Kimanius et al., 2016; Zivanov et al., 2018), using the subtomogram-analysis routines (Bharat et al., 2015). The nucleosome-like particles’ centres of mass were converted from a PEET .mod file to a text file with the IMOD program *model2point*. These positions were then read into RELION to extract subtomograms with a 24.4 nm box size.

Our previous studies showed that some canonical nucleosomes are lost during 2-D classification (Cai et al., 2018a; Tan et al., 2023), so we exclusively used direct 3-D classification here to maximise the number of detected canonical nucleosomes. A rounded 10 nm wide, 6 nm thick cylinder was used as a reference map, and a smooth cylinder of the same dimensions and a cosine-shaped edge was used as a reference mask, both created with *beditimg*, the latter with *relion_mask_create* used as well. All nucleosome-like particles identified in PEET were subjected to 3-D classification with 100 classes, an experimental image mask diameter of 120 Å, a resolution cutoff of 20 Å, and 4 translational search steps.

### Fourier power spectrum analysis

Tomographic slices were generated from 18 central slices, then imported into FIJI (Schindelin et al., 2012). A 800 × 800 pixel box of a region of the tomographic slice of the chromatin inside the nucleus (to exclude the nucleolar densities) was selected, then the Fourier Transform was generated using the FFT tool. The Radial Profile Angle plugin (Carl, 2020) was used to generate the rotationally averaged (1-D) power spectrum. The data was analysed and plotted with Excel (Microsoft Inc., Redmond, WA): the x axis has units of inverse nanometers according to the formula 1 / (2 × 0.68 nm) × (radius in pixels) / 512, where 1 / (2 × 0.68 nm) is the Nyquist limited resolution for data that has a pixel size of 0.68 nm.

To compare the nuclear power spectra from different cell types, we first normalised each spectrum by calculating the average of all amplitude values in the spectrum up to a resolution cutoff of 5 nm, then dividing each amplitude value by this average. After normalisation, the power spectra of all analysed tomograms of a given sample type were averaged at each spacing to produce the mean. The error bars represent the standard deviation of the amplitude values at each spacing.

### Megacomplex analysis

To facilitate megacomplex picking, cryotomograms were binned twofold in 3-D with the IMOD program *binvol*. Each cryotomogram was opened with the 3dmod *slicer* tool set to ∼30 nm thick tomographic slices. Nuclear particles that were (1) larger than 15 nm and (2) outside the nucleolus were manually picked as “scattered” contour points.

To estimate the chromatin volume, a “closed” contour was drawn around the chromatin position at the central slice of the cryotomogram. The area within this contour was calculated using the IMOD command:

imodinfo -F model.mod

This method reports the contour’s “Cylinder Volume”, defined in this case as Area × pixel_size^3^, with Area expressed in voxels and pixel_size expressed in nanometers. This area was then multiplied by the tomogram thickness (in voxels) to estimate the volume.

## Abbreviations

G0: the quiescent state
CTD-S2P: RNA polymerase II large-subunit C-terminal domain phospho-serine 2
CTD-S5P: RNA polymerase II large-subunit C-terminal domain phospho-serine 5
5-EU: 5-ethynyl-uridine
TSA: Trichostatin A
cryo-ET: electron cryotomography / cryo-electron tomography
Megacomplex: multi-megadalton globular complexes

## Data sharing

One proliferating cell, G1 cell, G0 cell and TSA-G0 cell lamella cryotomogram each has been deposited as EMDB entry EMD-0875. All the tilt series and cryotomograms presented in this paper were deposited as EMPIAR (Iudin et al., 2016) entry EMPIAR-10339. We focused our efforts on the highest-contrast cryo-ET data that contained nuclear positions, leaving a large number of tilt series unanalysed. These surplus tilt-series and many corresponding cryotomograms will be deposited with our yeast surplus-data (Gan et al., 2019) entry EMPIAR-10227.

## Figure preparation and statistics

Image format interconversion and contrast adjustments were applied to the entire field of view using FIJI (Schindelin et al., 2012) or Adobe Photoshop (Adobe Systems, San Jose, CA). Student’s t-tests were done with Google sheets (Alphabet Inc., Mountain View, CA). Plots of the rotationally average power spectra were made with Excel (Microsoft Inc., Redmond, WA). Gardner-Altman estimation statistics plots (Ho et al., 2019) were made with the Estimation Statistics website (Claridge-Chang and Assam, 2023).

## Contributions and Notes

ZYT, SC - project design, experiments, analysis, writing; SAP, XN - project design, experiments; JS - training; LG - project design, experiments, analysis, writing, training.

The authors declare no conflicts of interest.

## ACKNOWLEDGEMENTS

We thank the CBIS microscopy staff for support and training. We thank Mohan Balasubramanian for strains MBY99 (972 h-) and MBY165 (*cdc10-129*); Makoto Ohira and Nick Rhind for sharing and advising on the use of the TK-hENT strain yFS240; Duane Loh for advice on Fourier analysis; and Claris Chong and Jon Chen for comments. This work was supported by a Singapore Ministry of Education T2 MOE2019-T2-2-045 and a T1 R-154-000-B42-114 grant.

---

**Figure S1.**
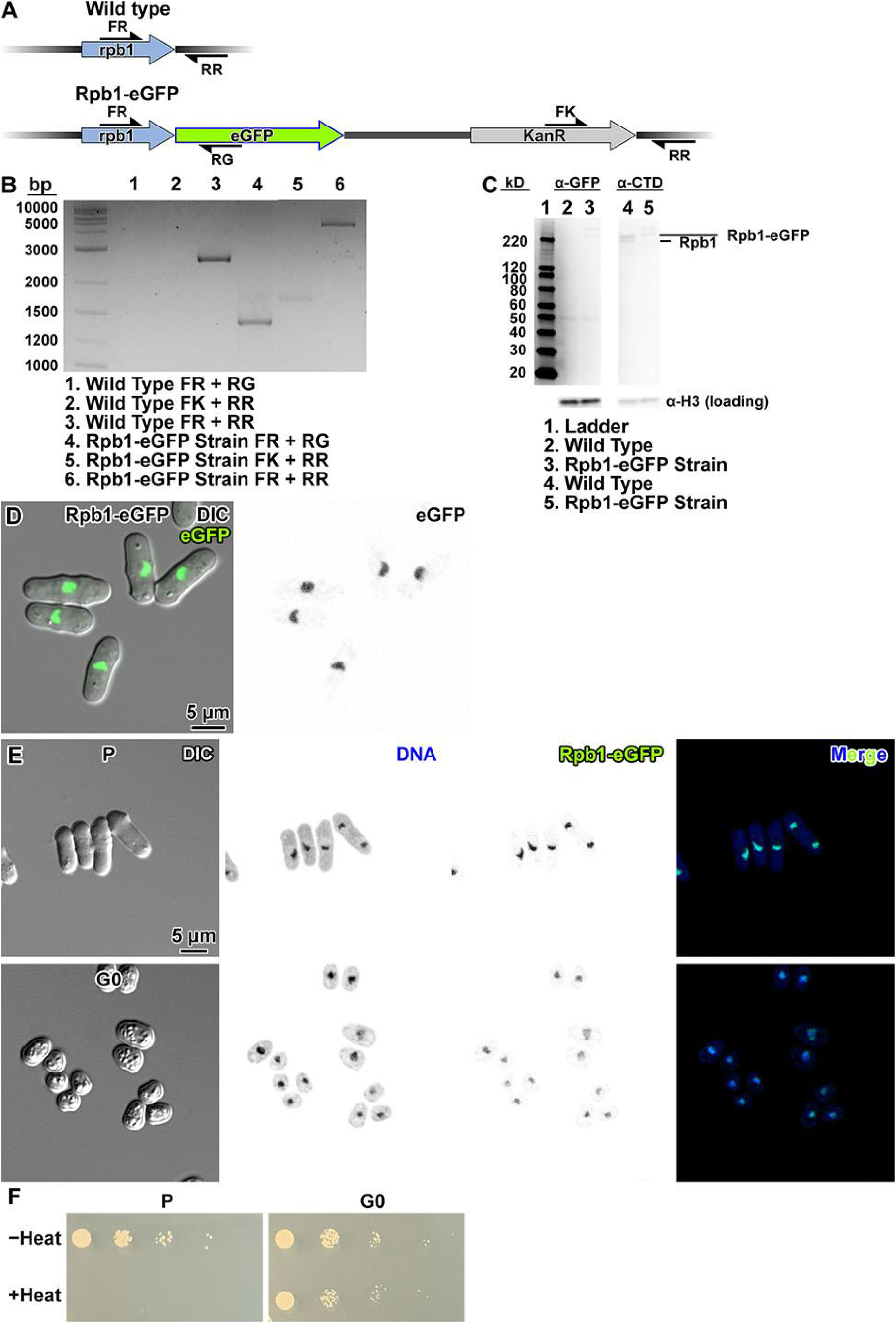
Construction of Rpb1-eGFP strain. (A) Map of the *rpb1* locus in the parent (wild-type) *S. pombe* strain MBY99 and the Rpb1-eGFP strain (LGSP0001). Primers used for PCR verification are indicated with the half arrow symbols. (B) Agarose gel of PCR amplicons expected from wild-type (control) and from Rpb1-eGFP genomic DNA, in which the *rpb1* locus is tagged with eGFP. (C) Immunoblot analysis of Rpb1-eGFP. The α-eGFP antibody correctly detected the Rpb1-eGFP fusion protein in the newly created strain (lane 3), but not in the wild type (lane 2, negative control) in the uncropped α-eGFP immunoblot. The α-CTD antibody detected the larger Rpb1-eGFP in the Rpb1-eGFP strain (lane 5) and the smaller unfused Rpb1 in the wild type (lane 4) in the α-CTD immunoblot. The H3 loading control is shown for each antibody because each antibody was imaged at a different exposure. Uncropped immunoblots are shown in Figure S29. (D) Differential interference contrast (DIC) and GFP fluorescence confocal microscopy images of Rpb1-eGFP cells. In the left panel, a merge of the DIC channel and the eGFP fluorescence channel is shown. In the right panel, eGFP signals are rendered with inverted contrast. (E) Fluorescence microscopy analysis of Rpb1-eGFP in interphase (P) and G0, in the Rpb1-eGFP strain. Columns, left to right: differential interference contrast (DIC), DAPI fluorescence (DNA) rendered in inverse contrast, eGFP fluorescence (Rpb1-eGFP) rendered in inverse contrast, and merge of the two fluorescence channels. Cells were fixed with formaldehyde. (F) Spot tests of proliferating (P) and G0 cells of the Rpb1-eGFP strain after a 30-minute incubation in YES and EMM−N respectively, with and without heat stress.

**Figure S2.**
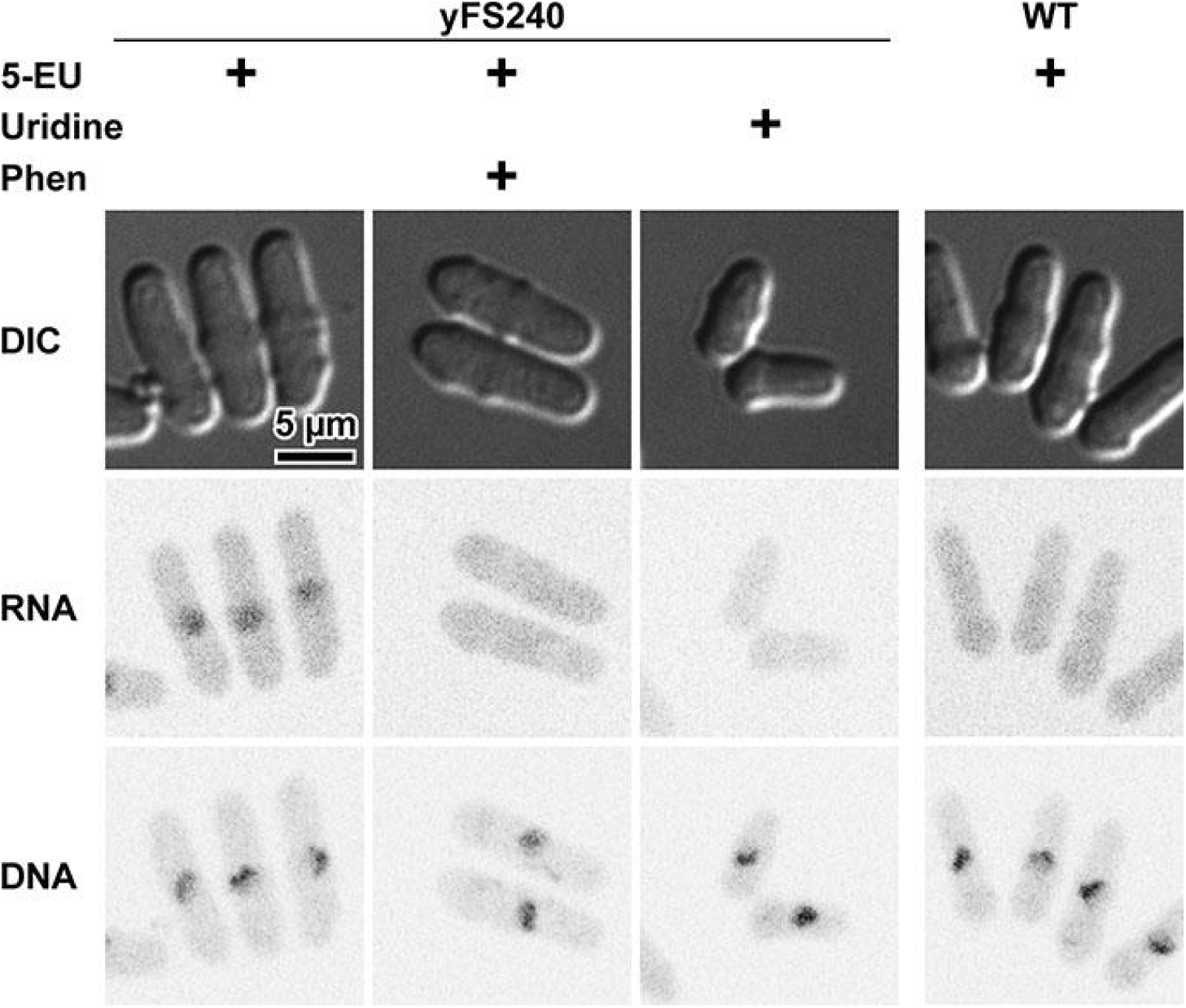
Controls for 5-EU RNA labelling. Proliferating yFS240 or wild-type cells incubated 10 minutes with 5-EU, uridine, or 5-EU plus the transcription inhibitor phenanthroline (Phen). The cells were then labelled with Alexa Fluor 488 and counterstained with DAPI.

**Figure S3.**
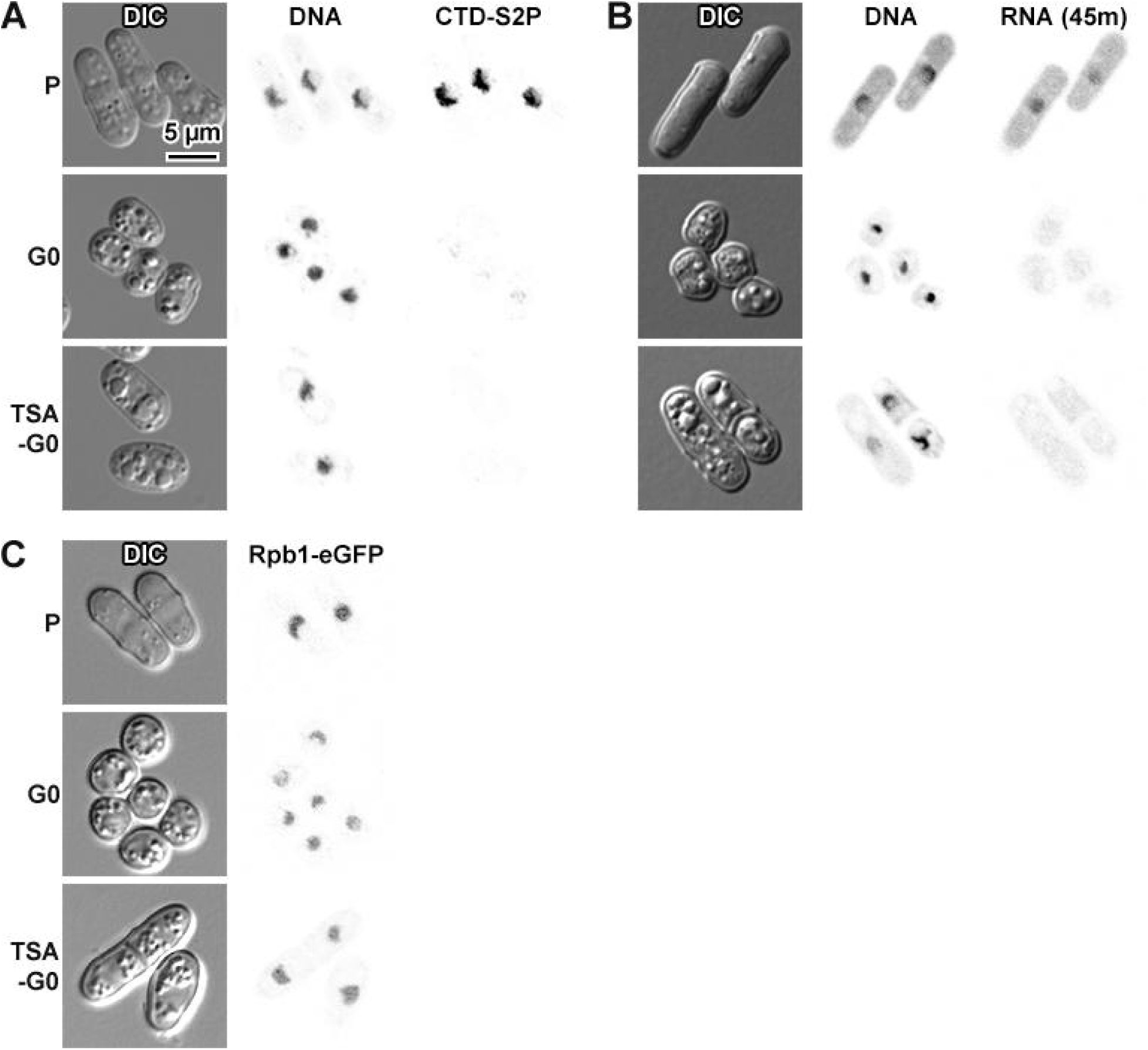
TSA treatment during G0 entry does not affect transcriptional repression. (A) DIC and fluorescence microscopy of proliferating (P), G0, and TSA-G0 wild-type cells. The cells were stained for DNA with DAPI and immunostained for RNA polymerase II with CTD-S2P. (B) DIC and fluorescence microscopy of proliferating (P), G0, and TSA-G0 yFS240 cells. The cells were incubated with 5-EU for 45 minutes to increase the amount of 5-EU incorporated into RNA. Nascent RNA was then ligated to Alexa Fluor 488 for fluorescence detection. (C) DIC and fluorescence microscopy of proliferating (P), G0 and TSA-G0 Rpb1-eGFP cells. The cells were unfixed and unstained; the fluorescence signal is from cell-expressed Rpb1-eGFP.

**Figure S4.**
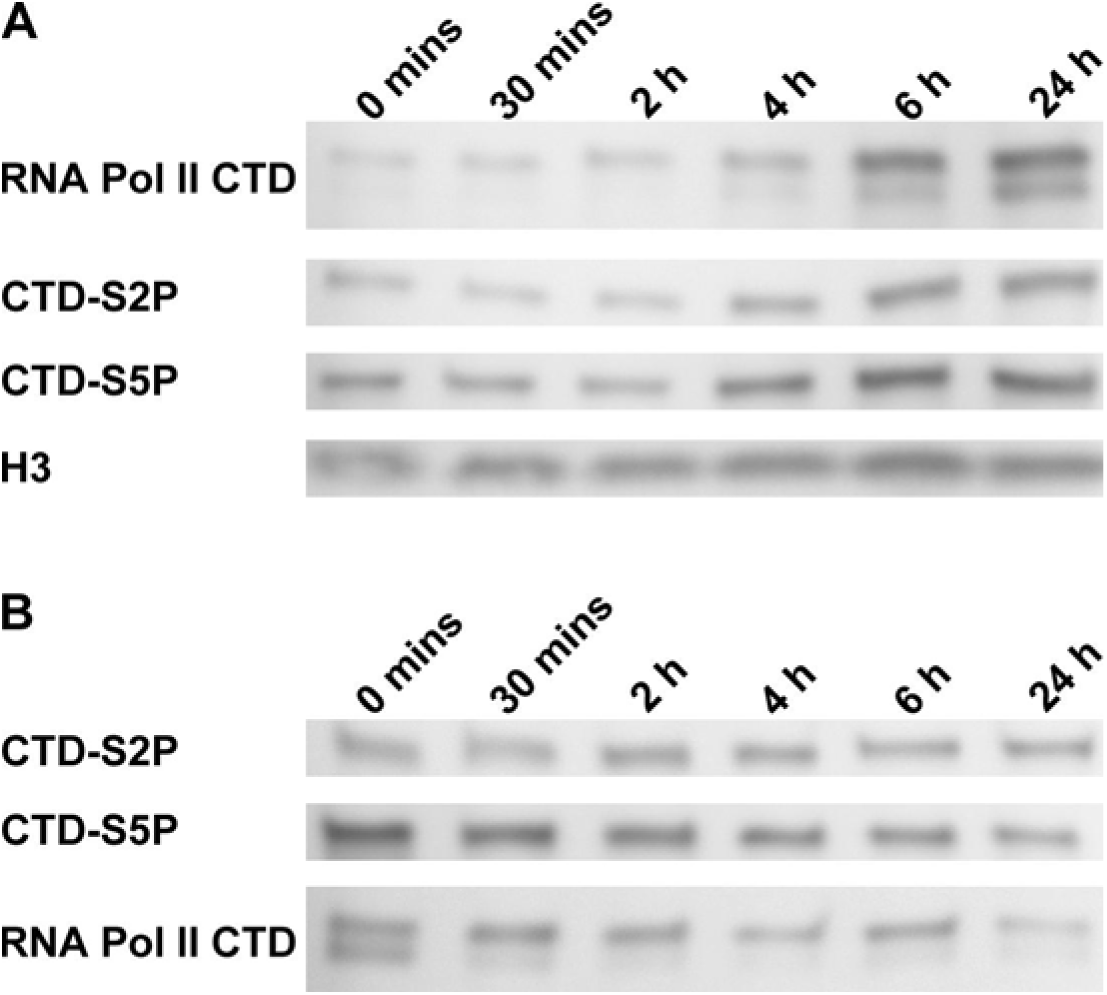
RNAPII-CTD transcription marker levels in whole-cell lysates do not markedly increase after G0 cells are switched to nutrient-rich medium. (A) Immunoblots of lysates from G0 wild type cells 0 minutes, 30 minutes, 2 hours, 4 hours, 6 hours or 24 hours after being transferred to YES medium, using a pan-CTD antibody, an α-CTD-S2P antibody and an α-CTD-S5P antibody. Loading controls were done with a H3 antibody (lower row). (B) Immunoblots of another batch of lysates from G0 wild type cells 0 minutes, 30 minutes, 2 hours, 4 hours, 6 hours or 24 hours after being transferred to YES medium, using an α-CTD-S2P antibody and an α-CTD-S5P antibody. Loading controls were done with a pan-CTD antibody (lower row). Uncropped immunoblots are shown in Figure S29.

**Figure S5.**
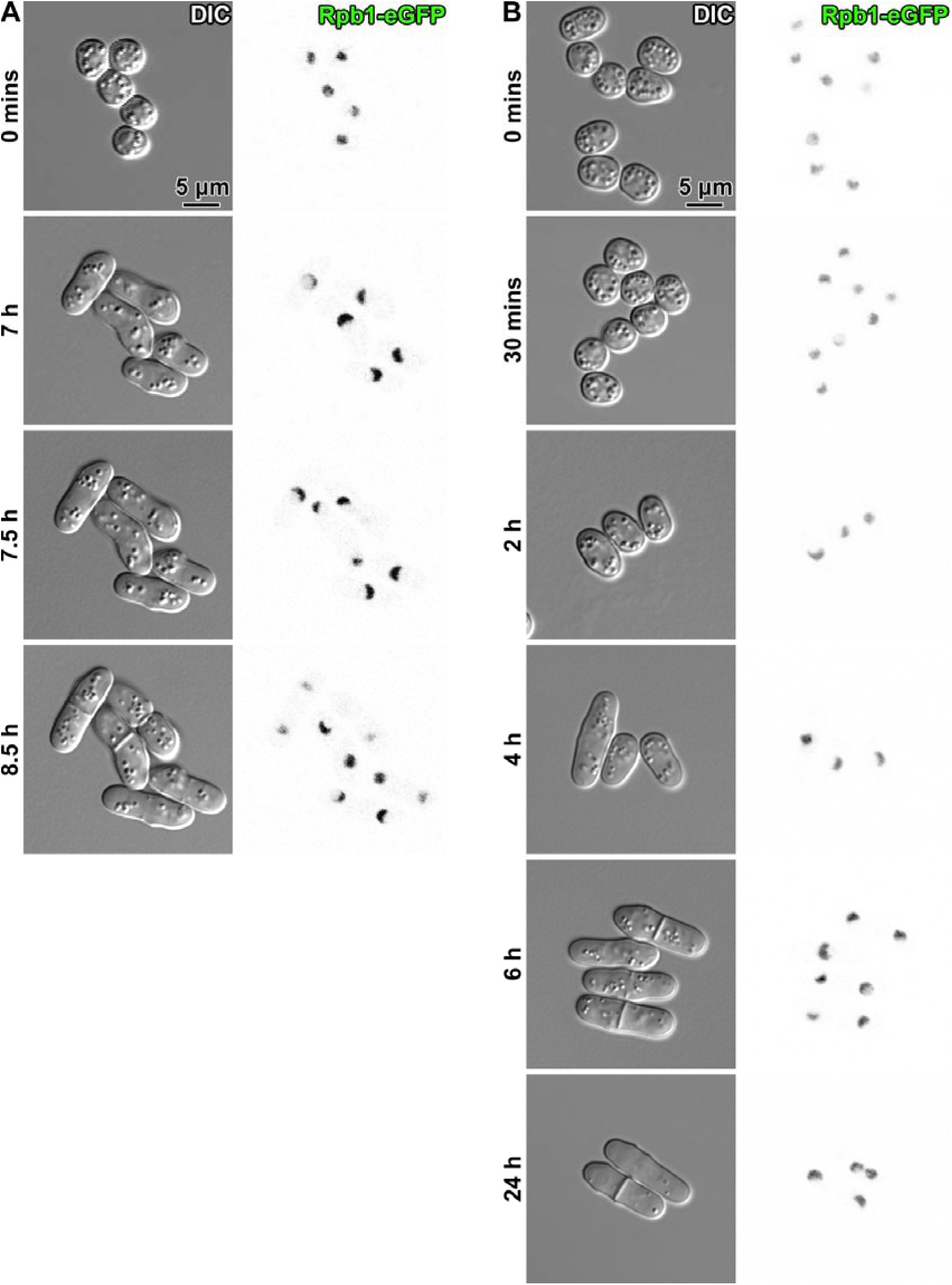
RNAPII does not exhibit observable changes within 6 hours after G0 cells are subjected to nutrient-rich medium. Fluorescence microscopy analysis of RNAPII content in cells after Rpb1-eGFP G0 cells are exposed to nutrient-rich medium. Columns, left to right: differential interference contrast (DIC) and eGFP fluorescence (Rpb1-eGFP). (A) Select samples from time-lapse images of a single group of cells imaged immediately after deposition on a YES agar pad (0 mins), just before (7 h), and just after (7.5 h) one of the cells finishes nuclear division, and just after all five of the cells finish nuclear division (8.5 h). See Movie S1 for the full set of time-lapse images. (B) Different groups of cells collected from liquid culture at different time points.

**Movie S1. Time-lapse fluorescence microscopy of Rpb1-eGFP cells exiting G0, deposited on an agar pad.**

The images on the left are differential interference contrast (DIC), the images on the right are eGFP fluorescence (Rpb1-eGFP). The interval between each frame is 30 minutes.

**Figure S6.**
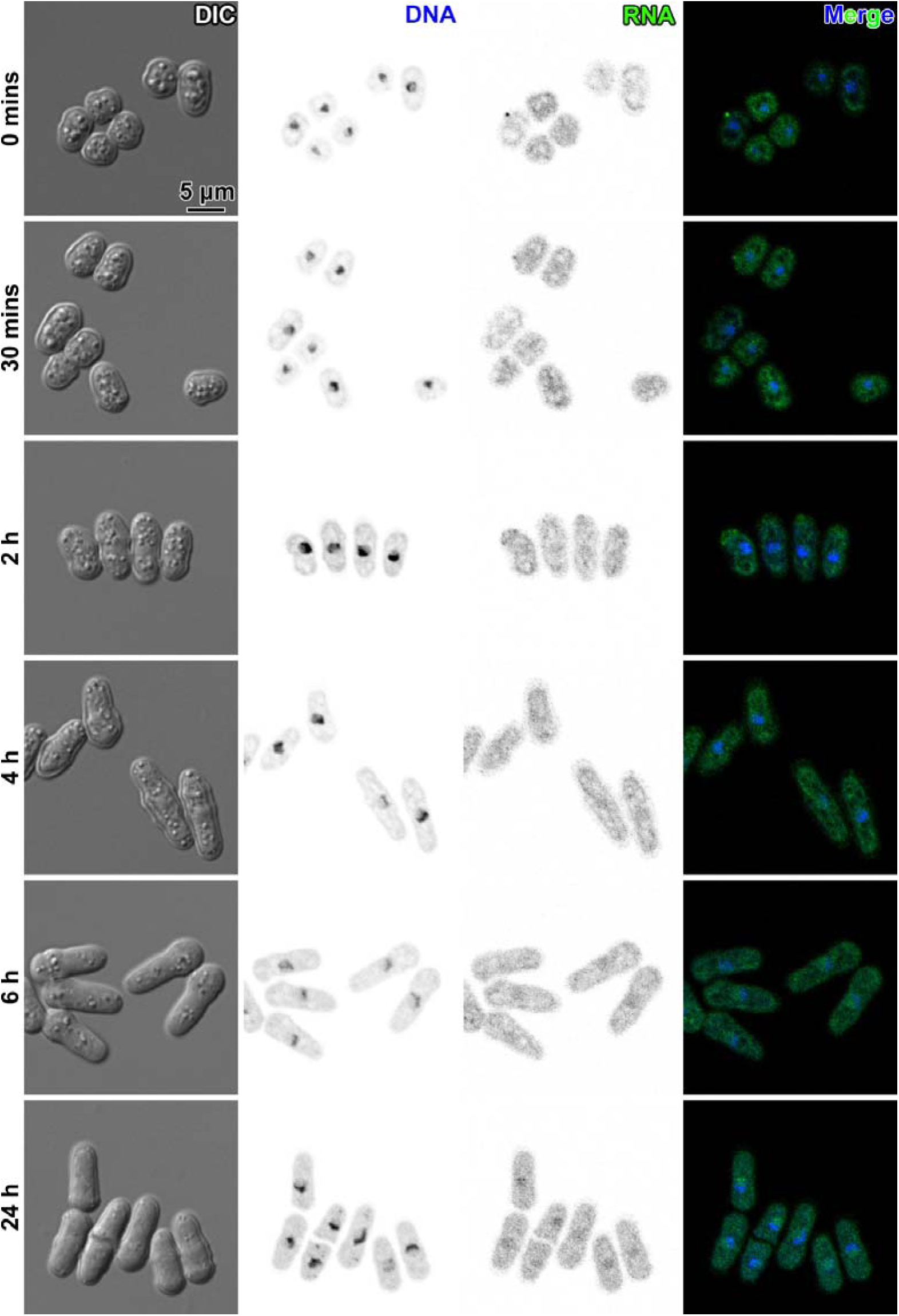
Nascent RNA transcripts are observable 6 hours after G0 cells are subjected to nutrient-rich medium. Fluorescence microscopy analysis of RNA content in cells after yFS240 G0 cells are exposed to nutrient-rich medium. Columns, left to right: differential interference contrast (DIC), DAPI fluorescence (DNA), Alexa Fluor 488 fluorescence (RNA) and merge.

**Figure S7.**
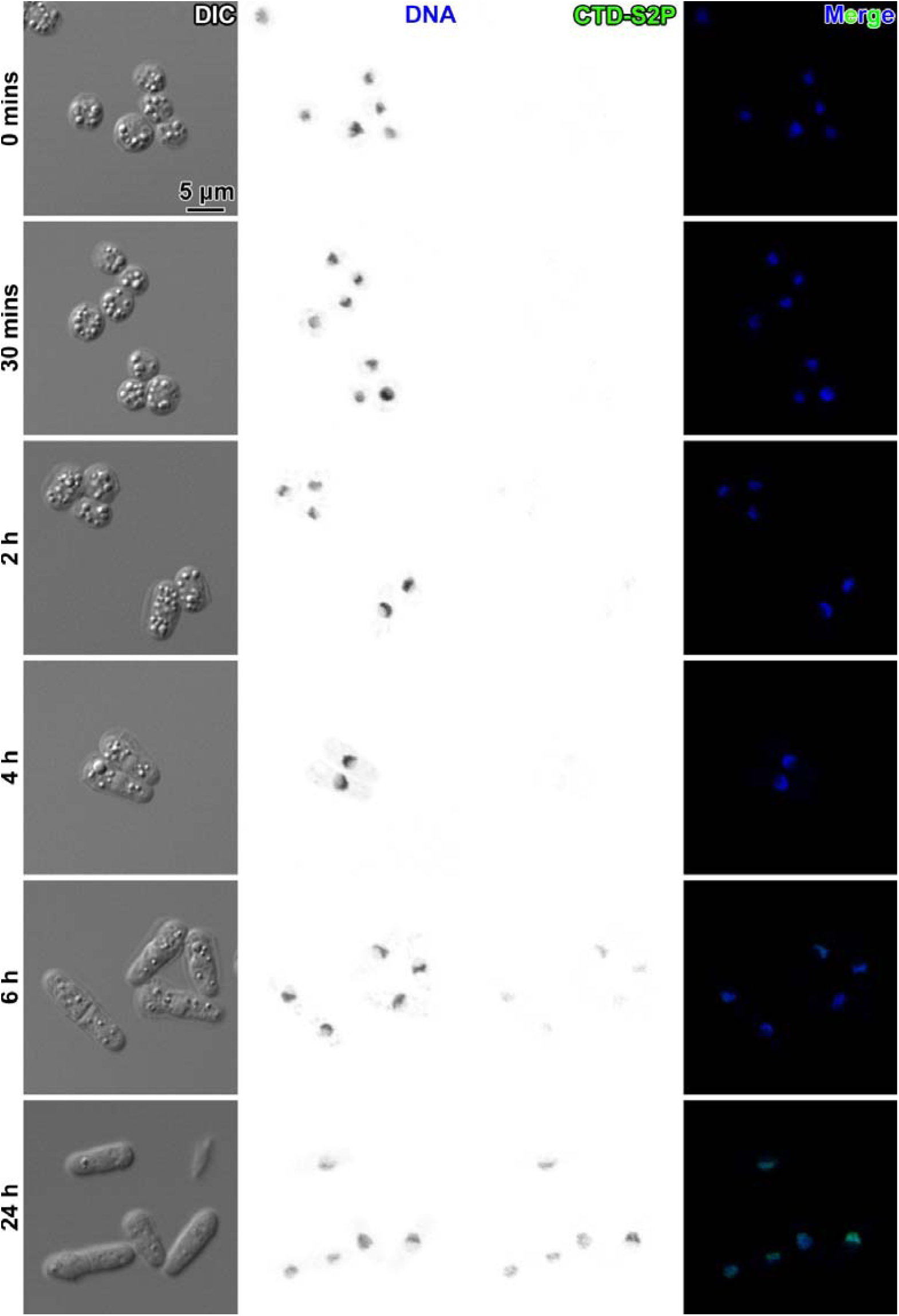
RNAPII-CTD-S2P levels increase 6 hours after G0 cells are subjected to nutrient-rich medium. Fluorescence microscopy analysis of RNAPII-CTD-S2P content in cells after wild-type G0 cells are exposed to nutrient-rich medium. Columns, left to right: differential interference contrast (DIC), DAPI fluorescence (DNA), Alexa Fluor 488 immunofluorescence (CTD-S2P) and merge.

**Figure S8.**
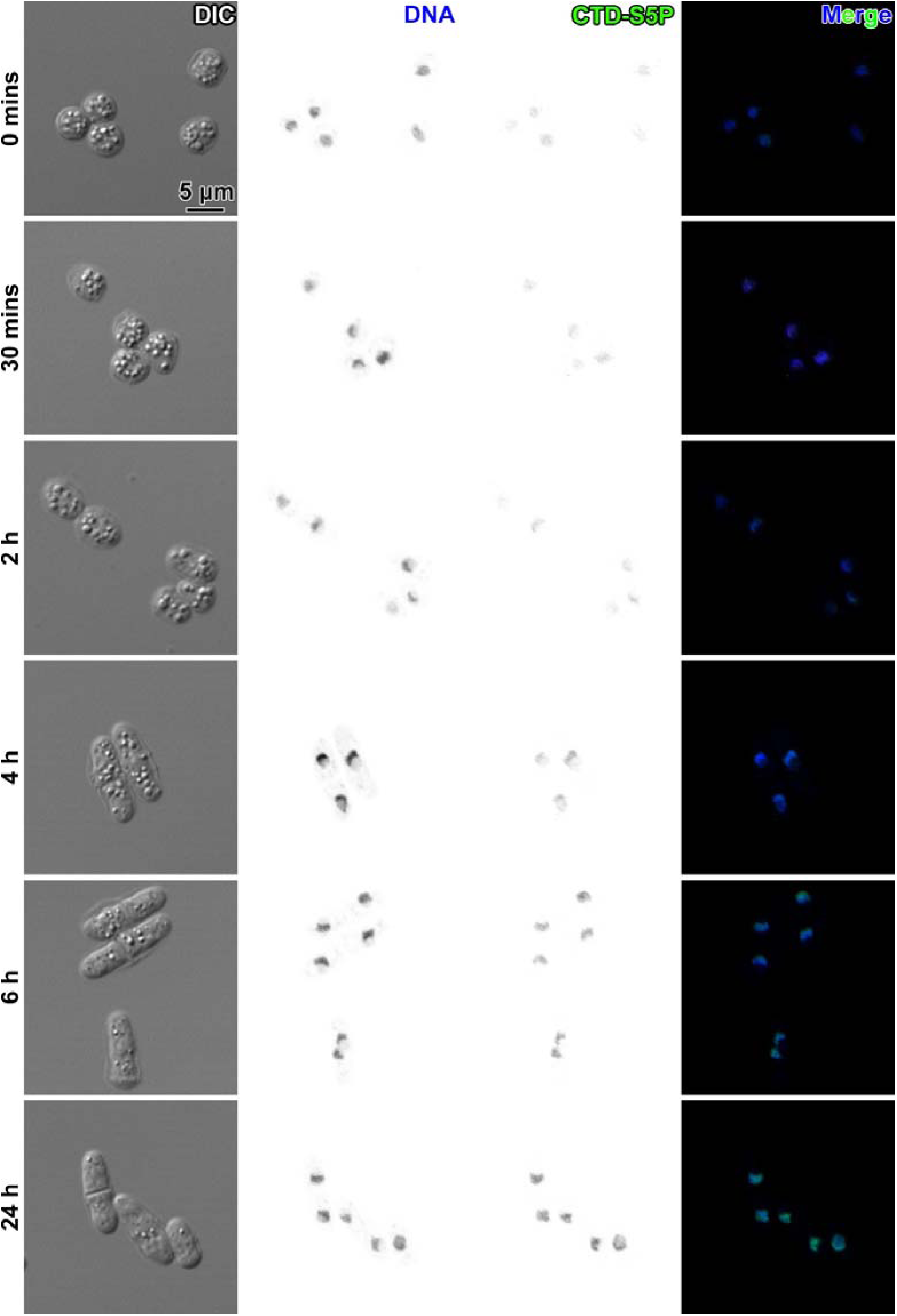
RNAPII-CTD-S5P levels increase 4 hours after G0 cells are changed into nutrient-rich medium. Fluorescence microscopy analysis of RNAPII-CTD-S5P content in cells after wild-type G0 cells are exposed to nutrient-rich medium. Columns, left to right: differential interference contrast (DIC), DAPI fluorescence (DNA), Alexa Fluor 488 immunofluorescence (CTD-S5P) and merge.

**Figure S9.**
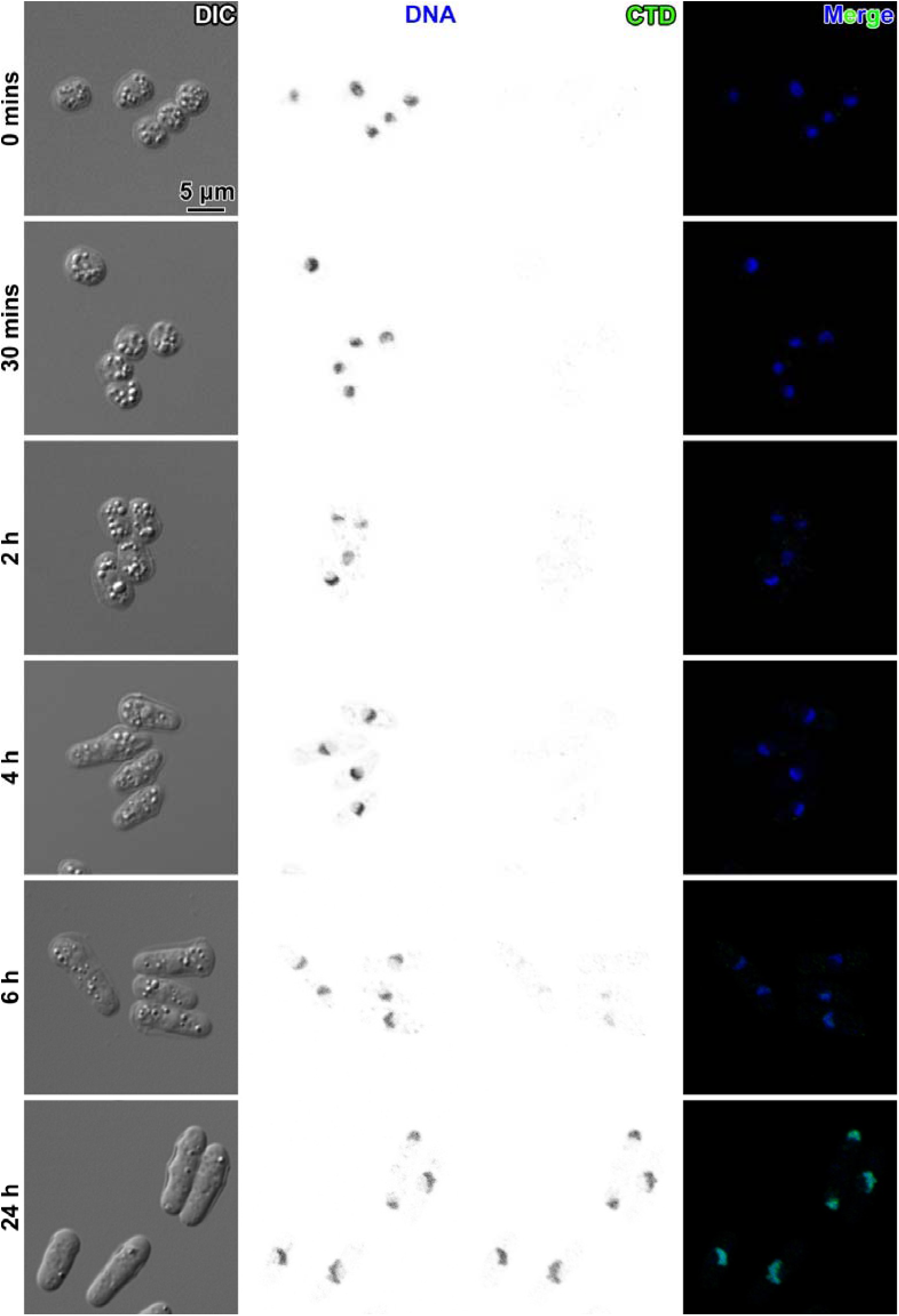
RNAPII CTDs become more detectable 6 hours after G0 cells are changed into nutrient-rich medium. Fluorescence microscopy analysis of RNAPII content in cells after wild-type G0 cells are exposed to nutrient-rich medium. Columns, left to right: differential interference contrast (DIC), DAPI fluorescence (DNA), Alexa Fluor 488 immunofluorescence (CTD) and merge.

**Figure S10.**
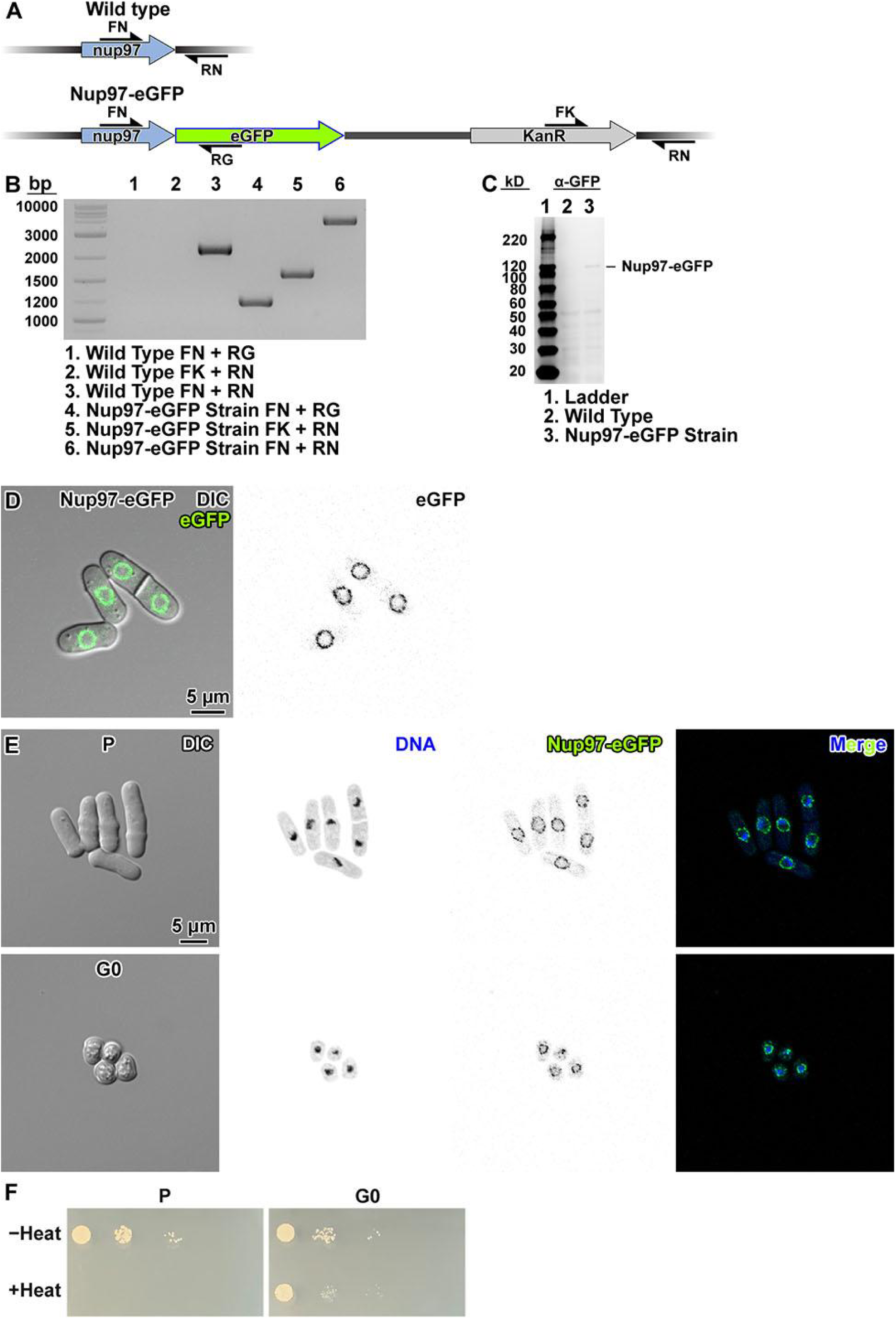
Construction of Nup97-eGFP strain. (A) Map of the *nup97* locus in the parent (wild-type) *S. pombe* strain MBY99 and the Nup97-eGFP strain LGSP0002. Primers used for PCR verification are indicated with the half arrow symbols. (B) Agarose gel of PCR amplicons expected from wild-type (control) and from Nup1-eGFP genomic DNA, in which the *nup97* locus is tagged with eGFP. (C) Immunoblot analysis of the Nup97-eGFP. The α-eGFP antibody correctly detected the Nup97-eGFP fusion protein in the newly created strain (lane 3), but not in the wild type (lane 2, negative control) in the uncropped α-eGFP immunoblot. (D) Differential interference contrast (DIC) and eGFP fluorescence confocal microscopy images of live Nup97-eGFP cells. In the left panel, a merge of the DIC channel and the eGFP fluorescence channel is shown. In the right panel, eGFP signals are rendered with inverted contrast. (E) Fluorescence microscopy analysis of Nup97-eGFP in interphase (P) and G0, in the Nup97-eGFP strain. Left to right: differential interference contrast (DIC), DAPI fluorescence (DNA) rendered in inverse contrast, eGFP fluorescence (Nup97-eGFP) rendered in inverse contrast, and merge of the two fluorescence channels. Cells were fixed with formaldehyde. (F) Spot tests of proliferating (P) and G0 cells of the Nup97-eGFP strain after a 30-minute incubation in YES and EMM−N respectively, with and without heat stress.

**Figure S11.**
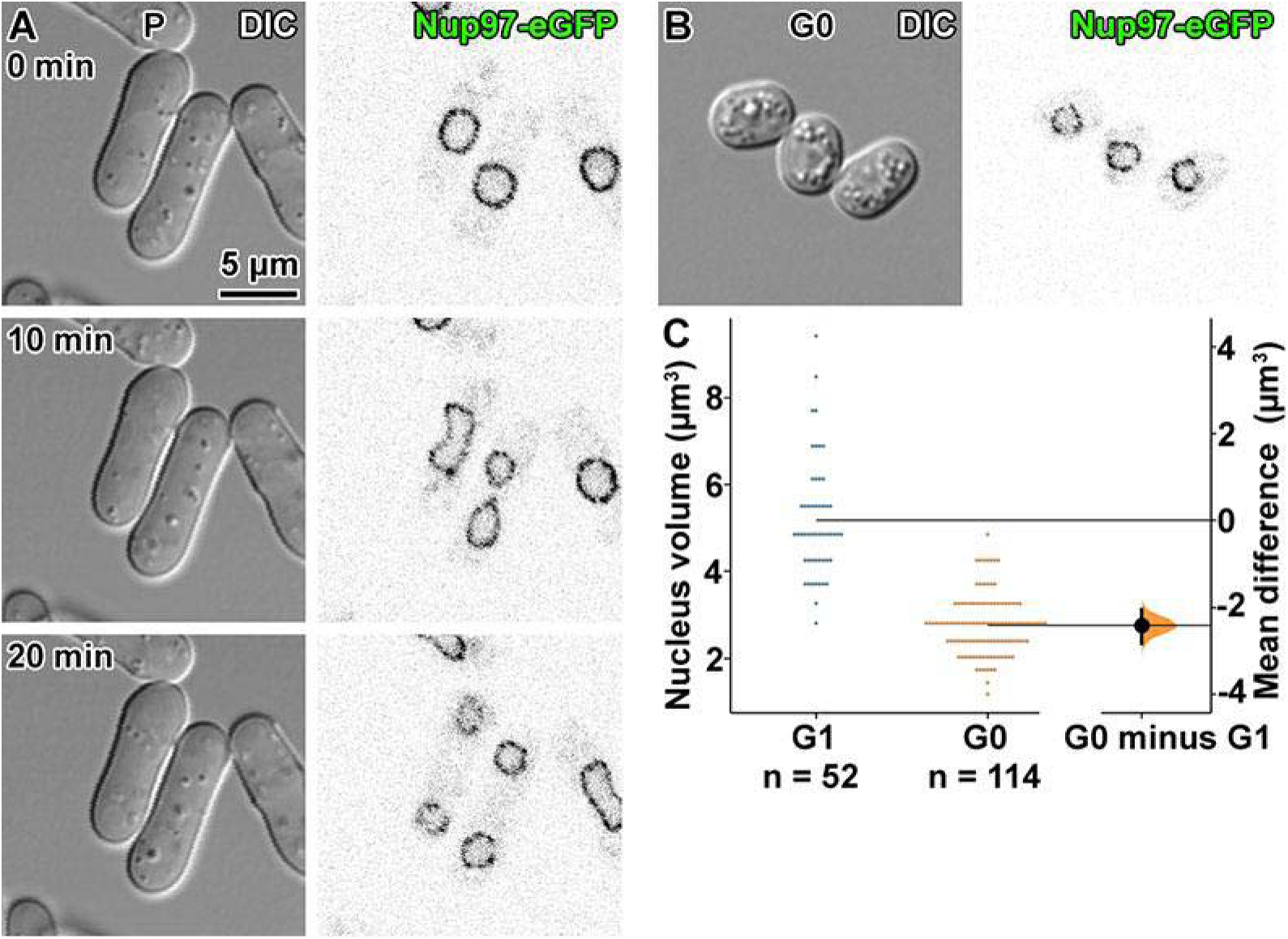
G0 Nup97-eGFP cells have smaller nuclei than G1 Nup97-eGFP cells. (A) Differential interference contrast (DIC) (left) and eGFP fluorescence (right) confocal fluorescence microscopy images of proliferating Nup97-eGFP cells just before (0 minutes), during (10 minutes), and just after (20 minutes) nuclear division. G1 nucleus sizes were determined from the nuclei that had just divided. See Movie S2 for the time-lapse images. (B) DIC and eGFP fluorescence confocal fluorescence microscopy images of G0 Nup97-eGFP cells. (C) Gardner-Altman plot of the mean difference between G1 and G0 nucleus volumes. Both groups are plotted on the left axes; the mean difference is plotted on a floating axis on the right. The mean difference is depicted as a dot; the 95% confidence interval is indicated by the ends of the vertical error bar (Ho et al., 2019).

**Movie S2. Time-lapse fluorescence microscopy of proliferating Nup97-eGFP cells, deposited on an agar pad.**

The images on the left are differential interference contrast (DIC), the images on the right are eGFP fluorescence (Nup97-eGFP). The interval between each frame is 10 minutes.

**Figure S12.**
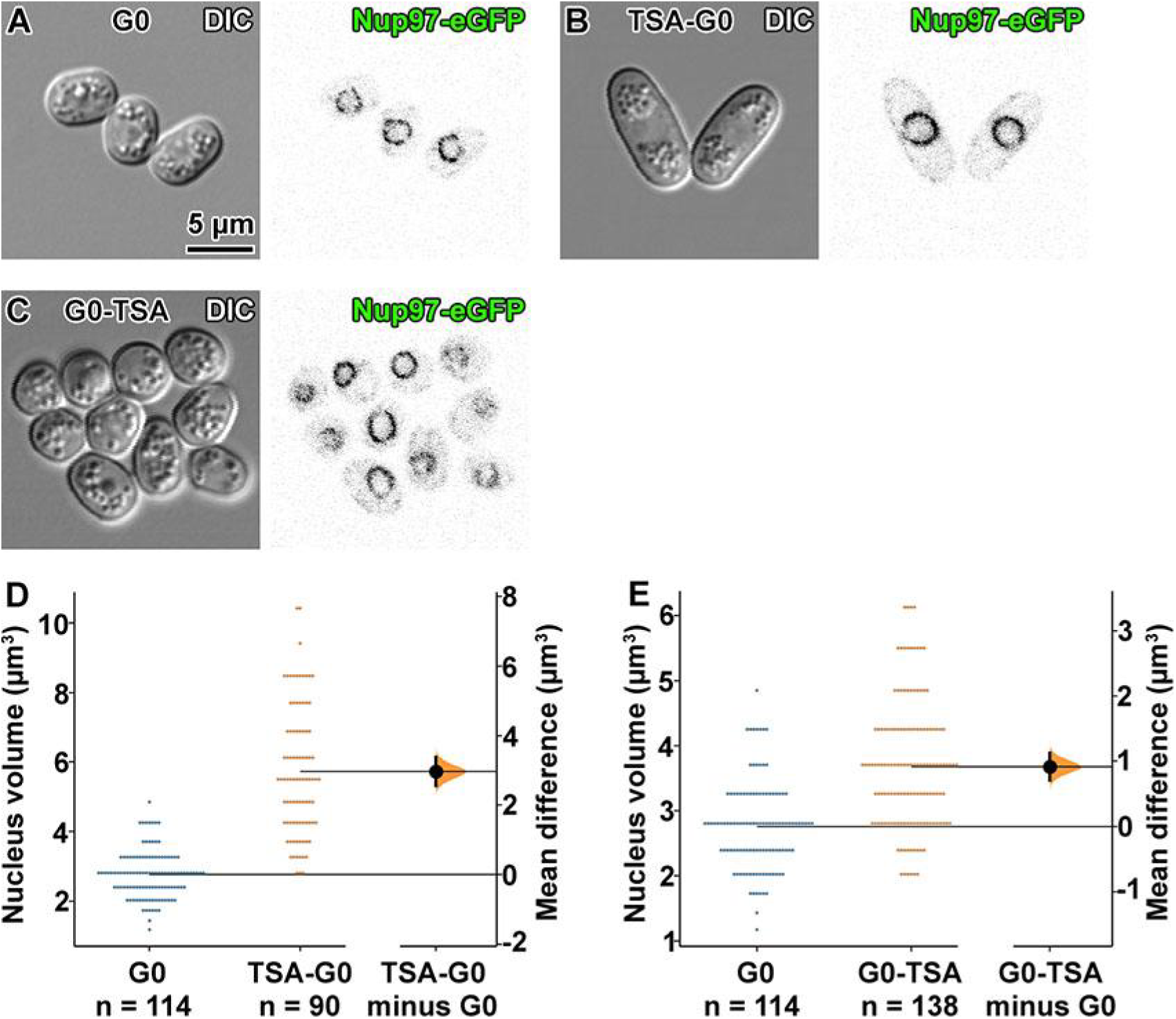
G0 Nup97-eGFP cells treated with TSA have larger nuclei than untreated G0 Nup97-eGFP cells. (A) Differential interference contrast (DIC) (left) and eGFP fluorescence (right) confocal fluorescence microscopy images of G0 Nup97-eGFP cells. (B) DIC and eGFP fluorescence confocal fluorescence microscopy images of TSA-G0 Nup97-eGFP cells. (C) DIC and eGFP fluorescence confocal fluorescence microscopy images of G0-TSA Nup97-eGFP cells. Gardner-Altman plot of (D) the mean difference between G0 and TSA-G0 nucleus volumes, and (E) the mean difference between G0 and G0-TSA nucleus volumes. In each plot, both groups are plotted on the left axes; the mean difference is plotted on a floating axis on the right. The mean difference is depicted as a dot; the 95% confidence interval is indicated by the ends of the vertical error bar (Ho et al., 2019).

**Figure S13.**
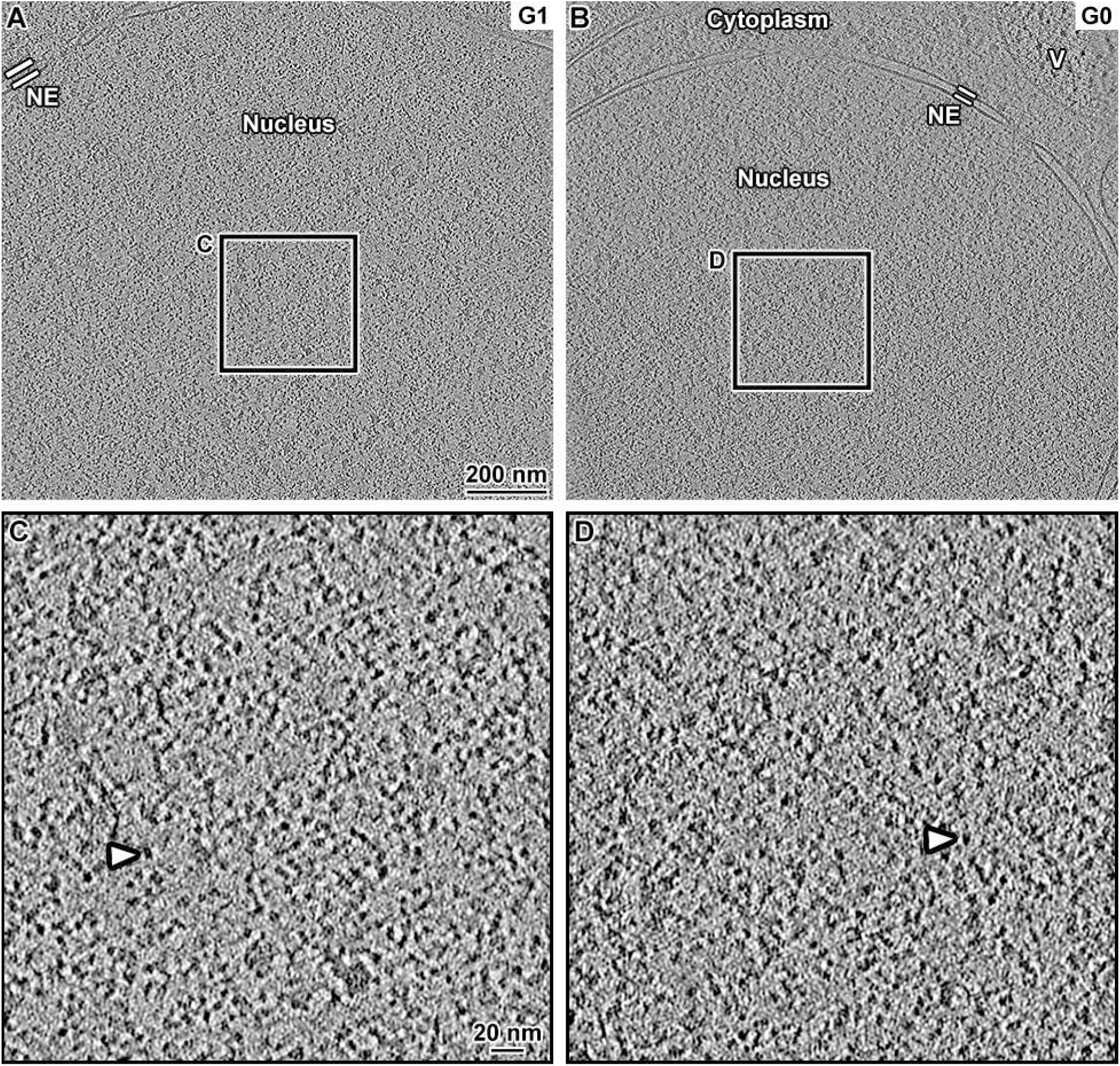
G0 cells have a denser nucleoplasm than G1 cells. Defocus phase-contrast cryotomographic slices (12 nm) of a G1-arrested (A) and a G0 (B) cell, imaged with defocus contrast. NE: nuclear envelope. (C and D) Four-fold enlargements of the regions boxed in panels A and B, respectively. Nucleosome-like densities are indicated by arrowheads.

**Figure S14.**
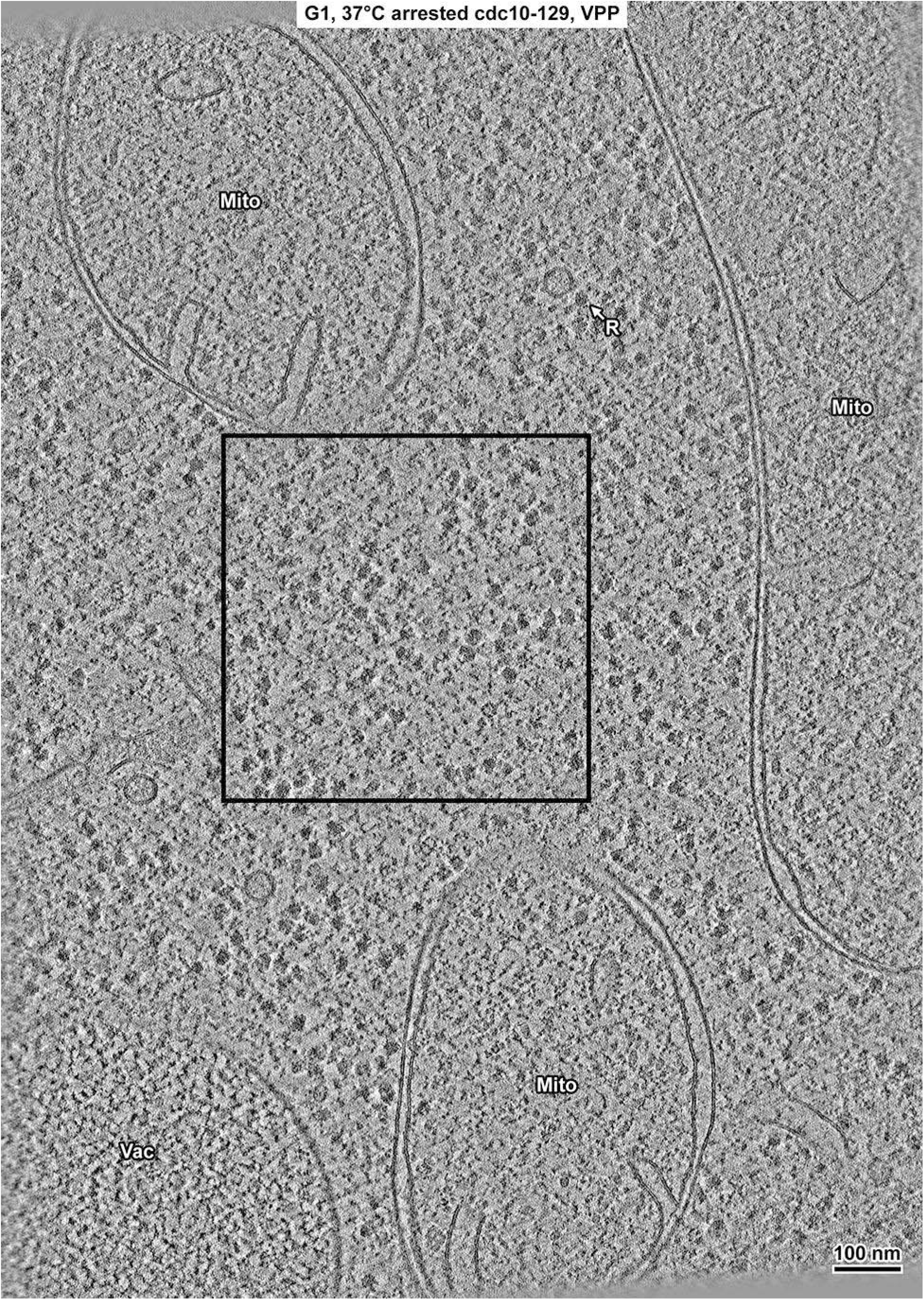
Overview of a G1 cytoplasm cryolamella. Volta cryotomographic slice (12 nm) of a ribosome-rich region of the cytoplasm in a G1-arrested *cdc10-129* cell cryolamella. The vacuoles (Vac), mitochondria (Mito), and a ribosome (R) are indicated. The boxed region was used for Fourier power spectrum analysis in Figures 6 and S28.

**Figure S15.**
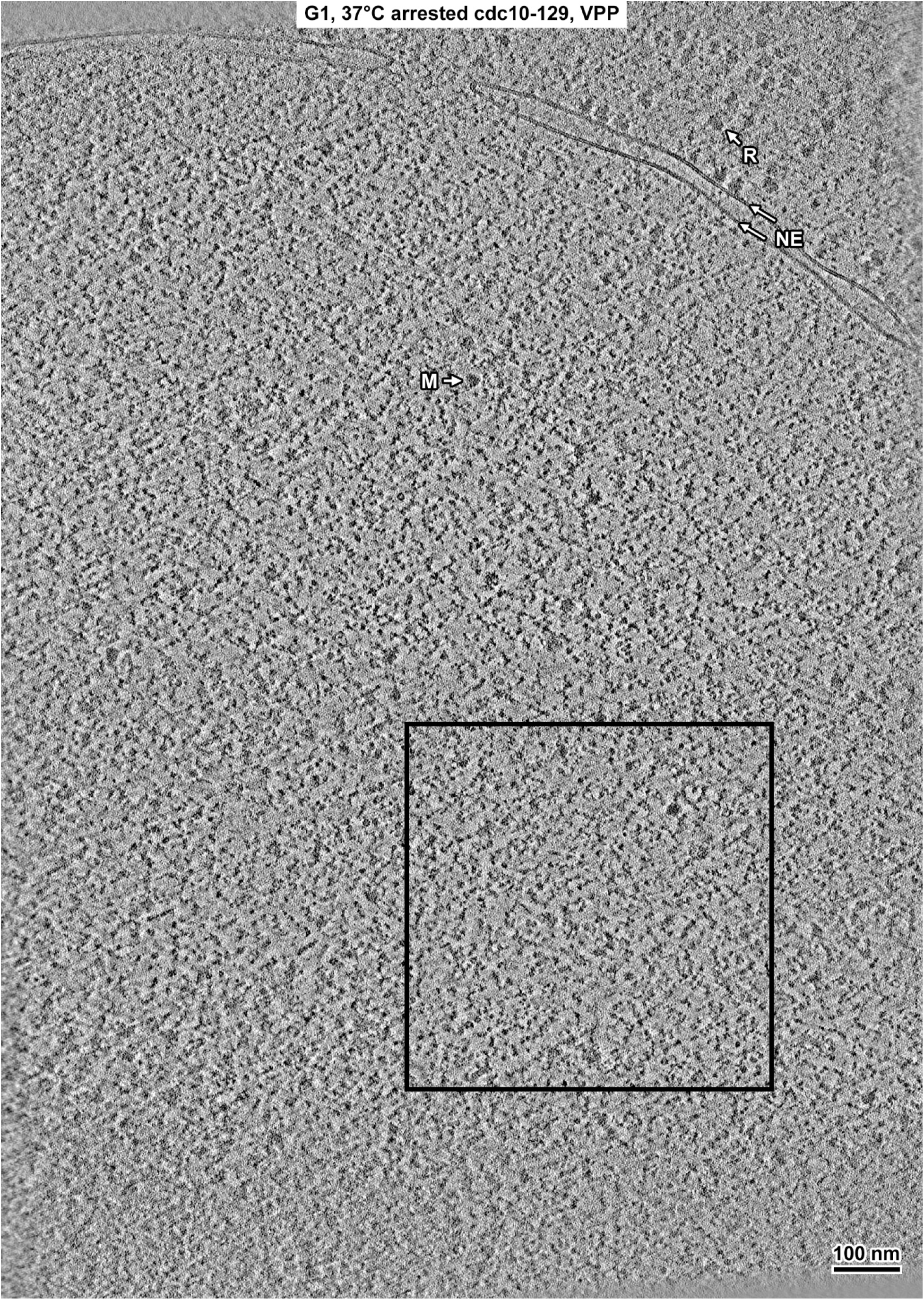
Overview of a G1 nucleus cryolamella. Volta cryotomographic slice (12 nm) of the nucleus in a G1-arrested *cdc10-129* cell cryolamella. Some non-chromatin features are indicated: nuclear envelope (NE), megacomplex (M), and ribosome (R). The boxed region is shown with twofold enlargement in Figure S16, and was used for Fourier power spectrum analysis in Figure 6.

**Figure S16.**
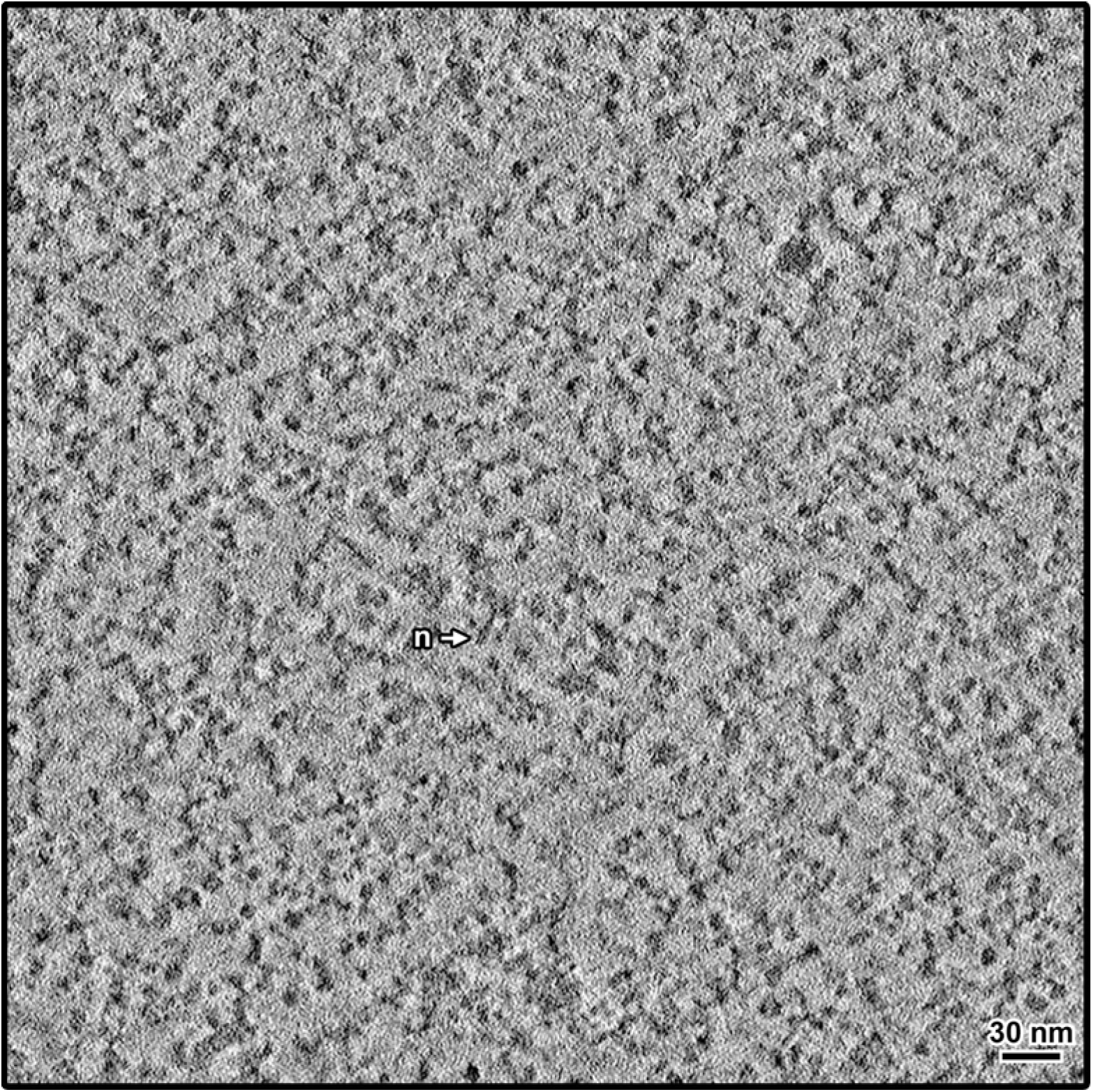
Twofold enlargement of selected region from G1 nucleus cryolamella. A region of a Volta cryotomographic slice (12 nm) of the nucleus in the G1-arrested *cdc10-129* cell from Figure S15, selected for Fourier power spectrum analysis in Figure 6. A nucleosome-like particle (n) is indicated.

**Figure S17.**
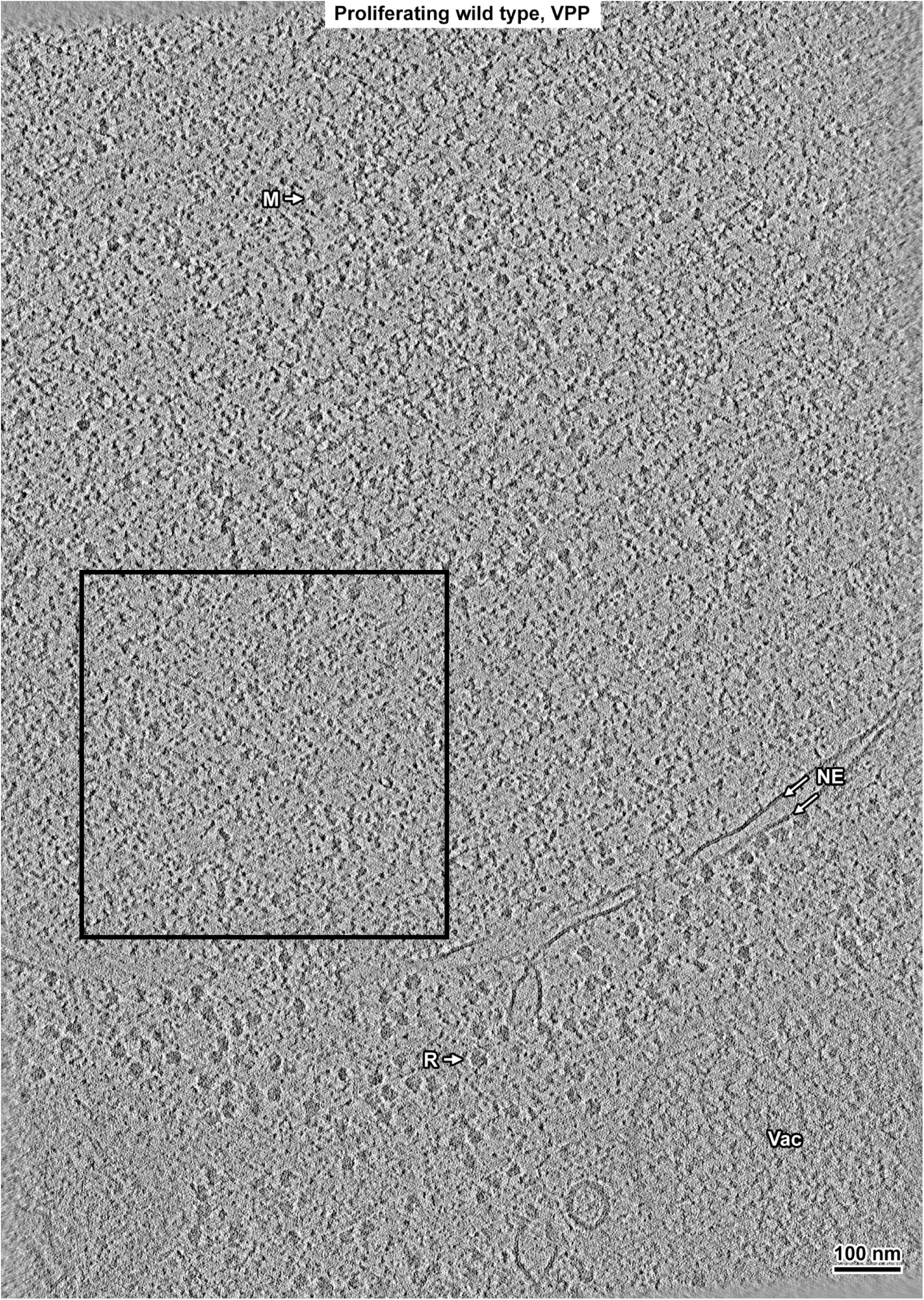
Overview of a proliferating cell nucleus cryolamella. Volta cryotomographic slice (12 nm) of the nucleus in a proliferating cell cryolamella. Some non-chromatin features are indicated: nuclear envelope (NE), megacomplex (M), ribosome (R) and vacuole (Vac). The boxed region is shown with twofold enlargement in Figure S18, and was used for Fourier power spectrum analysis in Figure 6.

**Figure S18.**
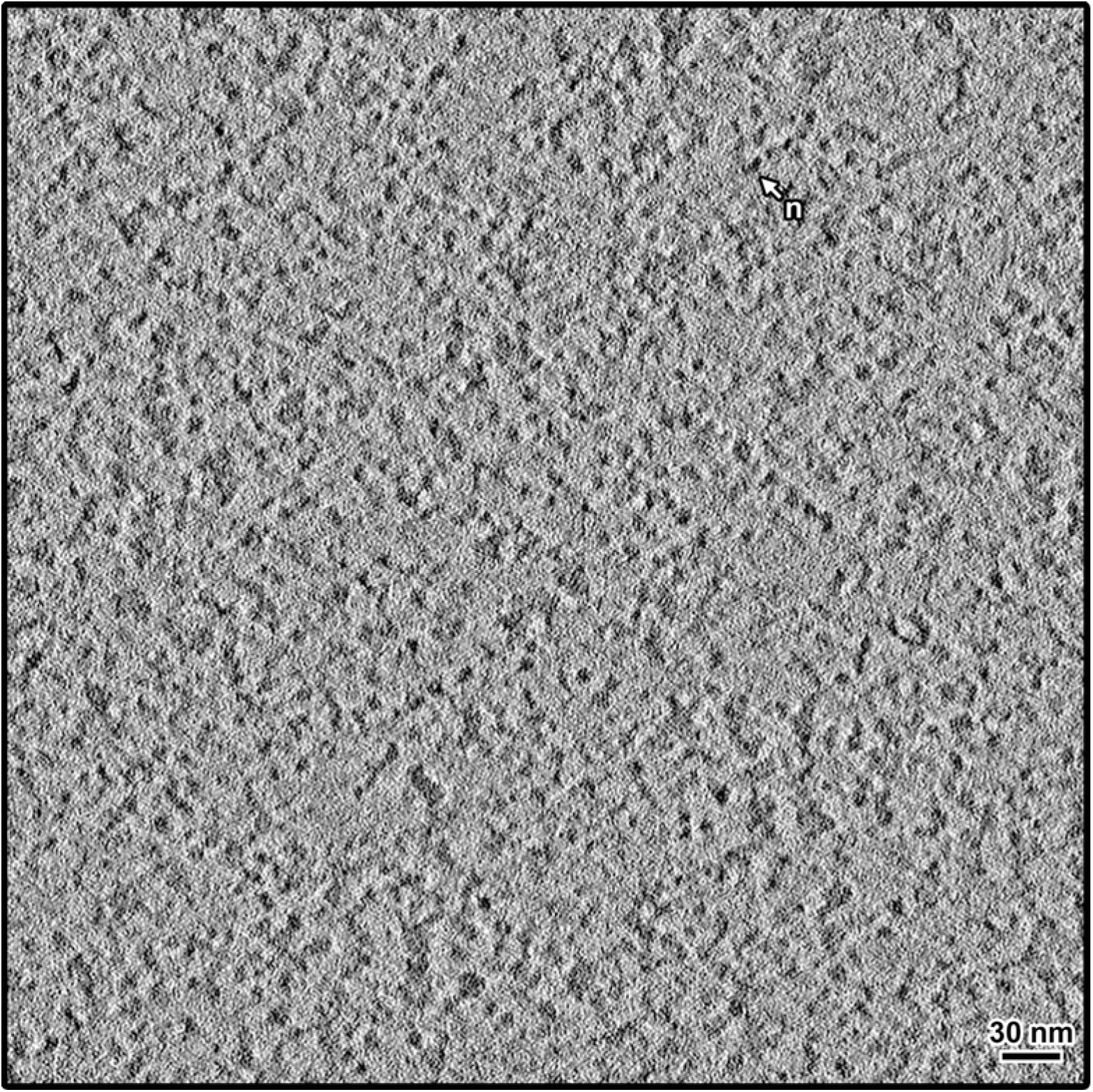
Twofold enlargement of selected region from proliferating nucleus cryolamella. A region of a Volta cryotomographic slice (12 nm) of the nucleus in the proliferating cell from Figure S17, selected for Fourier power spectrum analysis in Figure 6. A nucleosome-like particle (n) is indicated.

**Figure S19.**
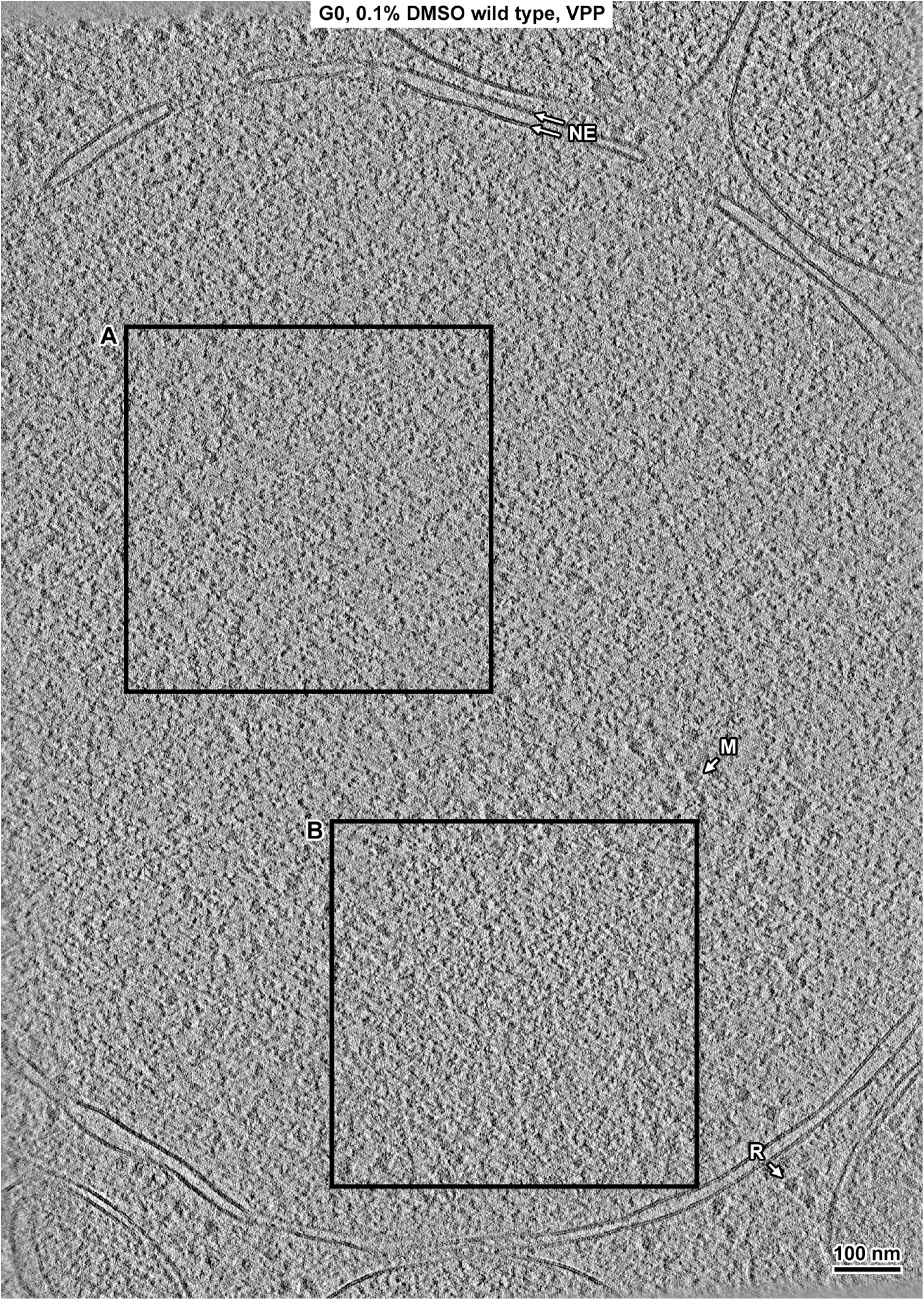
Overview of a G0 nucleus cryolamella. Volta cryotomographic slice (12 nm) of a nucleus cryolamella of a wild-type G0 cell treated with 0.1% DMSO during G0 entry. Some non-chromatin features are indicated: nuclear envelope (NE), megacomplex (M), and ribosome (R). The nucleoplasm boxed region (A) is shown with twofold enlargement in Figure S20, and was used for Fourier power spectrum analysis in Figures 6 and S28. The nucleolus boxed region (B) is shown with twofold enlargement in Figure S21, and was used for Fourier power spectrum analysis in Figure S28.

**Figure S20.**
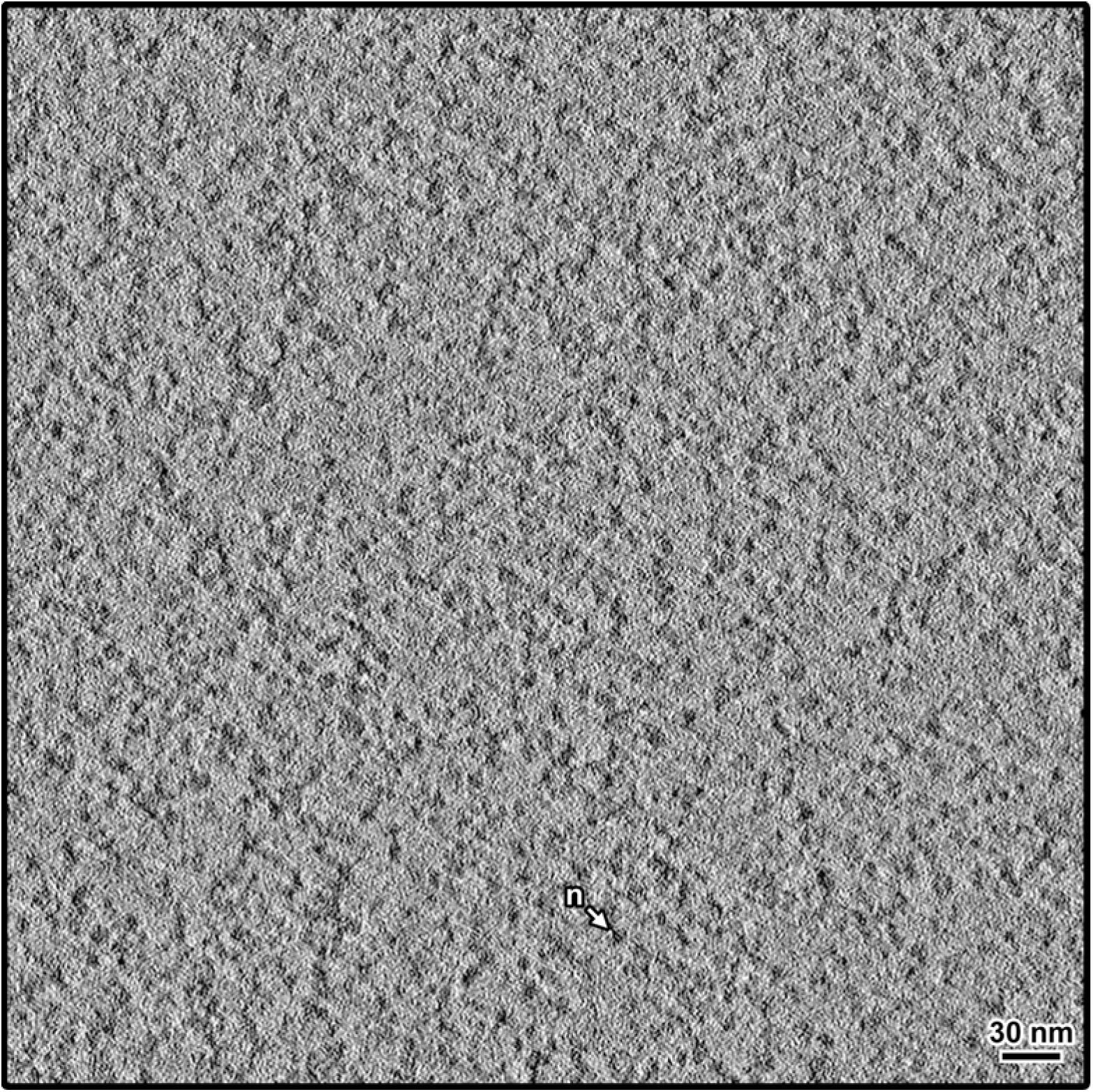
Twofold enlargement of selected nucleoplasm region from a G0 nucleus cryolamella. A region of a Volta cryotomographic slice (12 nm) of the nucleoplasm region of the nucleus in a wild-type G0 cell treated with 0.1% DMSO during G0 entry from Figure S19, selected for Fourier power spectrum analysis in Figures 6 and S28. A nucleosome-like particle (n) is indicated.

**Figure S21.**
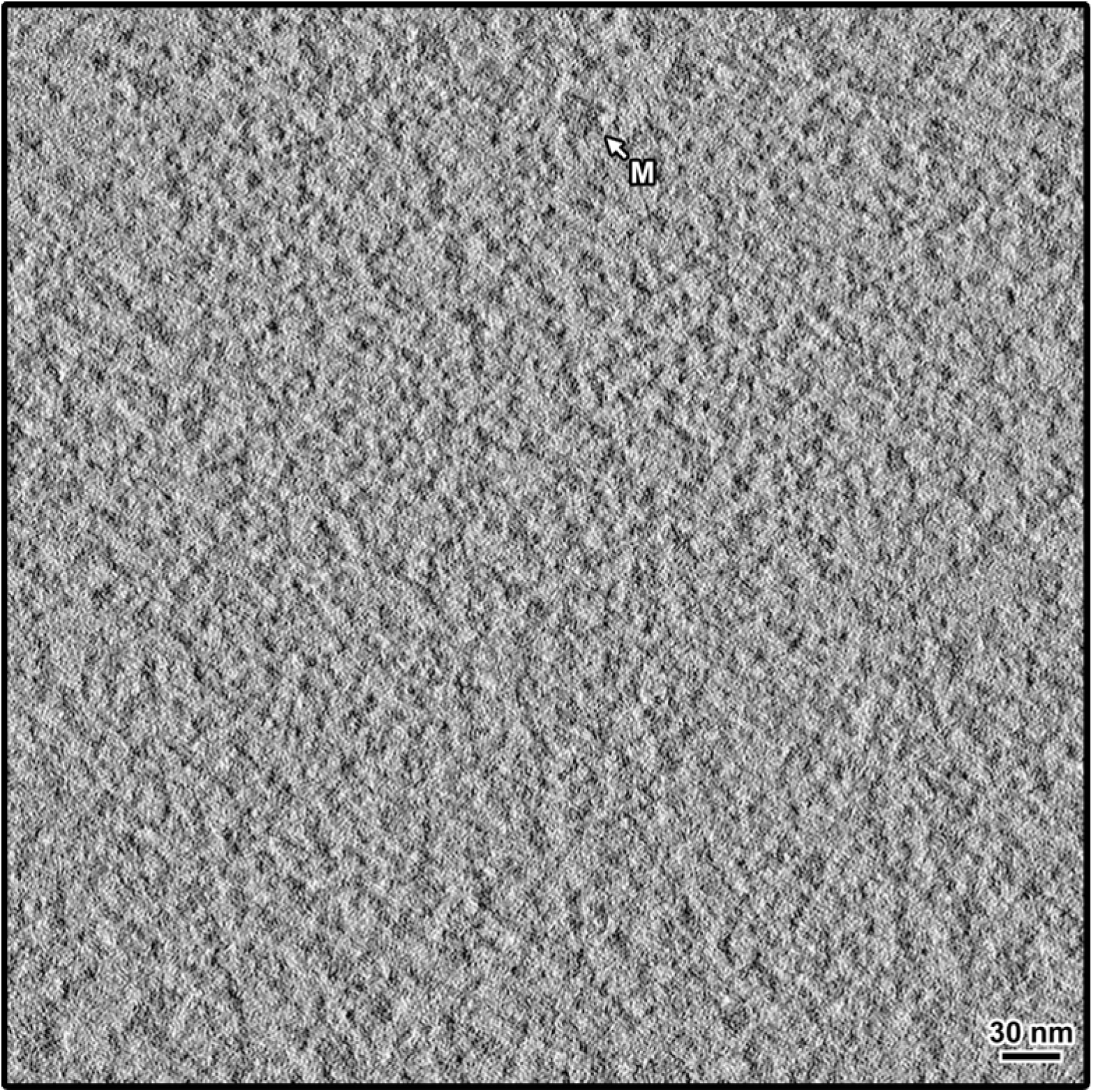
Twofold enlargement of selected nucleolus region from a G0 nucleus cryolamella. A region of a Volta cryotomographic slice (12 nm) of the nucleolus in a wild-type G0 cell treated with 0.1% DMSO during G0 from Figure S20, selected for Fourier power spectrum analysis in Figure S28. A megacomplex (M) is indicated.

**Figure S22.**
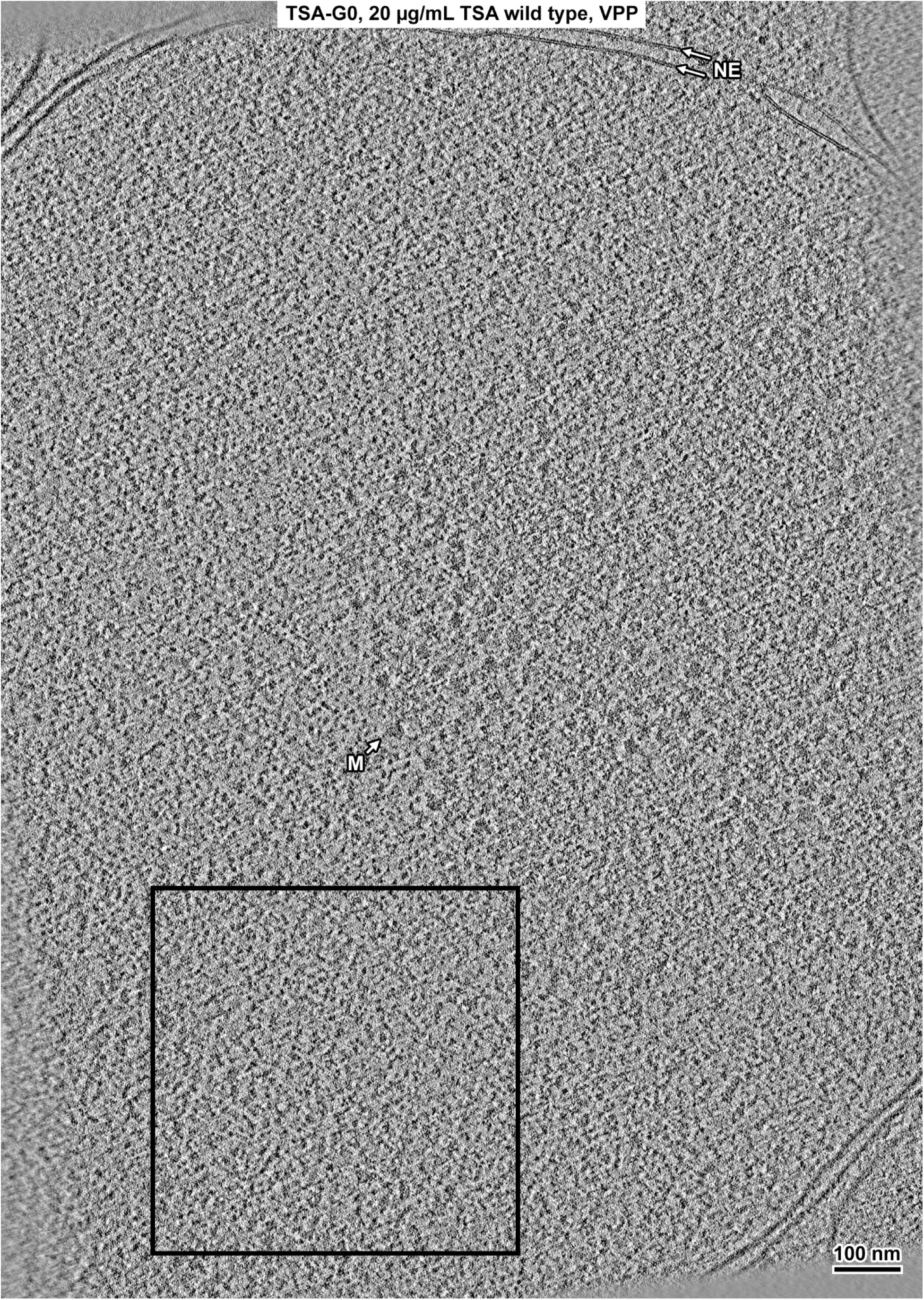
Overview of a TSA-G0 nucleus cryolamella. Volta cryotomographic slice (12 nm) of a nucleus cryolamella of a wild-type G0 cell treated with 20 ng/µL TSA during G0 entry. Some non-chromatin features are indicated: nuclear envelope (NE) and megacomplex (M). The boxed region is shown with twofold enlargement in Figure S23, and was used for Fourier power spectrum analysis in Figure 6.

**Figure S23.**
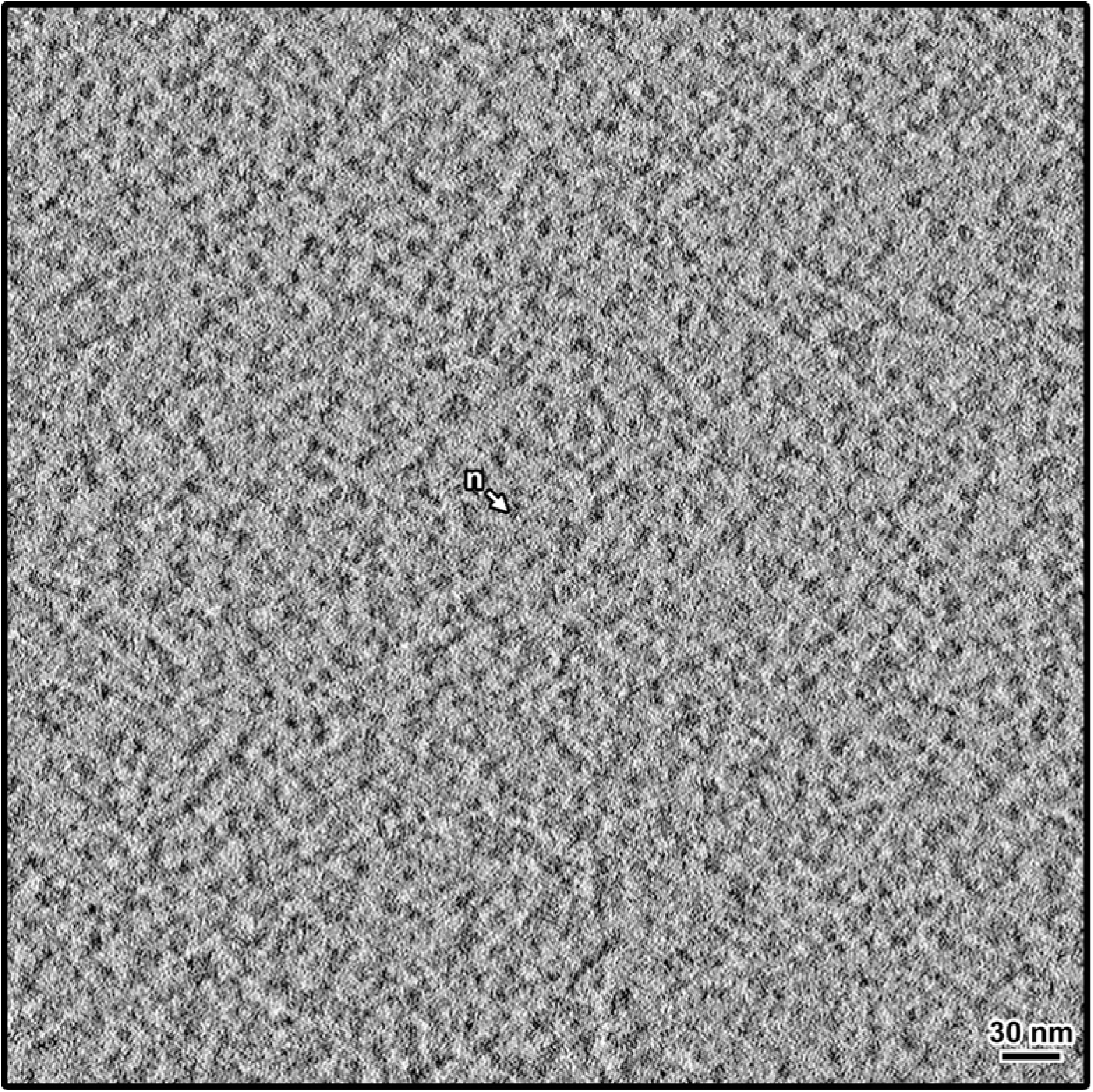
Twofold enlargement of selected region from TSA-G0 nucleus cryolamella. A region of a Volta cryotomographic slice (12 nm) of nucleus in a wild-type G0 cell treated with 20 ng/µL TSA during G0 entry from Figure S22, selected for Fourier power spectrum analysis in Figure 6. A nucleosome-like particle (n) is indicated.

**Figure S24.**
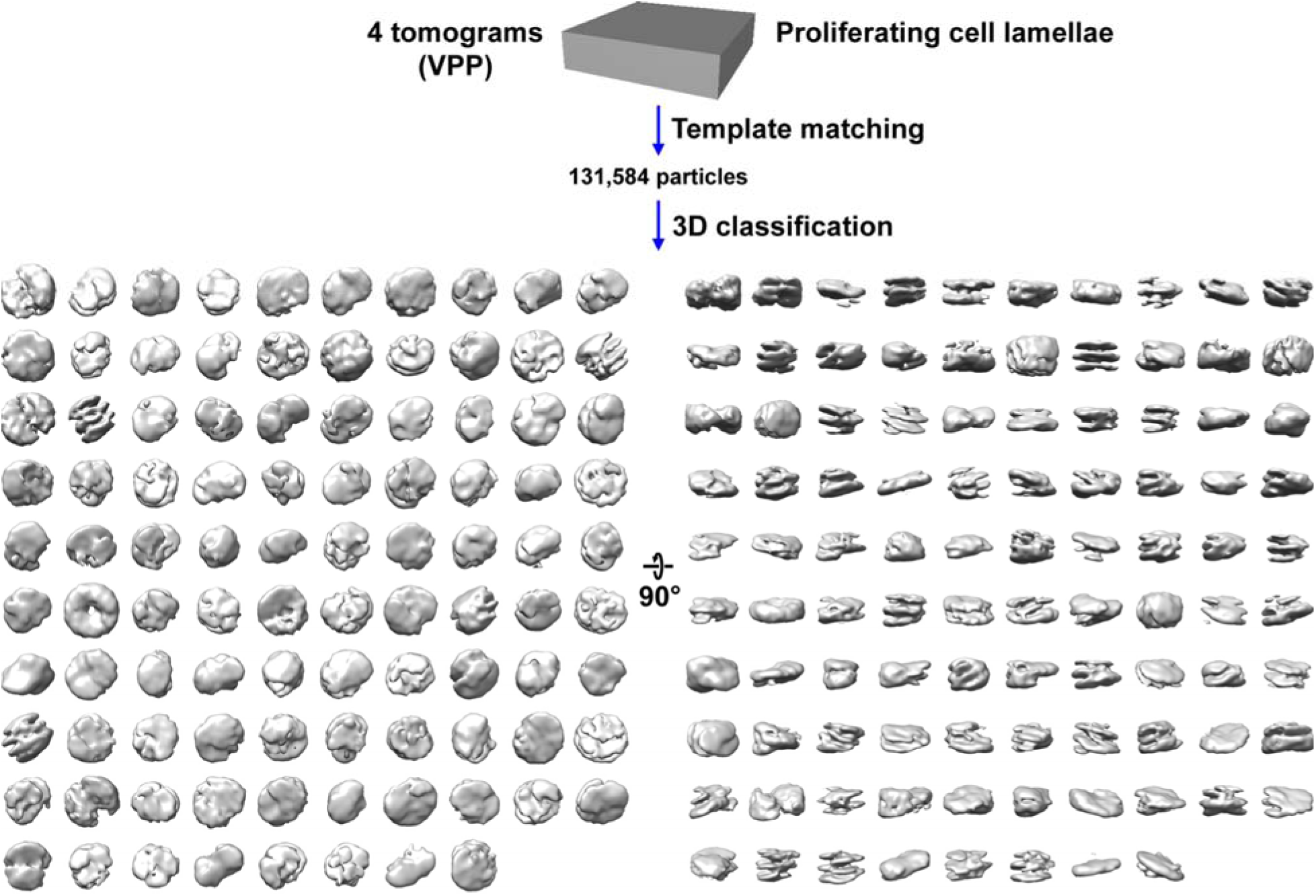
Direct 3-D classification of proliferating cell cryolamellae densities. Tomograms of proliferating cell cryolamellae were template matched using a smooth 5 nm radius 6 nm thick cylindrical reference and a 5.4 nm radius 6.1 nm thick cylindrical mask. Direct 3-D class averaging (2-D classification was bypassed) was done using a smooth 5 nm radius 6 nm thick cylindrical reference and a 5 nm radius 6 nm thick cylindrical mask with a cosine-shaped edge, with 100 classes. None of the class averages (grey) of the nucleosome-like template-matching hits resembled a canonical nucleosome. Two classes have no contributing particles.

**Figure S25.**
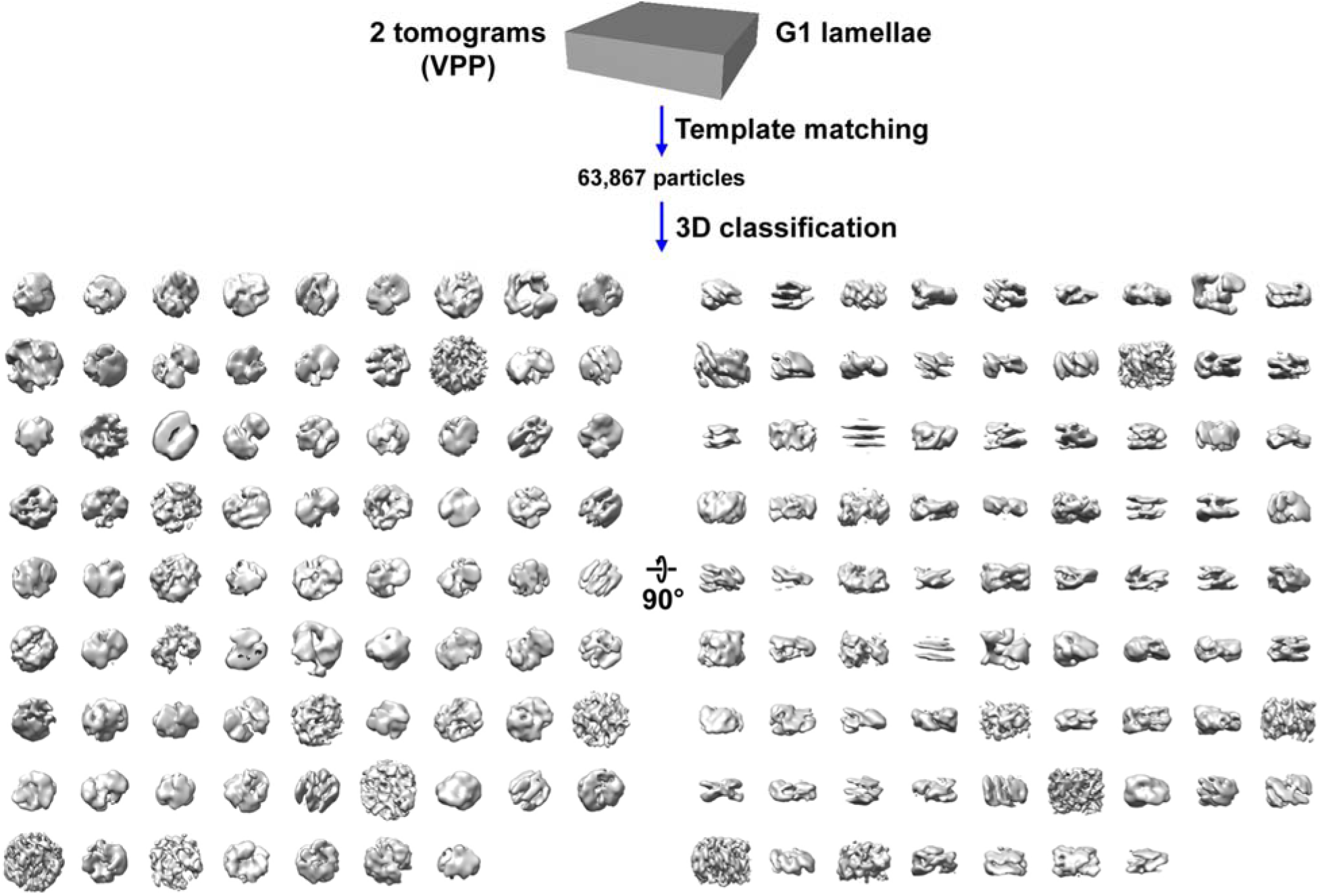
Direct 3-D classification of cdc10-129 G1 cryolamellae densities. Tomograms of *cdc10-129* G1 cryolamellae were template matched using a smooth 5 nm radius 6 nm thick cylindrical reference and a 5.4 nm radius 6.1 nm thick cylindrical mask. Direct 3-D class averaging (2-D classification was bypassed) was done using a smooth 5 nm radius 6 nm thick cylindrical reference and a 5 nm radius 6 nm thick cylindrical mask with a cosine-shaped edge, with 100 classes. None of the class averages (grey) of the nucleosome-like template-matching hits resembled a canonical nucleosome. Twenty-one classes have no contributing particles.

**Figure S26.**
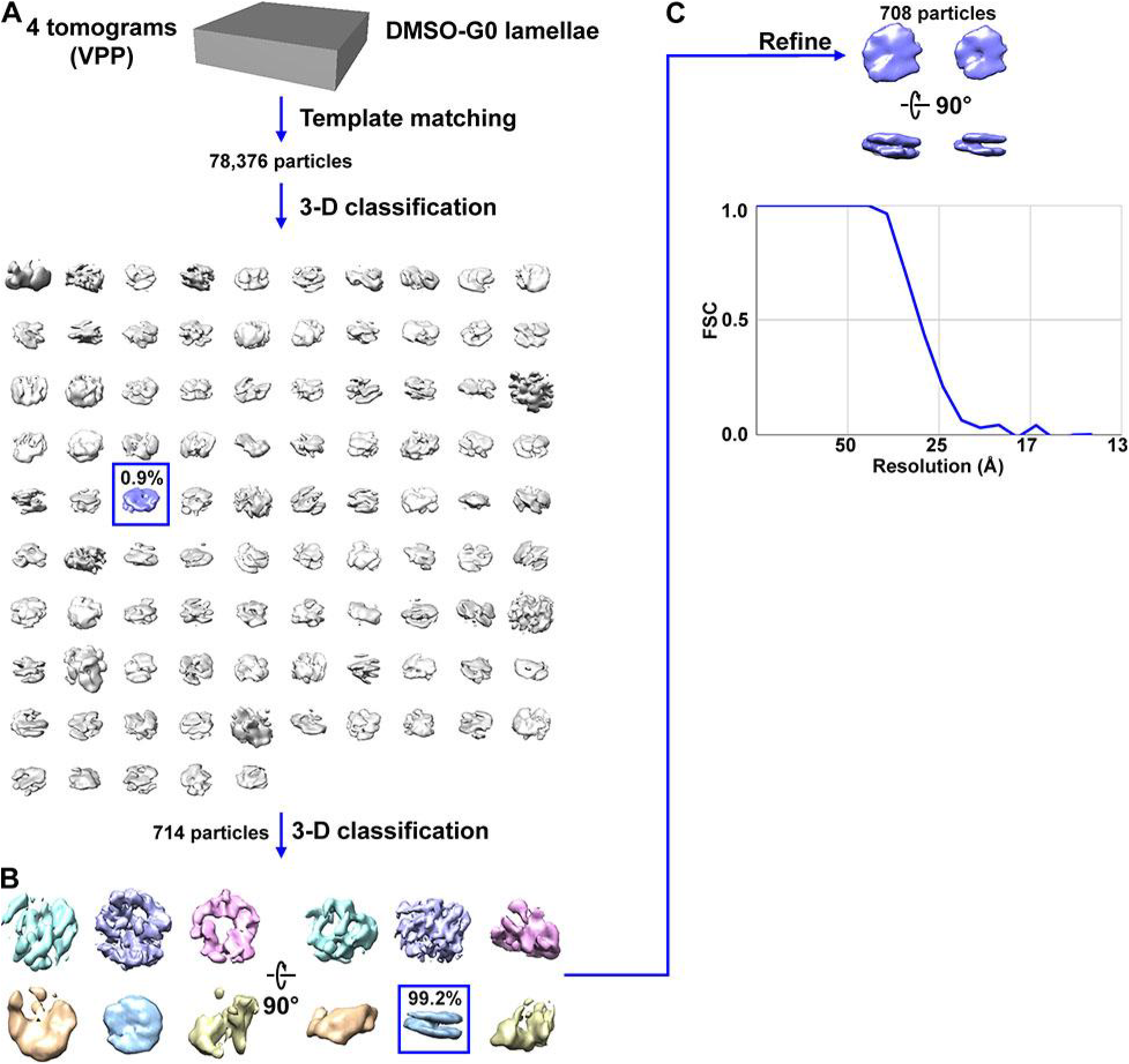
Direct 3-D classification of DMSO-G0 cryolamellae densities. (A) Tomograms of DMSO-G0 cryolamellae were template matched using a smooth 5 nm radius 6 nm thick cylindrical reference and a 5.4 nm radius 6.1 nm thick cylindrical mask. Direct 3-D class averaging (2-D classification was bypassed) was done using a smooth 5 nm radius 6 nm thick cylindrical reference and a 5 nm radius 6 nm thick cylindrical mask with a cosine-shaped edge, with 100 classes. The canonical nucleosome-like class average is shaded blue. Five classes have no contributing particles. (B) The second round of 3-D classification from the canonical nucleosome-like class from panel A, with 10 classes. Four classes have no contributing particles. (C) Refined density of candidate nucleosomes observed in DMSO-G0 cryolamellae. The density views on the left have a contour level of 0.25, the density views on the right have a contour level of 0.50. The resolution is ∼28 Å by the FSC = 0.5 criterion.

**Figure S27.**
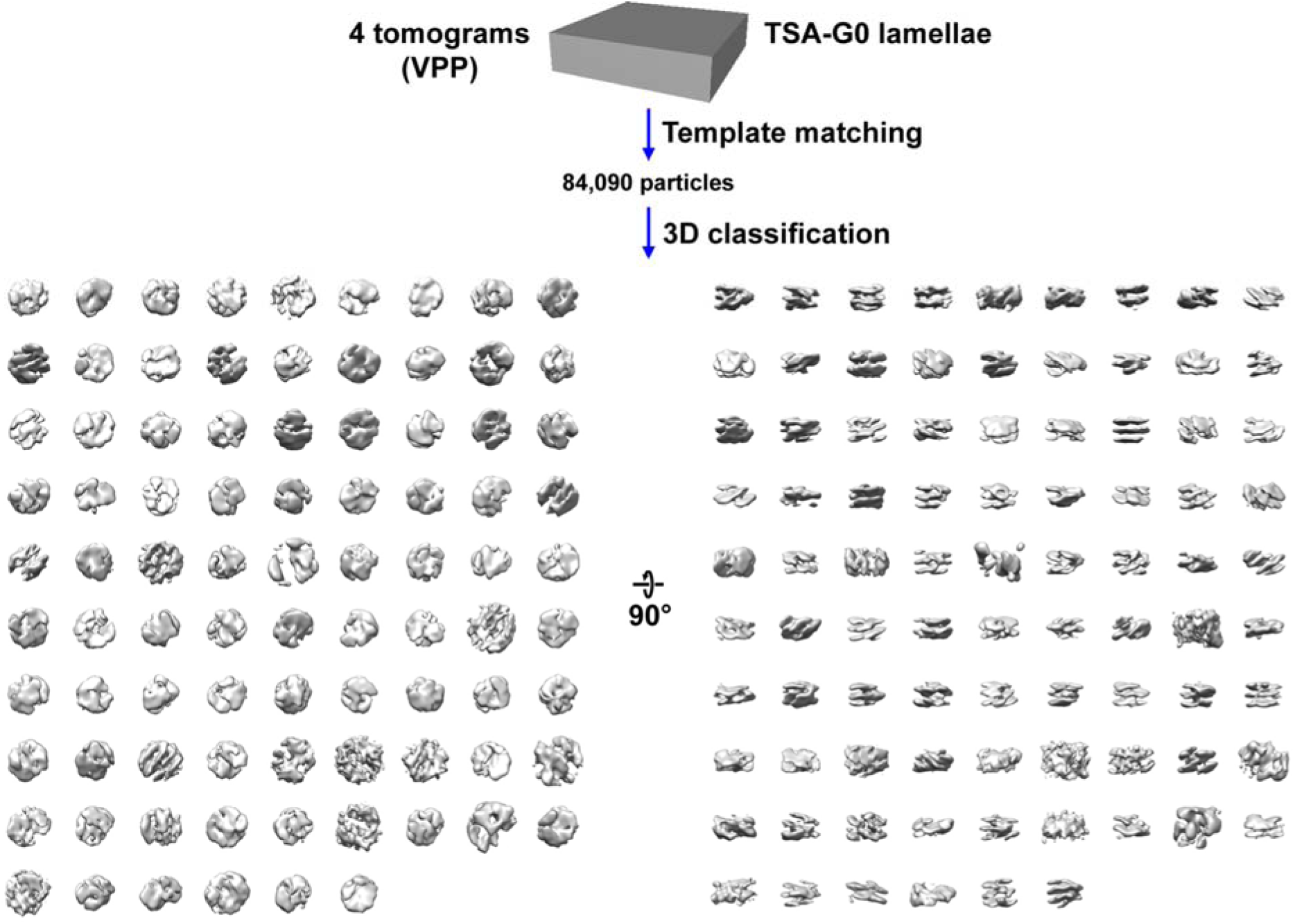
Direct 3-D classification of TSA-G0 cryolamellae densities. Tomograms of TSA-G0 cryolamellae were template matched using a smooth 5 nm radius 6 nm thick cylindrical reference and a 5.4 nm radius 6.1 nm thick cylindrical mask. Direct 3-D class averaging (2-D classification was bypassed) was done using a smooth 5 nm radius 6 nm thick cylindrical reference and a 5 nm radius 6 nm thick cylindrical mask with a cosine-shaped edge, with 100 classes. None of the class averages (grey) of nucleosome-like template-matching hits resembled a canonical nucleosome. Thirteen classes have no contributing particles.

**Figure S28.**
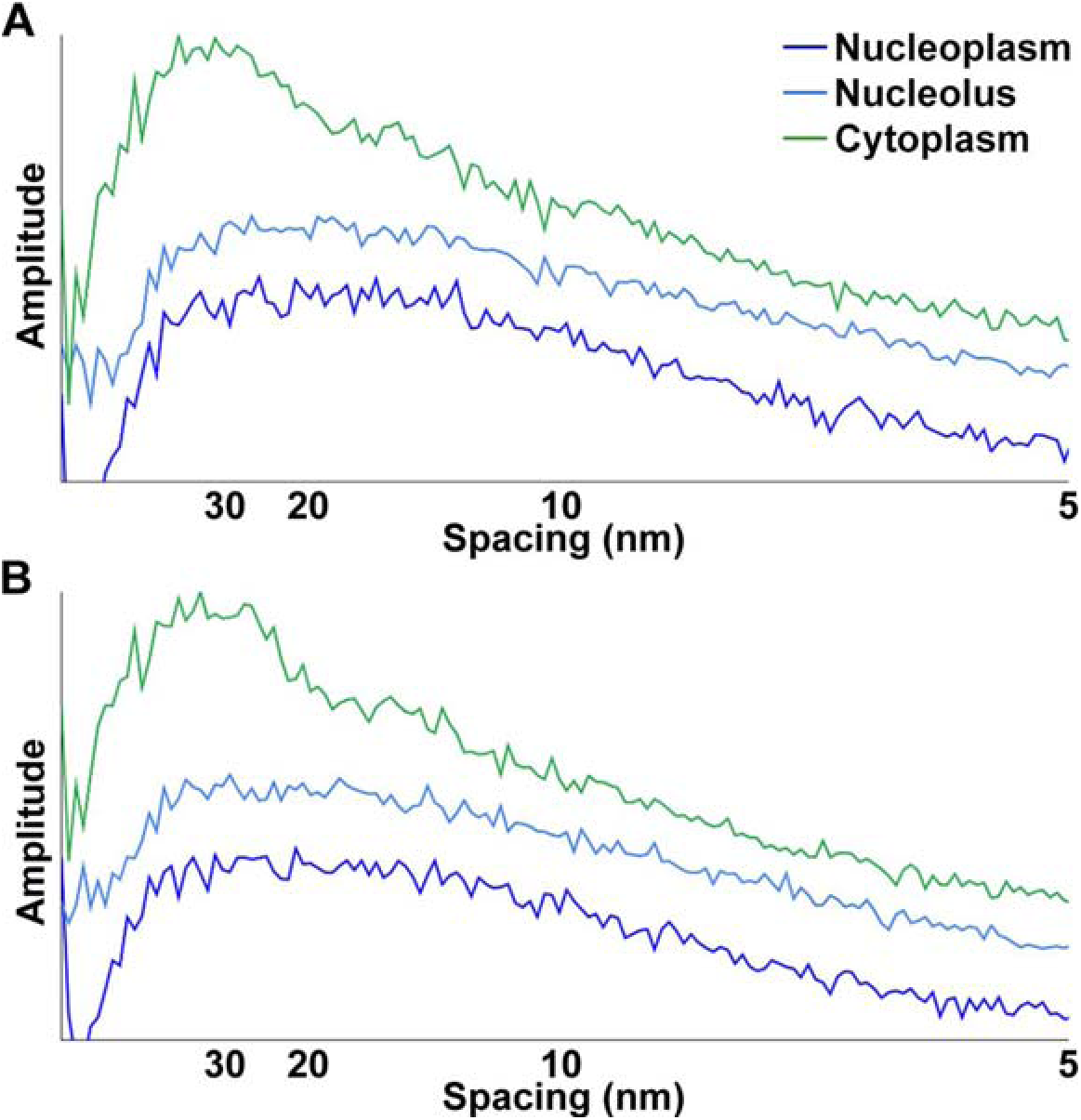
G0 nucleoplasm and nucleolus power spectra are similar to each other, but do not have the 30 nm peak present in the cytoplasm power spectrum. Rotationally averaged 1-D Fourier power spectra of chosen nucleoplasm (blue) and nucleolus (light blue) regions in the G0 tomogram analysed, with the Fourier power spectrum of a ribosome-rich cytoplasm region in a G1 tomogram (green) for comparison. Fourier analysis was done on Volta cryotomographic slices extracted at (A) 12 nm and (B) 30 nm thickness. All power spectra for each thickness are plotted on the same graph, separated with arbitrary offsets for clarity.

**Figure S29.**
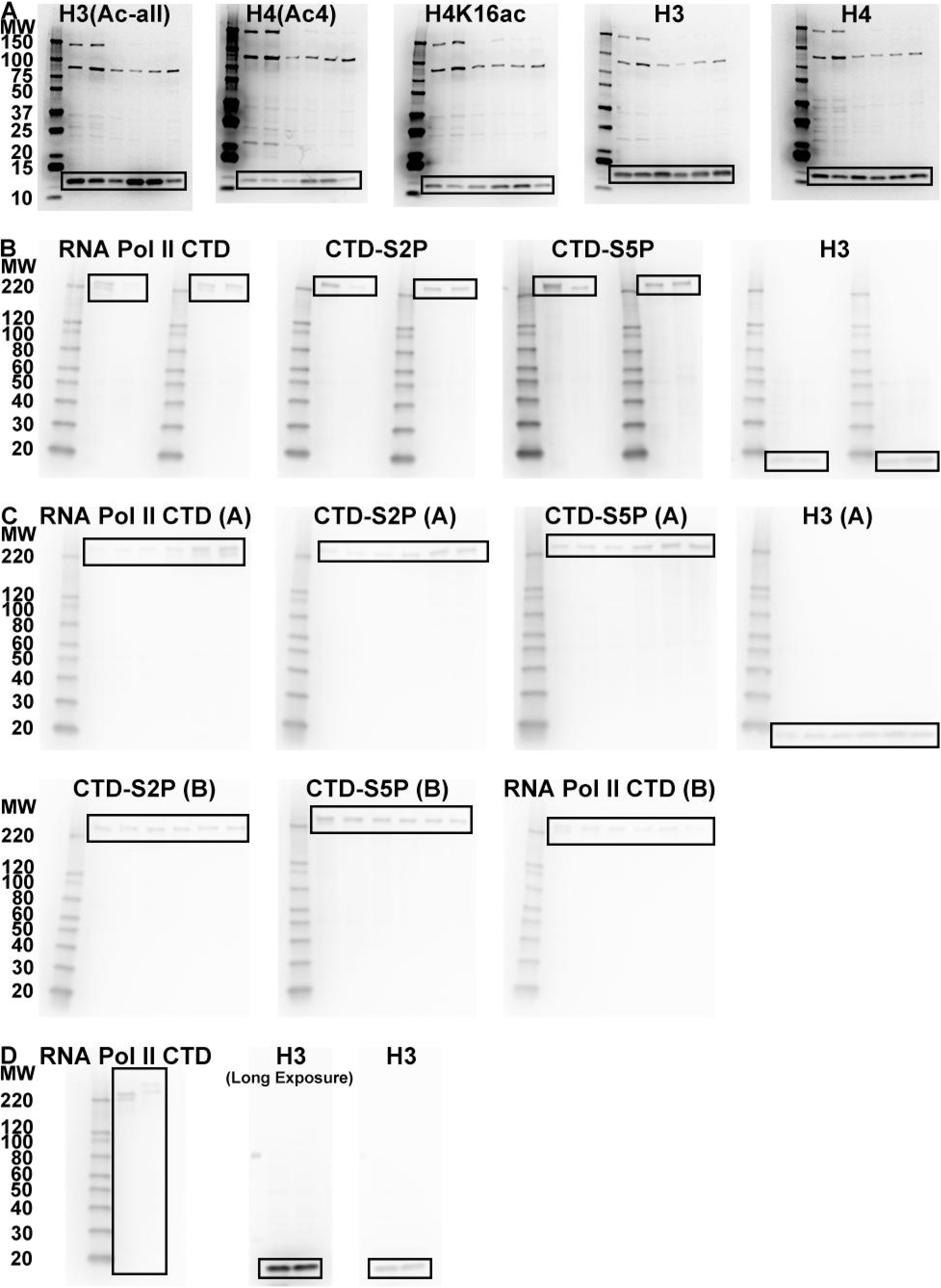
Uncropped immunoblots. For each immunoblot, the cropped bands are boxed at the bottom. (A) Immunoblots used in Figure 2A. The same molecular weight markers (MW, kilodaltons) were used for all 5 blots. Molecular weights are indicated to the left of the α-H3(Ac-all) blot. The detected high-molecular weight proteins (> 75 kilodaltons) are non-specifically bound by the StrepTactin-HRP conjugate. (B) Immunoblots used in Figures 3B and 3C. Molecular weight markers are different from those in part A, but the same across all 4 blots in this part. Molecular weights are indicated to the left of the α-CTD blot. For each antibody, the left set of lanes was loaded to have equal concentrations of H3, shown in Figure 3B, the right set of lanes was loaded to have equal concentrations of CTD, shown in Figure 3C. (C) Immunoblots used in Figure S4. Molecular weight markers used are the same as those used in part B. Molecular weights are indicated to the left of the α-CTD blot used in Figure S4A and the α-S2P blot used in Figure S4B. (D) Immunoblots used in Figure S1C. Molecular weight markers used for the first blot in this part are the same as those used in part B. Molecular weights are indicated to the left of the α-CTD blot. The GFP immunoblot and one of the H3 loading controls were exposed for 40 seconds during imaging. Both H3 loading controls do not have molecular weight markers. The anti-GFP blots shown in Figures S1C and S10C are already uncropped.

**Table S1.** Strains used in this paper.

| Strain | Parent | Genotype | Source |
| --- | --- | --- | --- |
| MBY99 | -- | 972 *h*- | MB lab |
| MBY165 | – | *cdc10-129* | MB lab |
| yFS240 (tk-hENT1) | – | *h- leu1-32 ura4-D18 ade6-210 his7-366 pJL218 (his7 adh1:tk)*<br>*pFS181 (leu1 adh1:hENT1)* | (Sivakumar et<br>al., 2004) |
| LGSP0001 | MBY99 | 972 *h- rpb1-eGFP-kanMX* | This paper |
| LGSP0002 | MBY99 | 972 *h- nup97-eGFP-kanMX* | This paper |

MB lab = gift from Mohan Balasubramanian lab

**Table S2.**
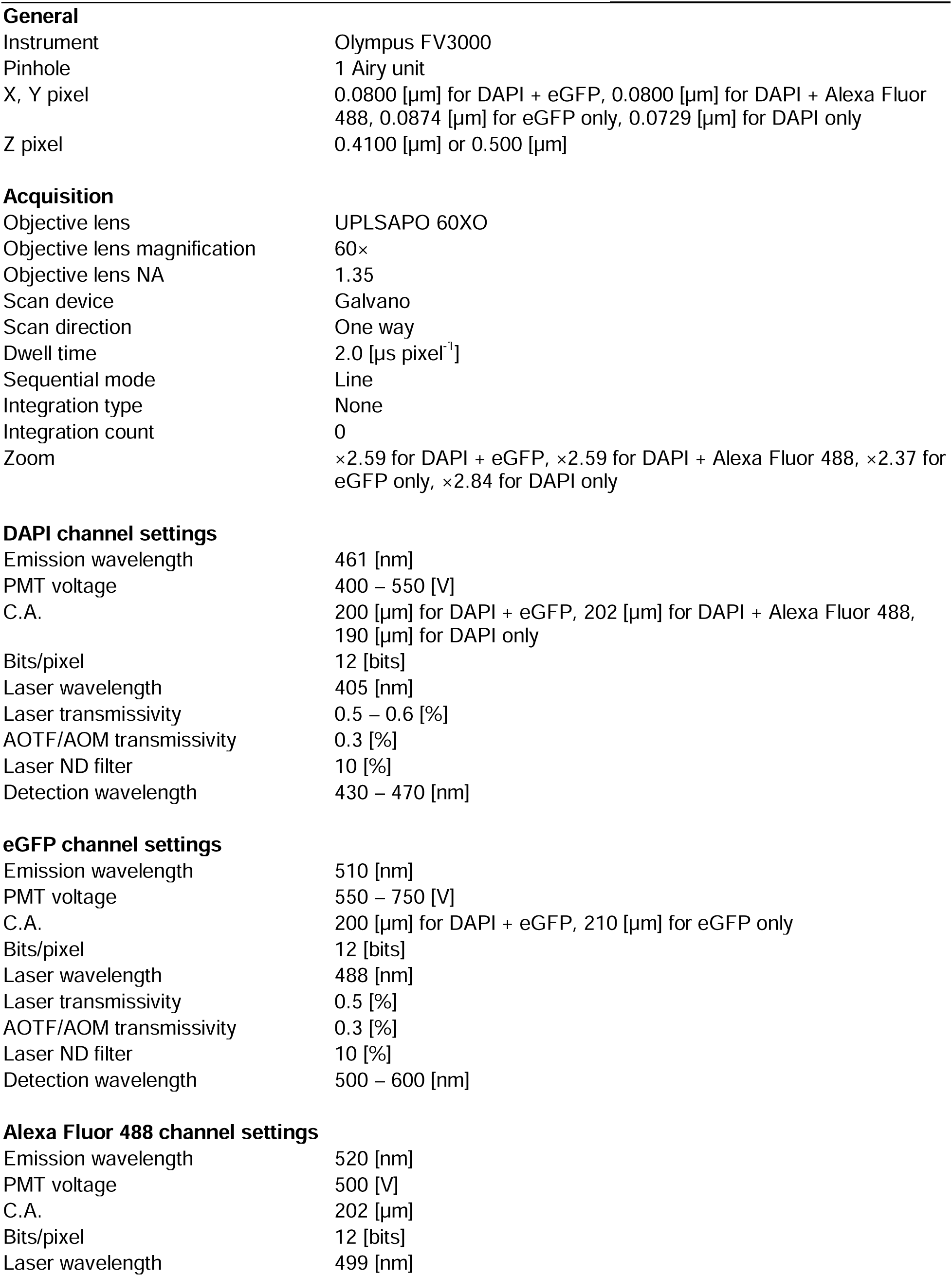
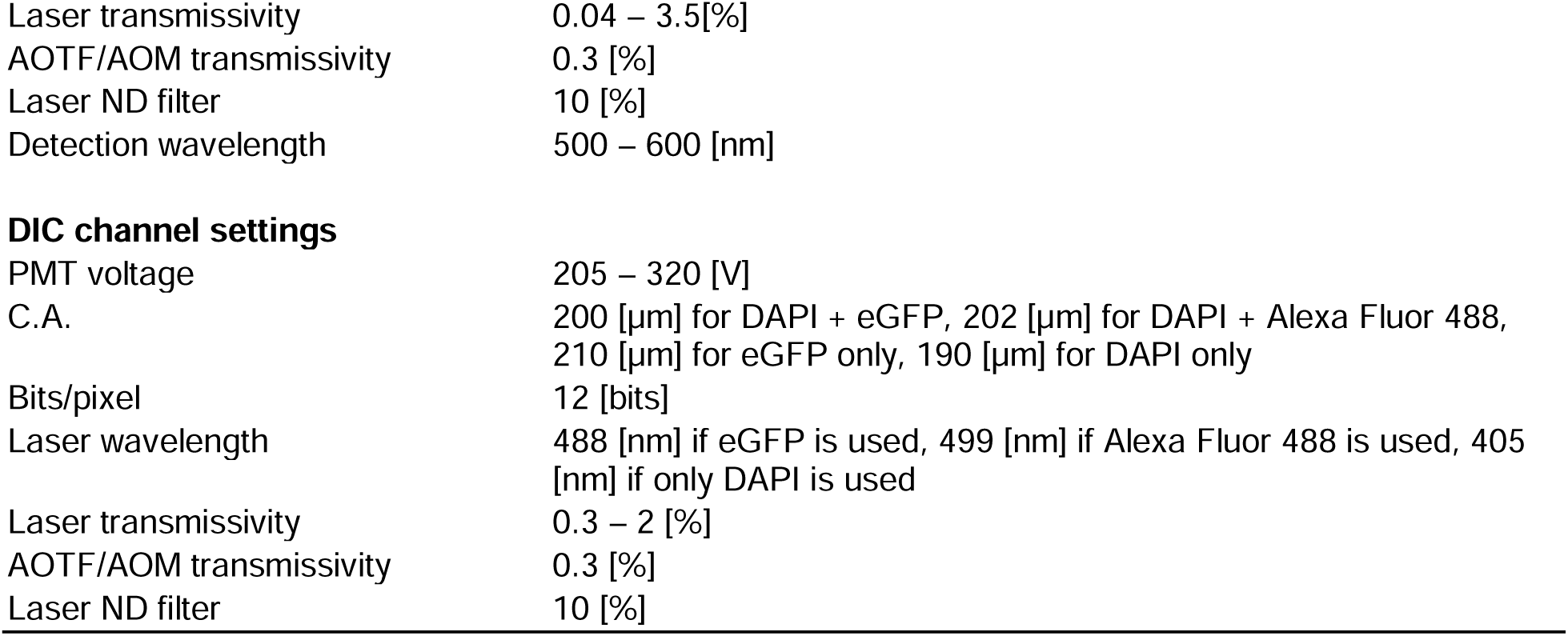
Confocal microscopy details.

|  |  |
| --- | --- |
| **General** |  |
| Instrument | Olympus FV3000 |
| Pinhole | 1 Airy unit |
| X, Y pixel | 0.0800 [μm] for DAPI + eGFP, 0.0800 [μm] for DAPI + Alexa Fluor 488, 0.0874 [μm] for eGFP only, 0.0729 [μm] for DAPI only |
| Z pixel | 0.4100 [μm] or 0.500 [μm] |
| **Acquisition** |  |
| Objective lens | UPLSAPO 60XO |
| Objective lens magnification | 60× |
| Objective lens NA | 1.35 |
| Scan device | Galvano |
| Scan direction | One way |
| Dwell time | 2.0 [μs pixel<sup>-1</sup>] |
| Sequential mode | Line |
| Integration type | None |
| Integration count | 0 |
| Zoom | ×2.59 for DAPI + eGFP, ×2.59 for DAPI + Alexa Fluor 488, ×2.37 for eGFP only, ×2.84 for DAPI only |
| **DAPI channel settings** |  |
| Emission wavelength | 461 [nm] |
| PMT voltage | 400 – 550 [V] |
| C.A. | 200 [μm] for DAPI + eGFP, 202 [μm] for DAPI + Alexa Fluor 488, 190 [μm] for DAPI only |
| Bits/pixel | 12 [bits] |
| Laser wavelength | 405 [nm] |
| Laser transmissivity | 0.5 – 0.6 [%] |
| AOTF/AOM transmissivity | 0.3 [%] |
| Laser ND filter | 10 [%] |
| Detection wavelength | 430 – 470 [nm] |
| **eGFP channel settings** |  |
| Emission wavelength | 510 [nm] |
| PMT voltage | 550 – 750 [V] |
| C.A. | 200 [μm] for DAPI + eGFP, 210 [μm] for eGFP only |
| Bits/pixel | 12 [bits] |
| Laser wavelength | 488 [nm] |
| Laser transmissivity | 0.5 [%] |
| AOTF/AOM transmissivity | 0.3 [%] |
| Laser ND filter | 10 [%] |
| Detection wavelength | 500 – 600 [nm] |
| **Alexa Fluor 488 channel settings** |  |
| Emission wavelength | 520 [nm] |
| PMT voltage | 500 [V] |
| C.A. | 202 [μm] |
| Bits/pixel | 12 [bits] |
| Laser wavelength | 499 [nm] |
| Laser transmissivity | 0.04 – 3.5[%] |
| AOTF/AOM transmissivity | 0.3 [%] |
| Laser ND filter | 10 [%] |
| Detection wavelength | 500 – 600 [nm] |
| **DIC channel settings** |  |
| PMT voltage | 205 – 320 [V] |
| C.A. | 200 [μm] for DAPI + eGFP, 202 [μm] for DAPI + Alexa Fluor 488,<br>210 [μm] for eGFP only, 190 [μm] for DAPI only |
| Bits/pixel | 12 [bits] |
| Laser wavelength | 488 [nm] if eGFP is used, 499 [nm] if Alexa Fluor 488 is used, 405<br>[nm] if only DAPI is used |
| Laser transmissivity | 0.3 – 2 [%] |
| AOTF/AOM transmissivity | 0.3 [%] |
| Laser ND filter | 10 [%] |

**Table S3.**
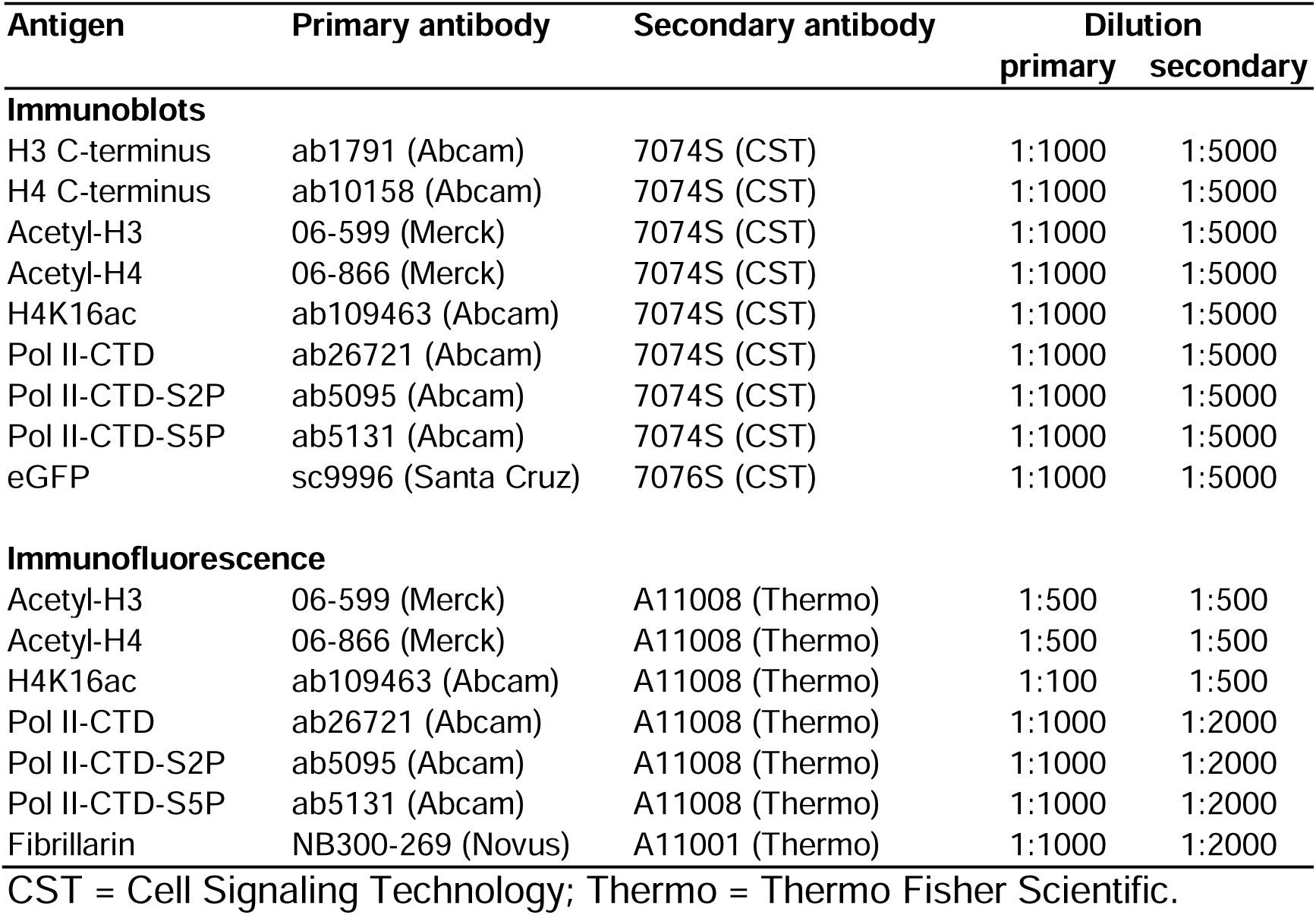
Antibodies used.

**Table S4.**
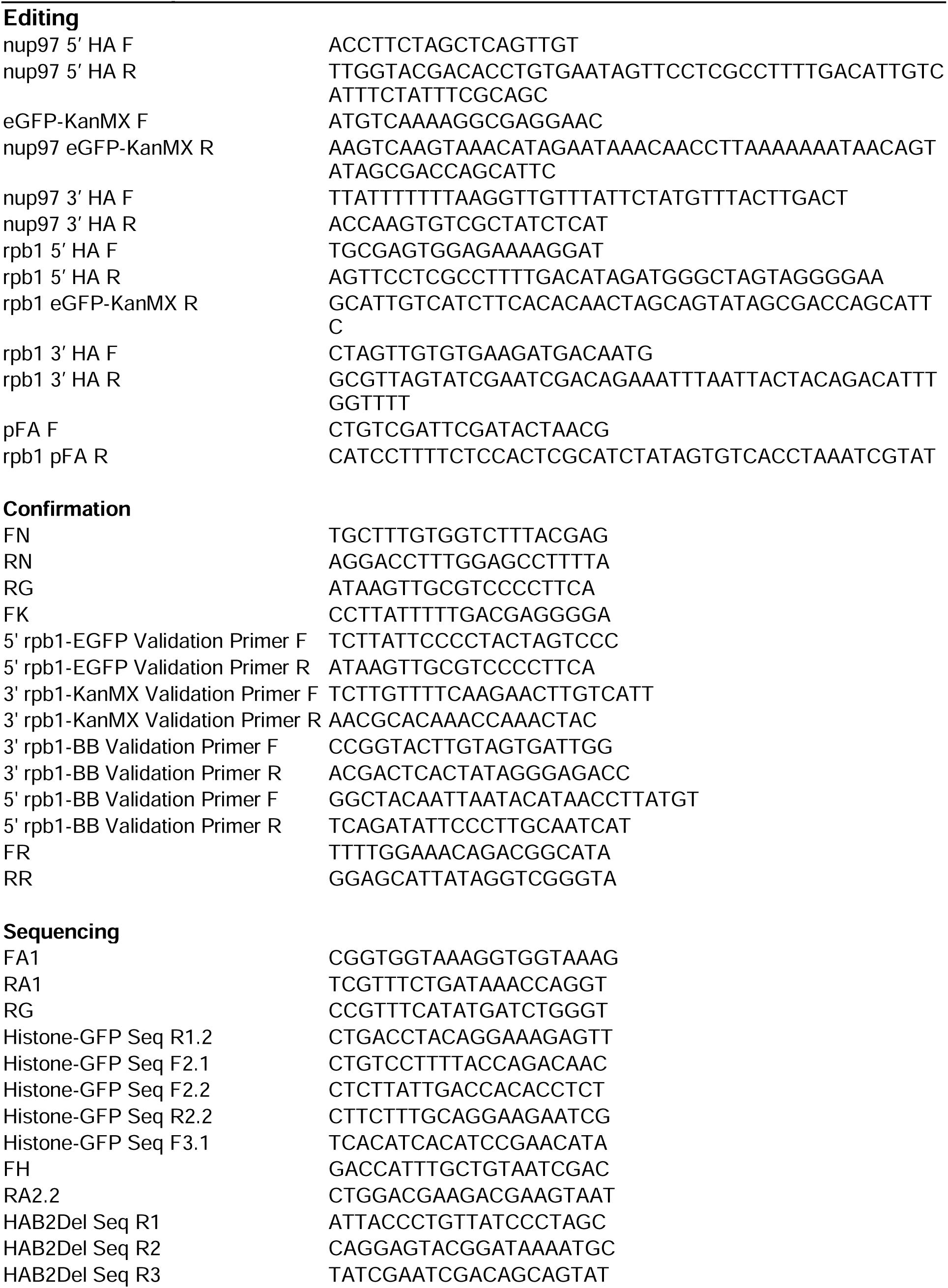
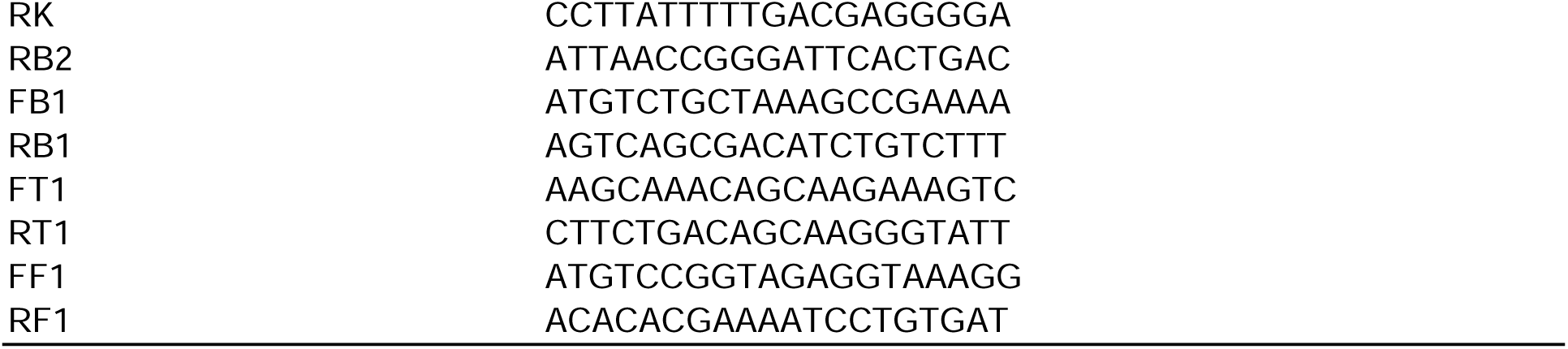
PCR primers, 5’ → 3’.

**Table S5.**
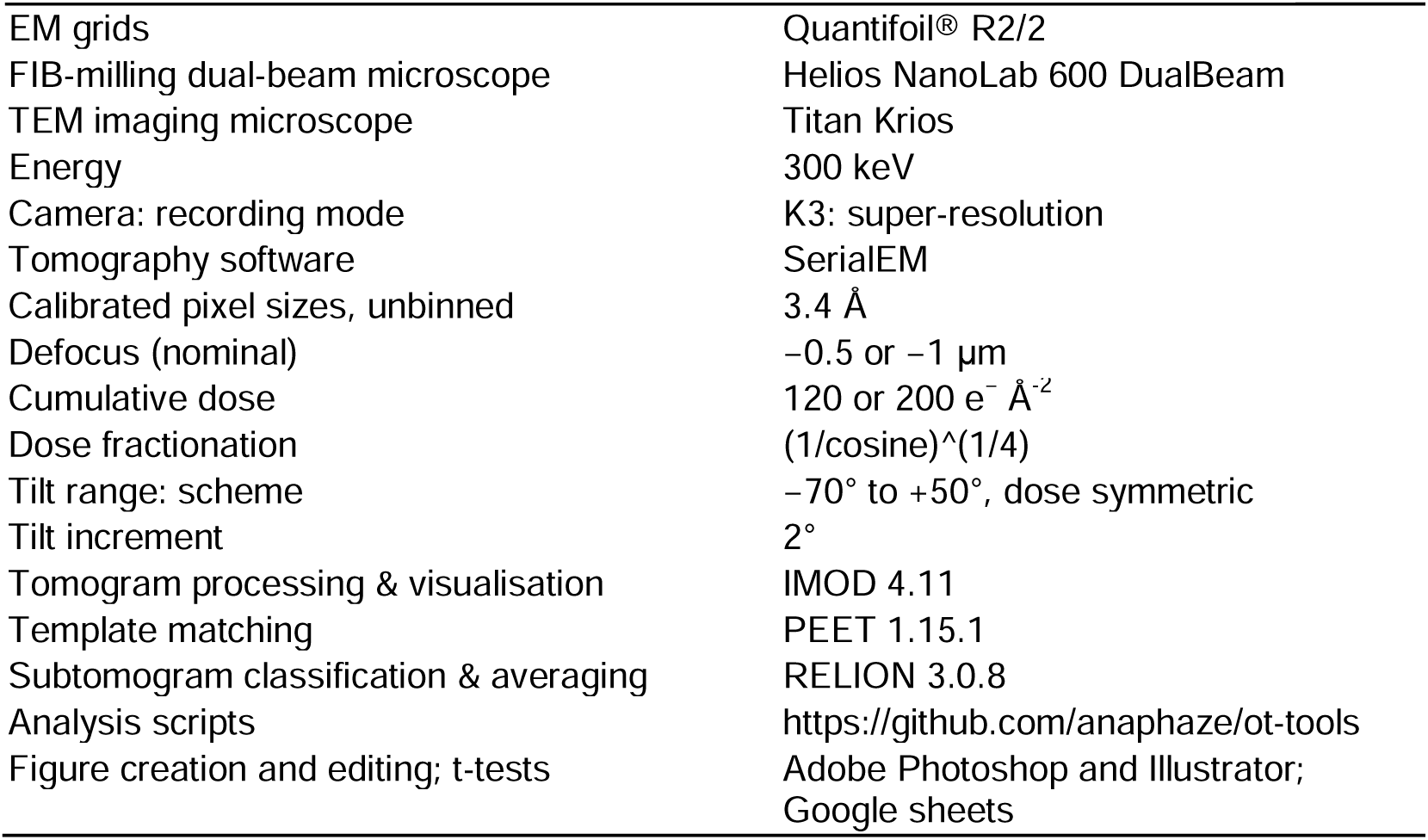
Lamella Cryo-EM imaging and analysis details.

**Table S6.**
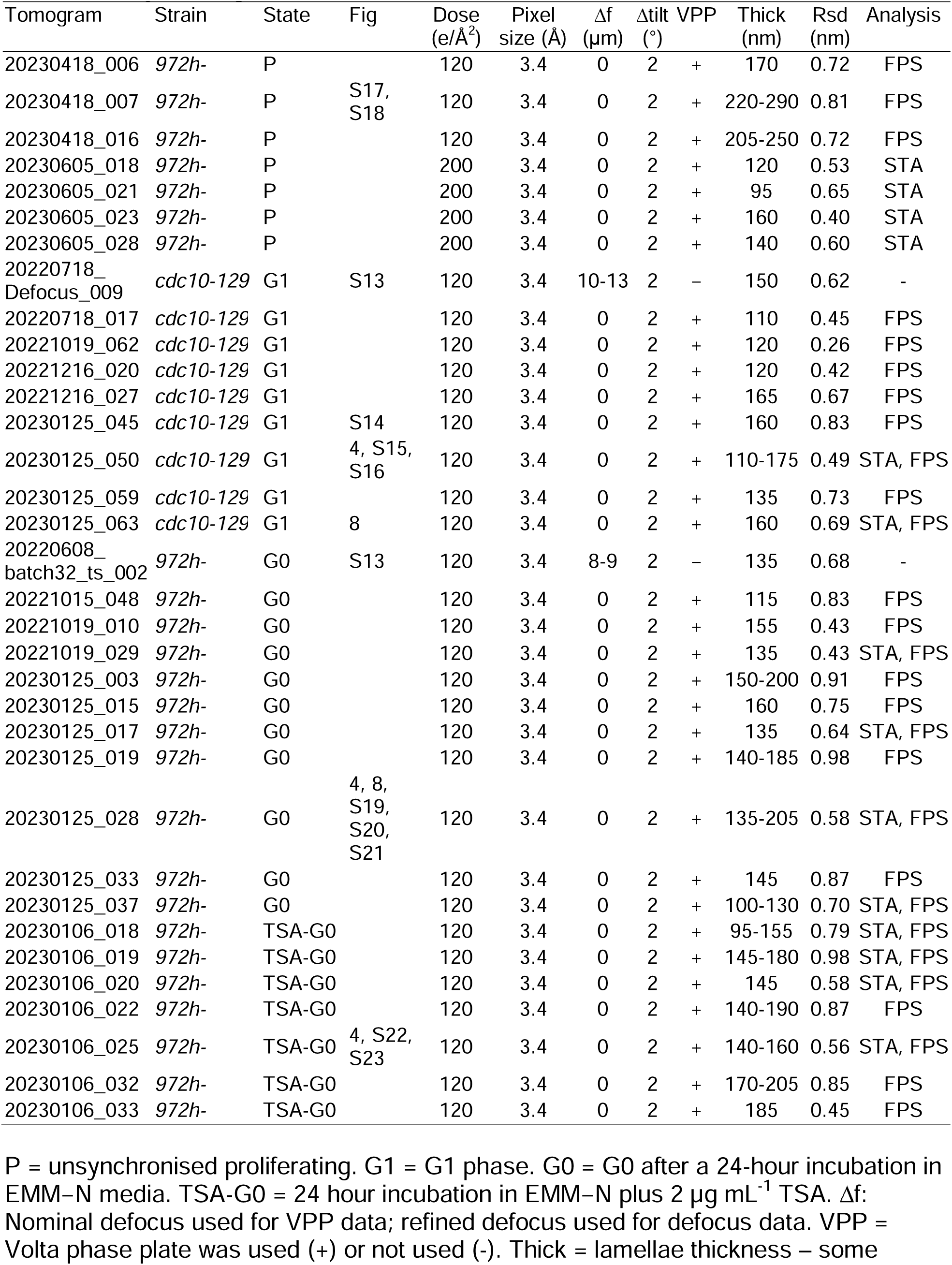
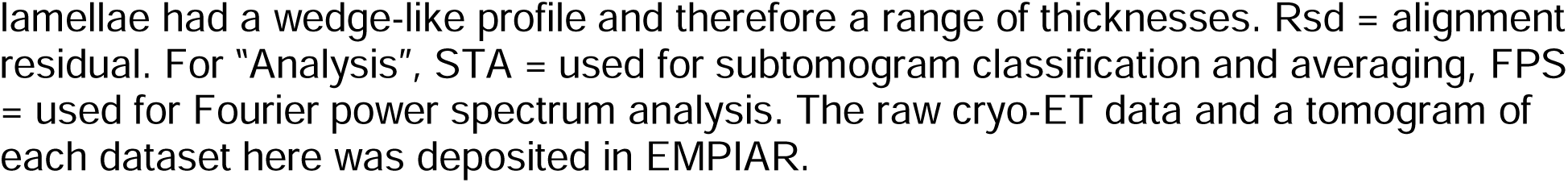
Cryotomogram details.

